# Hypothalamic plasticity for alloparenting in male mice

**DOI:** 10.64898/2026.10.21.760822

**Authors:** Patrick T. O’Neill, Rongzhen Yan, Yifan Wang, Qian Wang, Luping Yin, Gaybchoe Lhamo, Dylan J. Rhodes, Justin Little, Heping Cheng, Liangyi Chen, Shy Shoham, Xiaoke Chen, Robert C. Froemke, Dayu Lin

**Affiliations:** Neuroscience Institute, New York University Langone Medical Center; New York, NY, USA; Department of Biology, Stanford University; Stanford, CA, USA; Westlake Laboratory of Life Sciences and Biomedicine, School of Life Sciences, Westlake University, Institute of Biology, Westlake Institute for Advanced Study; Hangzhou, 310024, China; Tech4Health Institute, New York University Langone Medical Center; New York, NY, USA; National Biomedical Imaging Center, State Key Laboratory of Membrane Biology, Institute of Molecular Medicine, Center for Life Sciences, School of Future Technology, Peking University, Beijing, China; Department of Otolaryngology, New York University Langone Medical Center; New York, NY, USA; Department of Neuroscience and Physiology, New York University Langone Medical Center; New York, NY, USA; Center for Neural Science, New York University; New York, NY, USA; Department of Psychiatry, New York University Langone Medical Center; New York, NY, USA

## Abstract

Repeated exposure to young can induce caregiving in virgin female mice, a phenomenon known as pup sensitization, but whether virgin males can similarly acquire parental behavior and the neural mechanisms underlying this transition remain unclear. We show that 7 days of continuous pup exposure reliably converts virgin adult male mice to a parental state. This transition requires calcitonin receptor (CalcR)-expressing neurons in the medial preoptic area (MPOA): silencing MPOA^CalcR^ neurons disrupts both the acquisition and expression of paternal behavior, whereas optogenetic activation of GABAergic MPOA^CalcR^ neurons promotes caregiving in virgin males. Longitudinal two-photon calcium imaging reveals that MPOA^CalcR^ neurons develop enhanced pup-selective responses as males transition to a parental state. Moreover, CalcR signaling in the MPOA promotes synaptic potentiation and facilitates the emergence of parental behavior. These results identify an experience-dependent hypothalamic plasticity mechanism that enables parental behavior in non-father males.

## Main Text

Mammalian parents have evolved a set of robust and stereotyped behaviors to ensure offspring survival, including nursing, crouching, grooming, and young retrieving. Additionally, nulliparous female mice can display pup-caring behaviors after interacting with unrelated young^1–6^. This process, termed pup sensitization, occurs independently of sex hormones^5,7^.

Adult unmated non-father (referred to as virgin) male mice are typically aggressive or indifferent to pups, whereas fathers, like mothers, show robust parental behaviors. This behavioral switch during fatherhood appears to require ejaculation and is facilitated by co-habituation with females^8–11^. Unlike females^1^, however, repeated, short-term exposure to pups (15 minutes per day) was unable to induce parental behaviors in adult virgin male mice^12^. Consequently, virgin males have been considered incapable of alloparenting, i.e., caring for pups that they did not sire, and are rarely studied in the field. However, exposure to pups during development suppresses infanticide in adult male mice^13^ and cohabitation with a dam and litter can promote paternal behaviors in virgin male mice^14^. A recent study further demonstrated that daily one hour exposure to pups for three days can suppress infanticide and facilitate parental behaviors in some isolated male mice^15^. These studies collectively indicate that sexual experience may not be strictly required to affect pup-directed behaviors and promote alloparental care in non-fathering males. Thus, the first goal of our study is to address whether virgin male mice can show reliable alloparental care, as observed in humans^16,17^.

Numerous studies support the central role of the medial preoptic area (MPOA) in mediating parental behaviors^18–20^. Classic lesion or deafferentation studies demonstrated a total loss of maternal care after MPOA damage in mice and rats^18,21–29^. Newer work has focused on examining specific cell types in the MPOA, and identified cells expressing galanin (Gal), estrogen receptor alpha (Esr1), or the prolactin receptor (Prlr) as important for the regulation of parental behavior^19,30–36^. More recently, systematic snRNAseq and multiplexed error robust in situ hybridization (MERFISH) revealed that calcitonin receptor (CalcR) expressing MPOA (MPOA^CalcR^) cells are preferentially active (based on immediate early gene expression) during parental behaviors in virgin females, dams, and sires^35^. Later, silencing MPOA^CalcR^ or knocking down CalcR expression in the MPOA was shown to suppress parental behaviors in female mice^37^. MPOA^CalcR^ neurons are only a small subset of MPOA Gal, Esr1, or Prlr expressing cells^35,37^ and thus represent the most selective population for maternal behaviors known to date. The functional role of MPOA^CalcR^ neurons in male parental behaviors remains unclear.

In this study, we examined alloparenting behaviors in virgin male mice and the potential contribution of MPOA^CalcR^ cells and CalcR signaling in the emergence and expression of these behaviors.

## Results

### Induction of alloparenting in sexually naïve male mice

We first asked whether extensive pup interaction can induce the emergence of parental behaviors in virgin male mice (**Fig. 1a**). Because C57/BL6 male mice are often infanticidal prior to mating ^10,38^, we developed a pup sensitization protocol, involving placing 2-3 pups in a wire mesh ball and co-housing them with males over the course of one week. As a control, we placed a novel inanimate object in the mesh ball instead of pups (**Fig. 1a**). Before the first day of pup sensitization, we tested male behavior toward pups in a pup interaction test by introducing 10 pups, one at a time for 2 min, into the male’s home cage (**Fig. 1a**). On each subsequent day, we removed the pup-containing mesh ball and repeated the pup interaction test using unfamiliar pups (**Fig. 1a**). Over 7 days of pup sensitization, infanticidal behavior stopped in all male mice (days to last infanticidal event, mean ± SEM: 3.0 ± 0.7 days) (**Fig. 1b, c, e**), and 19/20 male mice retrieved pups (days to first retrieval, mean ± SEM: 3.5 ± 0.5 days) (**Fig. 1 b, d, f, g**). In contrast, animals co-housed with objects continued to be infanticidal and never retrieved pups (**Fig. 1b-g**). While both pup- and object-sensitized animals contacted pups more quickly across days of sensitization, those that underwent pup sensitization spent significantly more time investigating, grooming, and crouching over the pups, as well as building nests (**Fig. 1h-l**). The sensitization-induced parental behaviors did not depend on sex hormones; castrated males began to show parental behaviors on a similar time course to intact males, although castration abolished infanticide as previously reported^39^ (**Extended Data Fig. 1**).

**Fig. 1.**
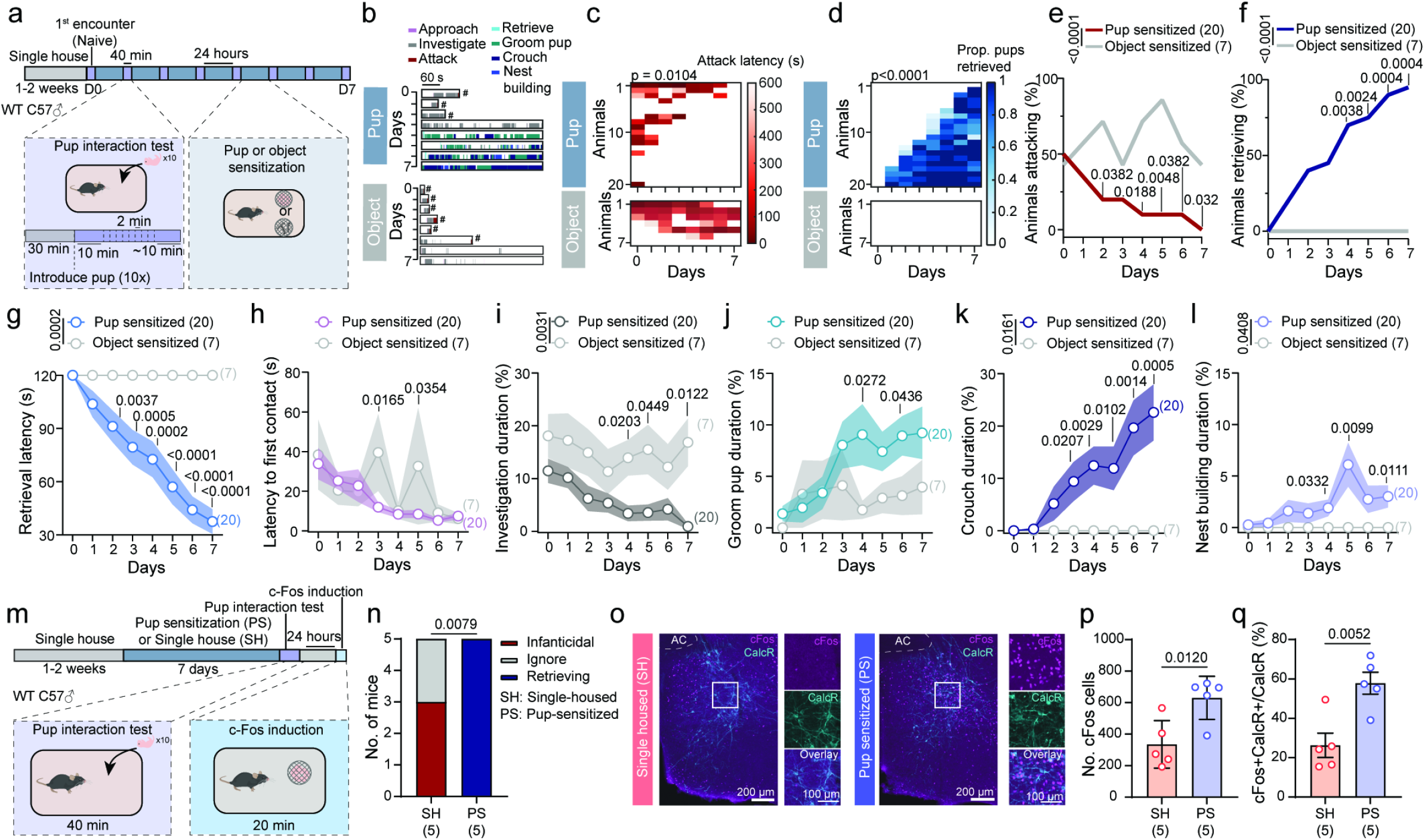
Pup sensitization promotes parental behavior in male mice. **(a)** Pup sensitization and novel object sensitization paradigm. **(b)** Representative raster plots showing pup-directed behaviors in the pup interaction test in pup- and object-sensitized groups. Raster plots represent the 10 minutes after introducing the last pup during the pup interaction test. (# indicates removal of the pup after being attacked) **(c, d)** Quantification of the latency to attack **(c)** and proportion of pups retrieved **(d)** during the pup interaction test across 7 days of pup and object sensitization. Day 0: before pup or object sensitization; days 1-7: during pup or object sensitization. If no attack occurred, the latency was scored as 600 s. **(e, f)** Percentage of animals attacking **(e)** or retrieving pups **(f)** during the pup interaction test across 7 days of pup or object sensitization. **(g-l)** The latency to retrieve pups **(g)**, latency to contact pups **(h)**, pup investigation duration **(i)**, grooming pup duration **(j)**, crouching over pups duration **(k)**, and nest building duration **(l)** during the pup interaction test across 7 days of pup or object sensitization. Day 0: before pup sensitization, days 1-7: during pup sensitization. **(m)** Paradigm to induce c-Fos following pup sensitization or single housing. **(n)** The number of mice that were infanticidal, non-infanticidal, and non-parental (ignore), or retrieving following single housing (SH) or pup sensitization (PS). **(o)** Representative histology images showing c-Fos (magenta) and calcitonin receptor (CalcR; cyan) immunostaining in a single-housed (SH) and a pup sensitized (PS) mouse following 20-min pup interaction. (AC = anterior commissure) **(p)** The number of c-Fos-expressing cells in the MPOA in single-housed (SH) and pup sensitized (PS) mice. **(q)** The number of double-positive c-Fos and CalcR co-expressing cells normalized by the number of CalcR-expressing cells in the MPOA in single-housed (SH) and pup sensitized (PS) mice. Mouse cartoons in this and all other figures are adopted from scidraw.io. Numbers in parentheses or axis titles indicate the subject animal number. Circles in **p, q** represent data of individual animals. **(c, d, g-l)** Two-way ANOVA with repeated measures followed by Tukey’s multiple comparisons test. **(e, f)** Log-rank test across groups and Fisher’s exact test followed by Benjamini Hochberg false-discovery rate correction for each day. **(n)** Fisher’s exact test. **(p, q)** Unpaired t-test. Lines and shaded error bars represent mean ± SEM. Bars and error bars represent mean ± SEM. All p < 0.05 are indicated.

We then asked whether MPOA^CalcR^ cells in males were activated during pup interaction, and if so, whether the activation level depends on the parental state of the males. We introduced 3 pups housed in a wire mesh ball to the home cage of the test male mouse for 20 minutes after either 7 days of pup sensitization or single housing and examined c-Fos expression (**Fig. 1m**). All 5 pup-sensitized animals showed reliable parental behaviors, while none of the single-housed males did (**Fig. 1n**). We found that c-Fos expression was nearly twice as abundant in the MPOA of pup-sensitized animals compared to single-housed animals, and a larger fraction of MPOA^CalcR^ cells expressed c-Fos after pup exposure in sensitized males than single-housed males (**Fig. 1o-q**). These results suggest that extensive pup exposure can induce the switch from negative to positive pup-directed behaviors in sexually naïve males independent of sex hormones, and this behavioral change is accompanied by increased responses of MPOA^CalcR^ cells to pups.

### Inhibiting MPOA^CalcR^ neurons impairs alloparenting behaviors

We next asked if MPOA^CalcR^ cells are required for alloparenting behaviors in virgin males. We bilaterally injected Cre-dependent hM4Di or mCherry virus into the MPOA of virgin Calcr-2A-Cre (Calcr^Cr*e*^) male mice ^40^ (**Fig. 2a**). Three weeks later, we sensitized these animals until they showed a high level of paternal behavior. On alternating days, we injected saline or CNO (1mg/kg) intraperitoneally and, 30 minutes later, introduced pups (one by one for 10 trials) into the male home cages (**Fig. 2b**). Following CNO injection, hM4Di animals, but not mCherry animals, became significantly less likely to retrieve pups, and some animals attacked pups (**Fig. 2c-f**). Additionally, the pup grooming duration significantly decreased, and the pup crouching duration also trended downwards (**Fig. 2i, j**), whereas the latency to contact pups, investigation duration, and nesting building time did not change significantly after CNO injection (**Fig. 2g, h, k**). Inhibiting these neurons neither affected inter-male aggression (**Extended Data Fig. 2a-d**) nor sexual behaviors toward adult females (**Extended Data Fig. 2e-h**). Together, these results indicate that MPOA^CalcR^ cells are necessary for expressing paternal behaviors in virgin males and seemingly dispensable for adult-directed social behaviors.

**Fig. 2.**
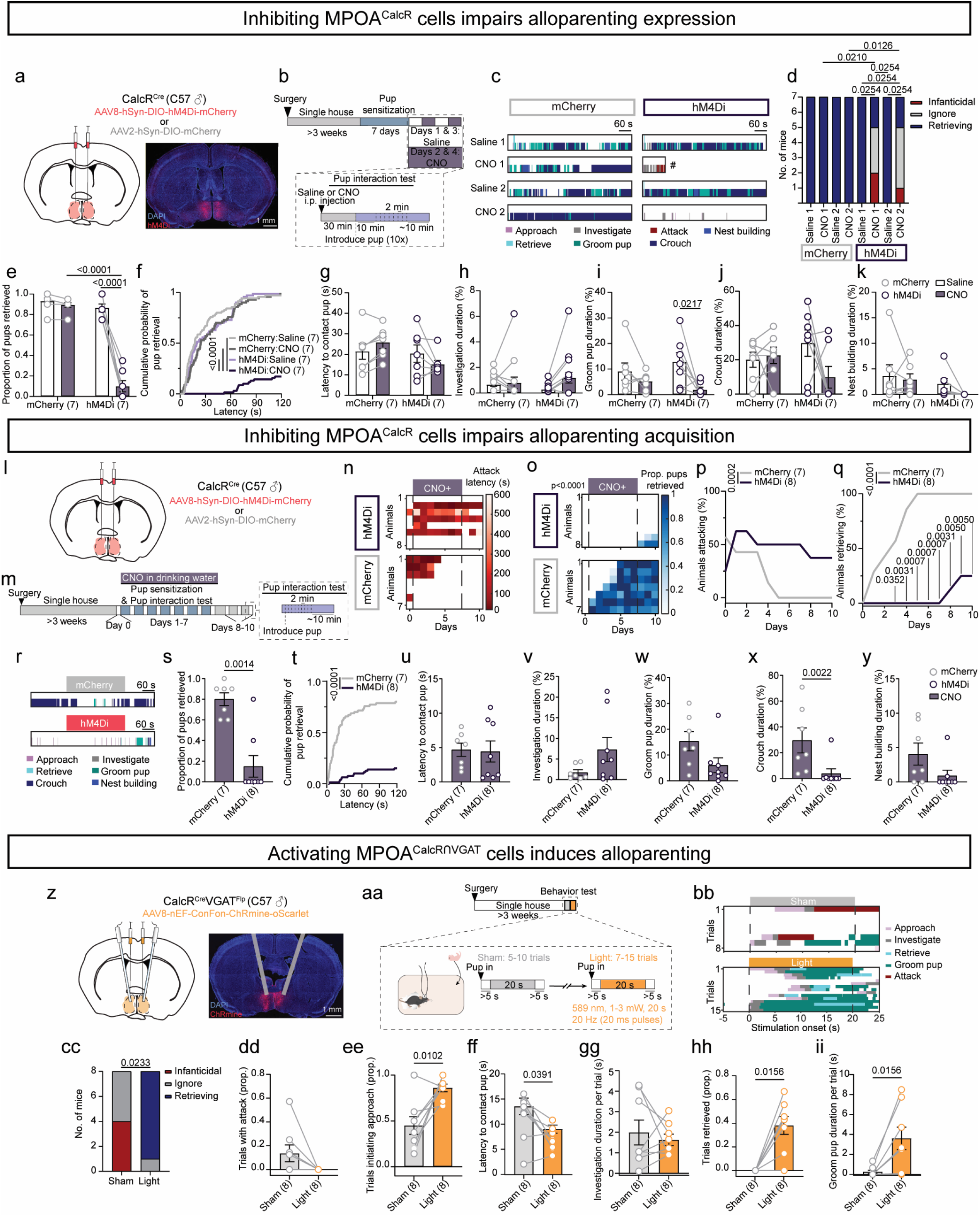
MPOA^CalcR^ cells are necessary and MPOA^CalcRՈVgat^ cells are sufficient for paternal behavior. **(a)** Virus injection schematic and representative histology image. **(b)** Timeline to chemogenetically inhibit MPOA^CalcR^ cells following pup sensitization. **(c)** Representative raster plots showing pup-directed behaviors in the pup interaction test following saline or CNO injection in an example mCherry- and hM4Di-injected animal. Raster plots represent the 10 minutes following the introduction of the last pup. (# indicates removal of the pup after being attacked) **(d)** The number of mice that were infanticidal, non-infanticidal, and non-parental (ignore), or retrieving after saline or CNO injection in mCherry and hM4Di mice. **(e-k)** The proportion of pups retrieved **(e)**, cumulative probability of pup retrieval over the 120 s testing period **(f)**, latency to contact pups **(g)**, pup investigation duration **(h)**, grooming pup duration **(i)**, crouching over pups duration **(j)**, and nest building duration **(k)** in mCherry and hM4Di mice after saline or CNO injection. Saline and CNO days are averaged within mCherry and hM4Di groups. **(l)** Virus injection schematic. **(m)** Timeline to chemogenetically inhibit MPOA^CalcR^ cells during pup sensitization. **(n)** The latency to attack pups across the duration of the experiment (day 0: before pup sensitization, days 1-7: during pup sensitization, days 8-10: after pup sensitization). **(o)** The proportion of pups retrieved across the duration of the experiment (day 0: before pup sensitization, days 1-7: during pup sensitization, days 8-10: after pup sensitization). **(p)** Percentage of animals attacking pups in the pup interaction test across the duration of the experiment (day 0: before pup sensitization, days 1-7: during pup sensitization, days 8-10: after pup sensitization). **(q)** Percentage of animals retrieving pups in the pup interaction test across the duration of the experiment (day 0: before pup sensitization, days 1-7: during pup sensitization, days 8-10: after pup sensitization). **(r)** Representative raster plots showing pup-directed behaviors in the pup interaction test on day 10 of the experiment in an mCherry- and an hM4Di-injected animal. Raster plots represent the 10 minutes following the introduction of the last pup. **(s-y)** The proportion of pups retrieved **(s)**, cumulative probability of pup retrieval over the 120 s testing period **(t)**, latency to contact pups **(u)**, pup investigation duration **(v)**, grooming pup duration **(w)**, crouching over pups duration **(x)**, and nest building duration **(y)** in mCherry and hM4Di mice during the pup interaction test on day 10 of the experiment. **(z)** Virus injection and optical fiber implantation schematic and representative histology image. **(aa)** Timeline to optogenetically activate MPOA^CalcR∩VGAT^ cells. **(bb)** Representative raster plots aligned to the onset of sham or light stimulation in a ChRmine-injected animal. **(cc)** The number of mice that were infanticidal, non-infanticidal and non-retrieving (ignore), and retrieving following sham or light stimulation. **(dd)** Proportion of sham or light trials with pup-directed attack. **(ee)** Proportion of sham or light trials with pup approach. **(ff)** The latency to contact pups following light or sham onset. **(gg)** The time spent investigating pups during sham or light stimulation. **(hh)** Proportion of sham or light trials with pup retrieval. **(ii)** The time spent grooming pups during sham or light stimulation. The brain illustrations in this and all other figures were produced based on a reference atlas from https://atlas.brain-map.org/. Numbers in parentheses or axis titles indicate the subject animal number. Circles and circle-line pairs represent data of individual animals. **(d)** Cochran’s Q test followed by Benjamini Hochberg false-discovery rate correction within hM4Di group, Fisher’s exact test between mCherry and hM4Di groups. **(e, g-k)** Two-way ANOVA with repeated measures followed by Šidák’s multiple comparisons test. **(f, t)** Logrank (Mantel-Cox) test. **(n, o)** Two-way ANOVA with repeated measures followed by Tukey’s multiple comparisons test. **(p, q)** Log-rank test across groups, Fisher’s exact test followed by Benjamini Hochberg false-discovery rate correction on each day. **(s, x, y)** Mann-Whitney U test. **(u-w)** Unpaired t-test. **(cc)** McNemar’s test (Outcome 1: retrieval; Outcome 2: no retrieval). **(dd, ff-ii)** Wilcoxon test. **(ee)** Paired t-test. Bars and error bars represent mean ± SEM. All p < 0.05 are indicated.

To understand if MPOA^CalcR^ cells regulate the emergence of paternal behaviors, we again used a chemogenetic approach to inhibit these cells throughout the period of pup sensitization (**Fig. 2l, m**). Specifically, we continuously delivered CNO through the drinking water. During the 7 days of pup sensitization, hM4Di animals remained largely infanticidal and did not retrieve pups during the pup-interaction tests (**Fig. 2n-q**). Meanwhile, all mCherry animals ceased attacking pups by day 5 of pup sensitization and began to retrieve pups (**Fig. 2n-q**). Because MPOA^CalcR^ cells are necessary for expressing paternal behaviors, we reasoned that while the cells were tonically inhibited during the sensitization period, the animals may be unable to commit motor actions corresponding to parenting, even if they are parental. Thus, we tested the animals for 3 additional days after stopping CNO administration and pup sensitization (**Fig. 2m**) and found only 2 out of 8 hM4Di-injected animals retrieved pups (**Fig. 2o, q**). On the 3^rd^ day of testing after CNO termination, compared to mCherry-injected animals, hM4Di animals retrieved significantly fewer pups and spent less time crouching over pups (**Fig. 2r-t, x**). Other parental behaviors, such as pup grooming and nest building, also showed trends of lower performance in hM4Di animals, although hM4Di and mCherry animals contacted the pups with a similar latency (**Fig. 2u-w, y**). These results suggest that MPOA^CalcR^ cells are required for the emergence of paternal behavior with pup sensitization.

### Stimulation of MPOA^CalcR∩VGAT^ neurons promotes paternal behaviors

We next asked if we could accelerate the onset of paternal care in naïve male mice by activating MPOA^CalcR^ cells. We injected a Cre-dependent channelrhodopsin (ChR2) virus bilaterally into the MPOA of Calcr^Cre^ male mice and delivered sham or blue light (20 ms, 20 Hz, 20 s) while introducing pups one by one (**Extended Data Fig. 3a, b**). Although MPOA^CalcR^ stimulation promoted pup approach, as reflected by the shorter latency to contact pups in light trials compared to sham trials (0 mW), the stimulated animal did not show pup retrieval or increase any other parental behaviors (**Extended Data Fig. 3c-j**).

The MPOA^CalcR^ cells are approximately 80% GABAergic and 20% glutamatergic based on snRNAseq analysis^35,41^. Most (approximately 75%) of the MPOA neurons expressing c-Fos during maternal behaviors are GABAergic^25^, and optogenetic inhibition of glutamatergic cells in the MPOA increases, rather than decreases, pup retrieval in virgin females^42^. Our previous study showed that MPOA^Esr1^ cells send primarily GABAergic inputs to the ventral tegmental area (VTA) to disinhibit dopamine cells and promote pup retrieval^30^. Thus, we hypothesized that the GABAergic subpopulation of MPOA^CalcR^ cells may be the most relevant subset of MPOA^CalcR^ cells for parenting. To this end, we stimulated MPOA^CalcR∩VGAT^ cells by injecting the Cre- and Flp-dependent opsin ChRmine^43^ into the MPOA of Calcr^Cre^Vgat^Flp^ *(Slc32a1*-2A-*Flp*^44^) male mice and delivered sham or yellow light (20 ms, 20 Hz, 20 s) after pup introduction (**Fig. 2z, aa**). Light stimulation ceased infanticide (**Fig. 2bb-dd**), robustly induced approach toward the pup, and shortened the latency to pup contact without altering pup investigation duration (**Fig. 2ee-gg**). Strikingly, 7/8 males repeatedly retrieved and groomed pups, signifying a rapid switch to paternal behavior (**Fig. 2hh, ii**).

To test if glutamatergic MPOA^CalcR^ cells also play a role in parenting, we used the same interactional strategy to target and stimulate the MPOA^CalcR∩VGlut2^ population in Calcr^Cre^Vglut2^Flp^ *(Slc17a6-*IRES2-*Flp*^44^) male mice (**Extended Data Fig. 4a, b**). Stimulation caused no change in pup-directed behaviors (**Extended Data Fig. 4c-j**). These results suggest that MPOA^CalcR∩VGlut2^ population is likely not involved in paternal behaviors and underscore the importance of MPOA^CalcR∩VGAT^ cells in parenting.

To understand the functional specificity of MPOA^CalcR∩VGAT^ cells, we tested their relevance in orchestrating behaviors toward other social targets. We found that MPOA^CalcR∩VGAT^ stimulation did not affect interactions with adult males (**Extended Data Fig. 5a-d**) or female mice (**Extended Data Fig. 5e-h**) but did induce a place preference in a real-time place preference assay, suggesting activating the cells is of positive valence (**Extended Data Fig. 5i, j**). Stimulation of MPOA^CalcR^ or MPOA^CalcR∩VGlut2^ cells also had no effect on male- or female-directed behaviors (**Extended Data Fig. 3k-r** and **Extended Data Fig. 4k-r**) and did not induce a place preference (**Extended Data Fig. 3s, t** and **Extended Data Fig. 4s, t**).

Anterograde tracing revealed that MPOA^CalcR∩VGAT^ cells send projections to regions known to orchestrate parental behaviors, including the ventral tegmental area (VTA)^30,31^, paraventricular nucleus of the hypothalamus (PVN)^3^, and anteroventral periventricular nucleus (AVPV)^45^ (**Extended Data Fig. 6**). Additionally, MPOA^CalcR∩VGAT^ cells also target infanticide-promoting regions, including the rhomboid^29^ and principal nuclei^46^ of the bed nucleus of the stria terminalis (BNSTrh and BNSTpr) and the perifornical area (PeFA)^47^ (**Extended Data Fig. 6**). As such, MPOA^CalcR∩VGAT^ population is well positioned to counteract hostile behaviors toward the young and orchestrate parental behaviors.

### Recording MPOA^CalcR^ neural activity during parenting

As we identified MPOA^CalcR^ cells as a critical population mediating parenting, we next wanted to understand how these cells respond during parental behaviors. We injected Cre-dependent GCaMP6s^48^ virus into the MPOA of Calcr^Cr*e*^ male mice and recorded the calcium activity of individual neurons using a head-mounted two-photon microscope^49^ during paternal behaviors after pup sensitization (**Fig. 3a-c**). During the recording, we introduced 10 pups, one at a time, into the male’s home cage (**Fig. 3c**). A total of 213 cells were imaged from 6 mice (26-50 cells recorded from each animal) that showed robust parental behaviors. An example session is shown in **Fig. 3d**. Across all recorded cells, 71% increased activity significantly during at least one pup-directed behavior, with many responding during two or more behavioral events (**Fig. 3e**). Activity changes during pup approach, investigation, retrieval, and grooming are positively correlated with each other, whereas response during crouching is negatively correlated with those during approach, investigation, and retrieval (**Fig. 3f, Extended Data Fig. 7**), suggesting distinct MPOA^CalcR^ cell response patterns during active vs. passive pup-directed behaviors.

**Fig. 3.**
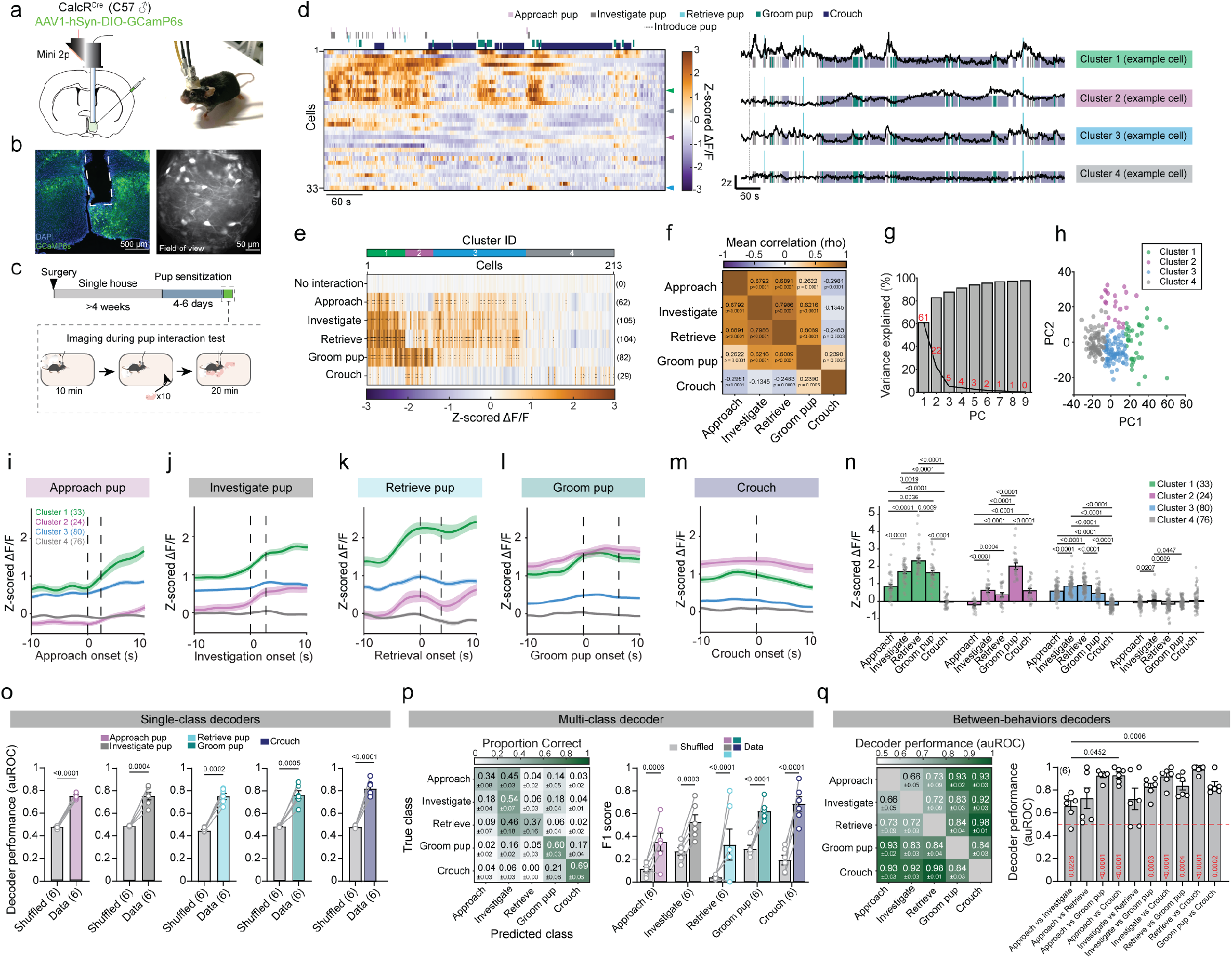
Activity of MPOA^CalcR^ cells during parenting. **(a)** Virus injection, GRIN lens implant strategy, and example mouse with a head-mounted 2-photon microscope (mini2p). **(b)** Histology showing GRIN lens track (white dashed line) and GCaMP6s expression (green) in the MPOA (left) and session-averaged imaging plane showing GCaMP6s-expressing cells (right). **(c)** Experimental timeline to image MPOA^CalcR^ cell calcium activity during parenting. **(d)** Left: Heatmap showing Z-scored ΔF/F of MPOA^CalcR^ cells during parenting (top) in an example recording session. Cells are arranged based on their membership in a k-means clustering analysis (n = 33 cells). Right: Representative traces of cells belonging to different clusters during parenting. Colored triangles next to the heatmap indicate representative cells to the right. **(e)** Heatmap showing average Z-scored ΔF/F MPOA^CalcR^ cells during pup-directed behaviors, arranged based on their membership in a k-means clustering analysis (n = 213 cells from 6 mice). Green, blue, magenta, and gray bars correspond to cluster identities (green: 1^st^ cluster = 33 cells, magenta: 2^nd^ cluster = 24 cells, magenta: 3^rd^ cluster = 80 cells, gray: 4^th^ cluster = 76 cells). Numbers on the right indicate the number of significantly responsive cells. Black circles indicate q values. ●q<0.01; ●●q<0.001. **(f)** Correlation matrix of the average Z-scored ΔF/F MPOA^CalcR^ cells across parental behaviors. **(g)** Scree plot showing the variance explained by the principal components of the population activity matrix. Red numbers indicate the variance explained by each component. **(h)** Scatterplot of cells belonging to each cluster against the first 2 principal components. **(i-m)** Average peri-event time histograms (PETHs) of Z-scored ΔF/F of the four MPOA^CalcR^ cell clusters (as shown in **e** and **h**) aligned to the onset of approaching **(i)**, investigating **(j)**, retrieving **(k)**, grooming **(l)**, and crouching over **(m)** pups. Dashed lines indicate the onset and average offset of behavior. **(n)** Average responses of the four MPOA^CalcR^ cell clusters across pup-directed behaviors. **(o)** Performance of bout-wise linear decoders trained to decode various pup-directed behaviors from the non-interaction period using MPOA^CalcR^ population activity. **(p)** Confusion matrix (left) and F1 score of a multi-class linear decoder trained to classify various pup-directed behaviors using MPOA^CalcR^ population activity. **(q)** Matrix (left) and bar plot (right) of the performance of linear decoders trained to distinguish between pairs of pup-directed behaviors using MPOA^CalcR^ population activity. Dashed red line indicates chance level. Numbers in parentheses or axis titles indicate cell number **(e, i-m)** or subject animal number **(o-q)**. Circles represent data of individual cells **(h, n)** or individual animals **(q)**. Circle-line pairs represent data of individual animals **(o, p)**. **(e)** Resampled distributions followed by permutation test against ‘no interaction’ period and Benjamini Hochberg false-discovery rate correction. **(f)** Spearman correlation. **(n, p)** Two-way ANOVA with repeated measures followed by Tukey’s multiple comparisons test (between-decoder test results not shown). **(o)** Paired t-test. **(q)** Red values on bars: one-sample t-test against chance performance (0.5) followed Benjamini Hochberg false-discovery rate correction. Between-decoder performance: Friedman test followed by Dunn’s multiple comparisons test. Lines and shaded error bars represent mean ± SEM. All p < 0.05 are indicated.

We calculated each cell’s temporal responses (Z-scored ΔF/F from -10 to +10 seconds surrounding behavior onset) during pup approach, investigation, retrieval, grooming, and crouching and then performed principal components analysis (PCA) based on these responses^50^ (**Extended Data Fig. 8**). We then performed k-means clustering based on scores of the first 4 PCs, which accounted for 92% of the variance, and identified 4 clusters (**Extended Data Fig. 3g, h**). The 1^st^ cluster (33 cells, green) showed high PC1 scores and high activities during active components of pup interaction, including pup approach, investigation, grooming, and retrieval, with the highest activity increase during retrieval, while crouching is accompanied by activity decline (**Fig. 3e, i-n**). The 2^nd^ cluster (24 cells, magenta) had high PC2 scores and was primarily tuned to pup grooming and crouching while being mildly modulated by approach and investigation (**Fig. 3e, i-n**). The third cluster (80 cells, blue) showed increased activity during pup approach and investigation; however, unlike the first cluster, activity during retrieval did not increase further beyond the level observed during investigation (**Fig. 3e, i-n**). The 4^th^ cluster (76 cells, grey) was largely non-responsive during pup-directed behaviors, although 10 cells showed mildly increased activity during crouching (**Fig. 3e, i-n**).

Using linear decoders, we found that all pup-directed behaviors, including pup approach, investigation, retrieval, grooming, and crouching, could be reliably decoded from MPOA^CalcR^ population activity relative to non-interaction periods (**Fig. 3o**). To further determine whether MPOA^CalcR^ activity distinguishes among distinct pup-directed behaviors, we trained a five-class linear decoder to classify these behaviors. The F1 scores for all five behaviors were significantly above label-shuffled controls, indicating that each behavior is associated with a discriminable activity pattern (**Fig. 3p**). Among them, crouching and grooming were decoded with the highest accuracy, and misclassifications between these two behaviors were more common than confusion with pup approach, investigation, or retrieval (**Fig. 3p**). In contrast, decoding performance was lowest for pup approach and retrieval, which were most often misclassified as pup investigation (**Fig 3p, left**). Consistent results were obtained using pairwise decoders trained to distinguish individual pairs of pup-directed behaviors: decoding accuracy was lowest for approach versus investigation and retrieval versus investigation, and highest for comparisons between active pup-directed behaviors and crouching or pup grooming (**Fig. 3q**). Together, these findings suggest that active pup-directed behaviors, which often occur in close temporal succession, are represented by overlapping but non-identical activity patterns in MPOA^CalcR^ neurons. This interpretation is further supported by the strong correlations among activity patterns associated with active pup-directed behaviors, whereas grooming- and crouching-related activity patterns were more distinct (**Fig. 3e, f**)

The activity elicited by each behavior varied across animals, likely due to differences in imaging sites along the anterior-posterior axis (**Extended Data Fig. 9**). Specifically, the animal with the most anteriorly placed GRIN lens (Bregma: 0.14 mm) showed weak responses during pup-directed behaviors in general, and during active components of pup interaction in particular (**Extended Data Fig. 9a**). However, pup investigation, retrieval, and pup grooming elicited large responses in animals with posteriorly placed GRIN lenses (**Extended Data Fig. 9c-f**). The average population response during these behaviors showed a trend toward positive correlation with the GRIN lens anterior-posterior placement (**Extended Data Fig. 9h-j**). These results, together with recent population Ca^2+^ recordings in females^51^, support the involvement of MPOA^CalcR^ cells, especially those located posteriorly, in various aspects of parental behaviors.

### Changes in MPOA^CalcR^ responses over pup sensitization

To determine how MPOA^CalcR^ cell responses change as animals become parental, we longitudinally tracked the cell responses before and after pup sensitization. We recorded cell responses as animals investigated pups, adult male, and adult female intruders before pup sensitization and after paternal behavior onset (**Fig. 4a**). A total of 131 cells from 7 animals could be confidently matched before and after pup sensitization (**Fig. 4b**). **Fig. 4c** shows recordings from one example animal. Across animals, 18% (23/131) of neurons responded during pup investigation before pup sensitization (**Fig. 4d, e**). When animals became parental, as defined by pup retrieval, the percentage of pup investigation-responsive cells more than doubled (49/131) (**Fig. 4d, e**). In contrast, the fraction of adult female and adult male responsive cells dropped by approximately 10% (adult female: 45% to 37%; adult male: 37% to 24%) (**Fig. 4d, e**). Consistent with a net increase of pup-responsive cells, the mean magnitude of Ca^2+^ activity during pup investigation increased in paternal animals across all tracked cells (**Fig. 4f**). In contrast, the population responses were largely unchanged during male and female investigation (**Fig. 4g, h**).

**Fig. 4.**
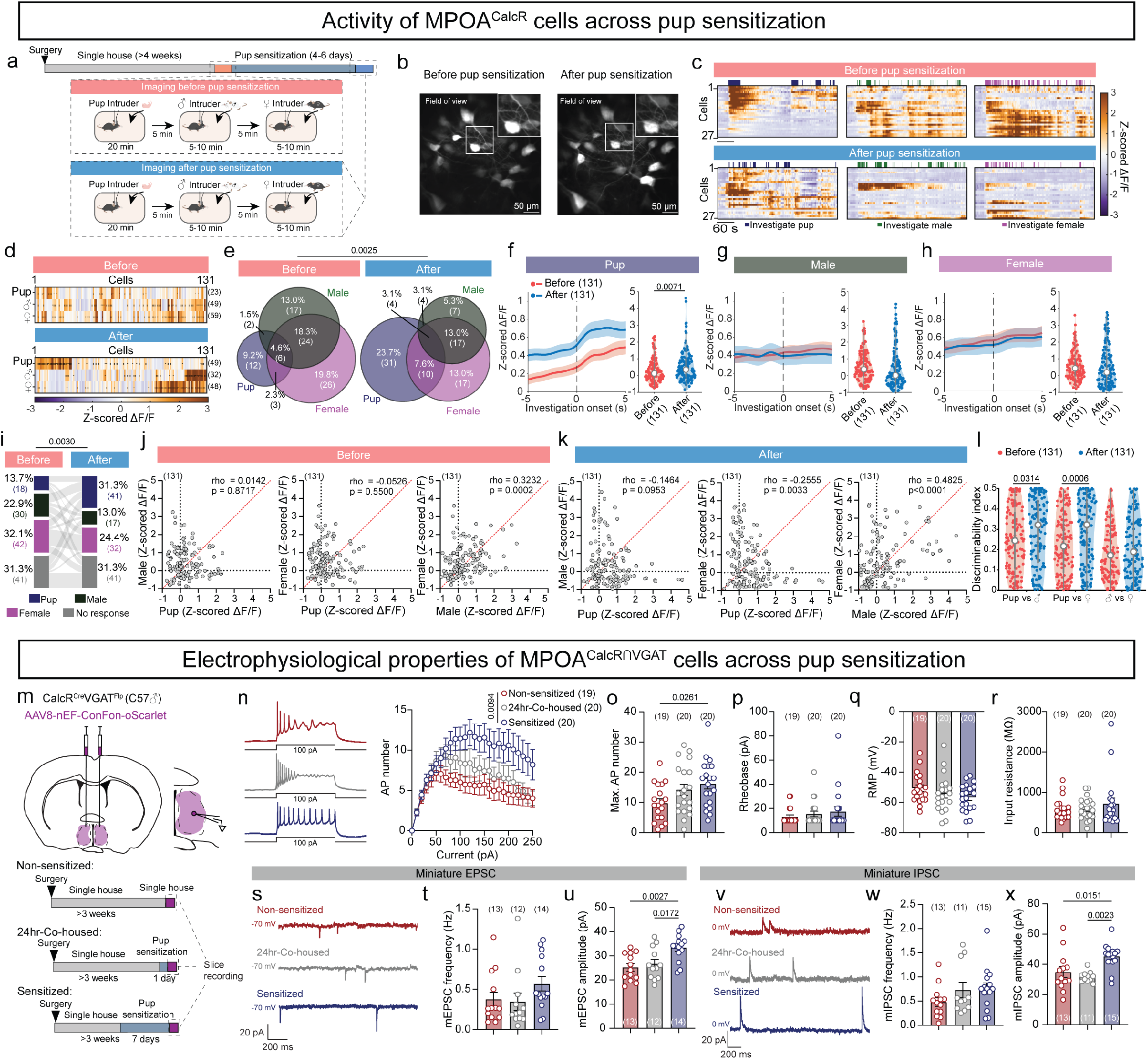
Plasticity of MPOA^CalcR^ activity across pup sensitization. **(a)** Timeline to image MPOA^CalcR^ cell activity during pup, male, and female interactions before and after pup sensitization. **(b)** Example session-averaged imaging plane showing GCaMP6s-expressing cells across two sessions (before and after pup sensitization). **(c)** Representative heatmaps showing Z-scored ΔF/F of longitudinally-imaged MPOA^CalcR^ cells from the mouse shown in **b** before (top) and after pup sensitization (bottom) when encountering pups (left), an adult male intruder (middle), and an adult female intruder (right). Cells are arranged in the same order before and after pup sensitization. **(d)** Heatmaps showing averaged Z-scored ΔF/F of MPOA^CalcR^ cells during pup, male, and female investigation. Cells are arranged in the same order before (top) and after pup sensitization (bottom) based on their membership in k-means clustering analyses (n = 131 longitudinally-imaged cells from 7 mice). Numbers on the right indicate the number of significantly responsive cells. Black circles indicate q values. ●q<0.01; ●●q<0.001. **(e)** Venn diagrams showing cells significantly responsive to pup, male, and female investigation before and after pup sensitization. **(f-h)** Left: population-averaged PETHs during pup **(f)**, male **(g)**, and female **(h)** investigation. Dashed lines indicate the onset of behavior. Right: Average Z-scored ΔF/F of each cell during pup **(f)**, male **(g)**, and female **(h)** investigation. Red: before pup sensitization; Blue: after pup sensitization. **(i)** Alluvial diagram showing pup-, male-, and female-preferring cells and non-responsive cells (total of 131 cells from 7 mice) before and after pup sensitization. **(j, k)** Correlations of cell responses (Z-scored ΔF/F) during pup and male investigation (left), pup and female investigation (middle), and male and female investigation (right) before **(j)** and after **(k)** pup sensitization. **(l)** Discriminability index of cell responses (Z-scored ΔF/F) across stimulus pairs. **_(m)_** Virus injection, recording schematic, and experimental timeline for ex vivo whole-cell recordings of _MPOA_^CalcR∩VGAT^ _cells._ **(n)** Left: Representative current-clamp recording traces of MPOA^CalcR∩VGAT^ cells after 100pA current injection. Right: F-I curve of MPOA^CalcR∩VGAT^ cells in non-sensitized, 24-hour-cohoused, and pup sensitized mice (non-sensitized: n = 19 cells from 3 mice; 24-hr-co-housed: n = 20 cells from 3 mice; sensitized: n = 20 cells from 3 mice). **(o-t)** Maximum action potential number across current steps **(o)**, rheobase **(p)**, resting membrane potential **(q)**, and input resistance **(r)** of MPOA^CalcR∩VGAT^ cells with various pup exposure experiences (non-sensitized: n = 19 cells from 3 mice; 24-hr-co-housed: n = 20 cells from 3 mice; sensitized: n = 20 cells from 3 mice). **(s, v)** Representative voltage-clamp recording traces of MPOA^CalcR∩VGAT^ cells in 1 μM TTX. Numbers indicate the holding potential. **(t, u, w, x)** The frequency **(t, w)** and amplitude **(u, x)** of mEPSCs **(t, u)** and mIPSCs **(w, x)** of MPOA^CalcR∩VGAT^ cells in mice with various pup exposure experiences (non-sensitized: n = 13 cells from 3 mice; 24-hr-co-housed: n = 12, 11 cells from 3 mice; sensitized: n = 14, 15 cells from 3 mice). Numbers in parentheses or axis titles indicate cell number. Circles represent data of individual cells. **(d)** One-sample t-test followed by Benjamini Hochberg false-discovery rate correction. **(e, i)** Chi-square test. **(f-h)** Permutation test. **(j, k)** Spearman correlation. **(l)** Two-way ANOVA with repeated measures followed by Tukey’s multiple comparisons test. **(n)** Two-way ANOVA with repeated measures followed by Tukey’s multiple comparisons test. **(o, q, u, x)** Ordinary one-way ANOVA. **(p, r, t, w)** Kruskal-Wallis test. Bars and error bars represent mean ± SEM. Lines and shaded error bars represent mean ± SEM. Violin plots: white circles represent the median and interquartile and 1.5x interquartile ranges are represented by thick and thin gray bars, respectively; shaded areas represent the range of all values. All p < 0.05 are indicated.

We wondered whether parental behavior-responsive cells are predisposed to be activated by pups before the emergence of parental behaviors and examined pre-sensitization MPOA^CalcR^ cell social cue responses (**Extended Data Fig. 10**). Among the four MPOA^CalcR^ clusters identified based on their post-sensitization responses during parental behaviors (**Fig. 3e**), the 1^st^ (retrieval cells) and 2^nd^ cluster (grooming and crouching cells) responded preferentially to adult stimuli before sensitization (**Extended Data Fig. 10a, b, e, f, i, j**). After sensitization, both clusters decreased their responses during adult investigation and increased their responses to pups (**Extended Data Fig. 10a, b, e, f, i, j**). The 3^rd^ cluster, i.e., moderately responsive during approach and retrieval, responded similarly to all social stimuli before sensitization and showed a trend of increased responses to pups after sensitization (**Extended Data Fig. 10c, g, k**). The 4^th^ cluster, i.e., the parental behavior non-responsive cells, showed no activity increase during pup investigation either before or after sensitization and responded overall more to adult stimuli (**Extended Data Fig. 10d, h, l**). These results suggest that MPOA parental cells are not wired to receive pup cues exclusively, although parental behavior-unrelated cells may receive minimal pup-related sensory inputs.

To further understand the flexibility of sensory inputs to the MPOA cells, we compared the preferred response of each cell, defined by the social stimulus that evokes the largest significant response, before and after pup sensitization (**Fig. 4i**). After pup sensitization, pup-preferring cells increase from 14% (18/131) to 31% (41/131), while adult-preferring cells decreased by roughly the same percentage (55% to 37%). Among pup-preferring cells, while a portion of neurons consistently responded to the pup preferentially (8/41) before and after the emergence of parental behaviors, other cells switched from previously male-preferring (12/41), female-preferring (9/41), or non-responsive profiles (13/41) to pup-preferring (**Fig. 4d, i**). Beyond a net increase of pup preferring responses, a subset of cells shifted their responses from male preferring to female preferring (**Fig. 4d, i**). These results further support that MPOA^CalcR^ cell response profile to social sensory inputs is plastic.

To address whether the responses of MPOA^CalcR^ cells become more selective with the transition to alloparental care, we first examined the pairwise cross-correlation among pup, adult male, and adult female responses. MPOA^CalcR^ cell responses to pups and adult males or females were uncorrelated before pup sensitization. The responses became slightly but significantly anti-correlated to females and showed a trend of anti-correlation to males after parental behavior onset (**Fig. 4j, k**). In contrast, responses during male and female investigation were significantly positively correlated both before and after pup sensitization (**Fig. 4j, k**).

We next computed the choice probability^52,53^ for each neuron to estimate how the cell response differs between two social stimuli. Specifically, ΔF/F distribution histograms of each pair of stimuli were used to generate a Receiver Operating Characteristic (ROC) curve, and the absolute deviation of the area under the ROC (auROC) from 0.5 was used to measure the cell’s ability to discriminate between the two stimuli (0: indiscriminate; 0.5: fully distinct) (**Extended Data Fig. 11**). After sensitization, the discrimination ability of MPOA^CalcR^ cells between pup and adult stimuli, but not between adult male and female, increased significantly (**Fig. 4l**).

To address whether MPOA^CalcR^ cell responses to social cues change without pup sensitization, we additionally recorded the activity of 145 MPOA^CalcR^ cells from 7 single-housed male mice between 2 sessions, with 3-6 days apart, and examined the responses during investigating pups, adult males, and adult females (**Extended Data Fig. 12a**). At the population level, a largely stable number of neurons (pup: 30 vs. 27; female: 47 vs. 55; male: 54 vs. 43) responded during investigation of each social target between these two sessions (**Extended Data Fig. 12b, c**). The average magnitude of responses to pups and adult males remained consistent across the two sessions, while the responses to females showed a trend of increase in the second recording session (**Extended Data Fig. 12d-f**). When examining the preferred response of individual cells, unlike pup-sensitized males, we found no increase in the number of pup-preferring cells (**Extended Data Fig. 12g**). However, we did find changes in individual cell response tuning between the two sessions. Most noticeably, a subset of cells switched from male-preferring to female-preferring or non-responsive (**Extended Data Fig. 12g**). Across the population, the male and female investigation responses were highly correlated with each other and largely uncorrelated with pup responses in both recording sessions (**Extended Data Fig. 12h, i**). The discriminability between pup and female, but not pup vs. male, showed a slight increase in the second session, although it is lower than the discriminability after pup sensitization (**Extended Data Fig. 12j**). These results suggest increased pup response is unique after pup sensitization, although short periods of social experience with pups and adults may enhance MPOA^CalcR^ cell response selectivity to some extent.

When examining response profiles of individual animals, we found that the tuning of MPOA^CalcR^ neurons to social stimuli differed along the anterior-posterior axis of the MPOA. Interestingly, MPOA^CalcR^ cells showed a strong adult-biased response pattern in animals with the most anteriorly located imaging sites (Bregma: 0.04 mm), whereas the posteriorly located MPOA^CalcR^ cells preferentially responded to pups **(Extended Data Fig. 13**). This tuning difference was present before pup sensitization and largely persisted following sensitization, suggesting an experience-independent topographical organization of MPOA^CalcR^ cell tuning to social stimuli (**Extended Data Fig. 13h, i**).

Altogether, these single-cell longitudinal imaging results suggest that, first, many MPOA^CalcR^ cells are activated during parental behaviors; second, MPOA^CalcR^ cell responses to pups expand after pup sensitization; third, MPOA^CalcR^ cells increase discriminability between pup cues and adult social cues after pup sensitization although short periods of social interactions may also induce response divergence; third, individual MPOA^CalcR^ cells could respond to multiple social targets, and their sensory tuning could change with social experience; fourth, MPOA^CalcR^ cell responses during adult male and female investigation are more similar than those during pup investigation at the population level; fifth, responses of anterior MPOA^CalcR^ cells are adult-biased whereas those in the posterior part are pup-biased regardless of animals’ experience with pups.

### Physiological changes of MPOA^CalcR^ cells over pup sensitization

To determine whether in vivo response changes are supported by cell physiological changes, we performed in vitro patch-clamp recordings of MPOA^CalcR∩VGAT^ cells (**Fig. 4m**). We labeled MPOA^CalcR∩VGAT^ cells with Cre- and Flp-dependent oScarlet and collected brain slices from male mice after undergoing 0, 1, or 7 days of pup sensitization (**Fig. 4m**). Compared with non-sensitized animals, MPOA^CalcR∩VGAT^ cells from 7-day-sensitized male mice showed enhanced firing upon current injection, yielding a significantly higher maximal firing rate (**Fig. 4n, o**). This change appeared to occur quickly and gradually: animals undergoing 24 hours of pup exposure and whose behavior had not yet switched to parenting already showed a trend of increased excitability (**Fig. 4n, o**). Other parameters, including rheobase, resting membrane potential (RMP), and input resistance, did not differ across groups (**Fig. 4p-r**).

To understand whether pup sensitization also induces synaptic plasticity in MPOA^CalcR∩VGAT^ cells, we recorded miniature excitatory and inhibitory postsynaptic currents (mEPSCs and mIPSCs). Compared to both non-sensitized and 24-hour pup-exposed animals, MPOA^CalcR∩VGAT^ cells in 7-day pup sensitized males showed increased mEPSC amplitude (**Fig. 4s, u**). The mEPSC frequency also showed an upward trend (**Fig. 4t**). Likewise, mIPSC amplitude increased in sensitized animals compared to both other groups (**Fig. 4v-x**). These results suggest that pup sensitization, particularly to the point that parental behaviors emerge, is associated with both synaptic changes and increased excitability of MPOA^CalcR∩VGAT^ cells.

### Disrupting CalcR signaling in the MPOA impairs paternal behaviors

To address whether CalcR is itself functionally important for the emergence and expression of parental behaviors in virgin male mice, we generated viruses expressing short hairpin RNA (shRNA) against *Calcr* or scrambled shRNA controls and injected them into the MPOA (**Fig. 5a**). Immunostaining revealed that *Calcr* shRNA reduced CalcR-positive cells by approximately 60% (**Fig. 5b, c**). Patch-clamp recordings in non-sensitized males showed that MPOA^CalcR∩VGAT^ cells expressing *Calcr* shRNA and scrambled shRNA did not differ in cell excitability, mEPSCs, or mIPSCs, suggesting that *Calcr* knockdown does not alter the cells’ basal physiological properties (**Extended Data Fig. 14**).

**Fig. 5.**
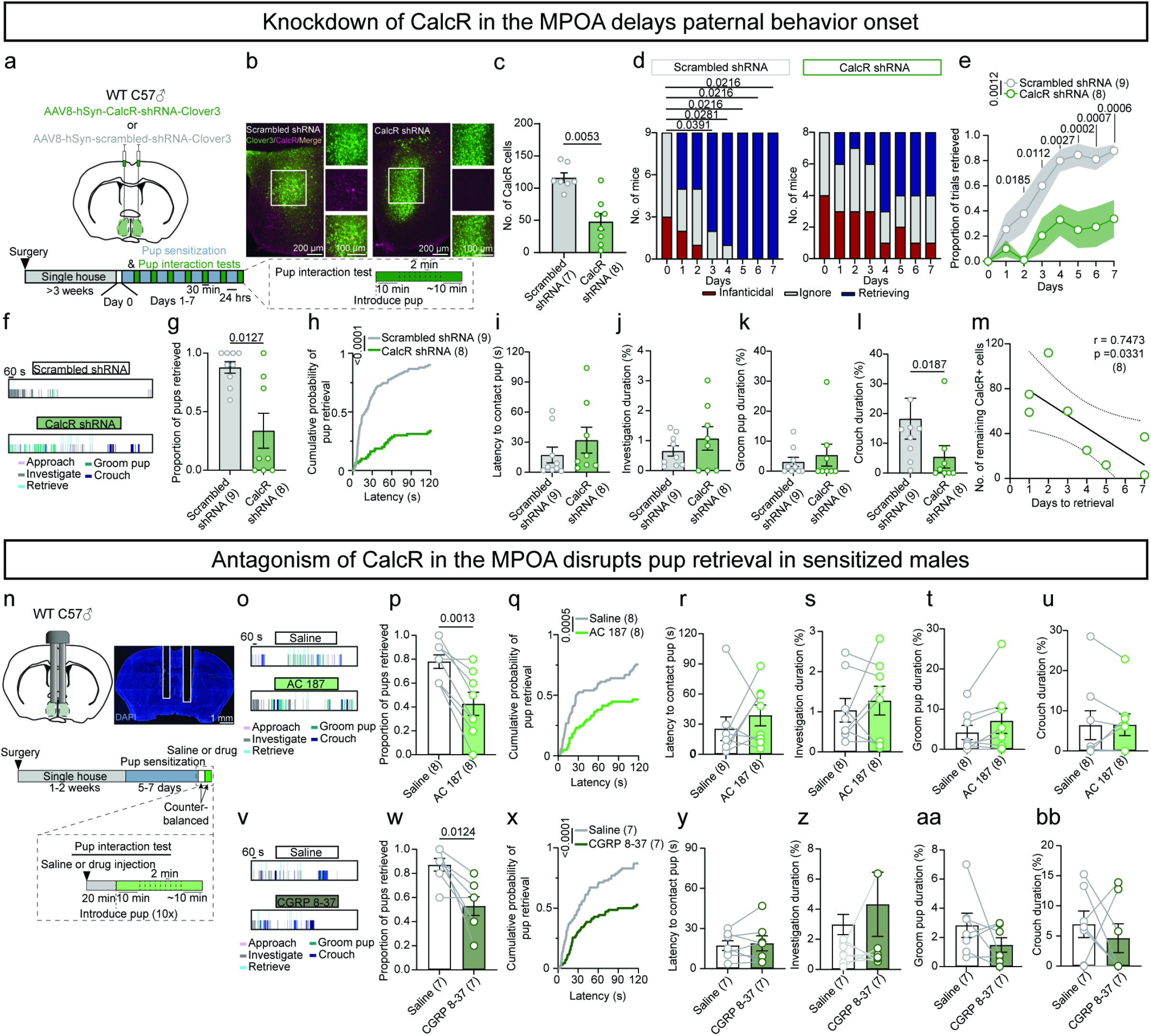
MPOA CalcR signaling is necessary for parental behaviors. **(a)** Virus injection schematic and timeline to test behaviors across pup sensitization. **(b)** Representative histology images showing scrambled shRNA (left) or calcitonin receptor (CalcR) shRNA (right; green) and CalcR immunoreactivity (IR; magenta) in the MPOA. **(c)** Quantification of CalcR-expressing cells in scrambled shRNA- and CalcR shRNA-injected animals. **(d)** The number of scrambled shRNA and CalcR shRNA animals that were infanticidal, non-infanticidal and non-retrieving (ignore), and retrieving in the pup interaction test across days of pup sensitization (day 0: before pup sensitization, days 1-7: during pup sensitization). **(e)** Proportion of pups retrieved in the pup interaction test across days of pup sensitization (day 0: before pup sensitization, days 1-7: during pup sensitization). **(f)** Representative raster plots showing pup-directed behaviors on day 7 of the pup interaction test from a scrambled shRNA mouse and a CalcR shRNA mouse. **(g-l)** The proportion of pups retrieved **(g)**, cumulative probability of pup retrieval over the 120 s testing period **(h),** latency to contact pups **(i)**, pup investigation duration **(j)**, grooming pup duration **(k)**, crouching over pups duration **(l)** in scrambled shRNA and CalcR shRNA mice during the pup interaction test on day 7 of the experiment. **(m)** Correlation of the number of immunoreactive CalcR cells remaining in the MPOA and pup retrieval onset across pup sensitization in CalcR shRNA-injected animals. **(n)** Cannula implant schematic, representative histology image, and timeline to test behavior in pup-sensitized mice following saline or drug infusion into the MPOA. **(o)** Representative raster plots showing pup-directed behaviors following saline or AC187 infusion during the entire pup interaction test. **(p-u)** The proportion of pups retrieved **(p)**, cumulative probability of pup retrieval **(q)**, latency to contact pups **(r)**, pup investigation duration **(s)**, grooming pup duration **(t)**, and crouching over pups duration **(u)** in animals after saline or AC187 infusion. **(v)** Representative raster plot showing pup-directed behaviors following saline or CGRP 8-37 infusion. **(w-bb)** The proportion of pups retrieved **(w)**, cumulative probability of pup retrieval **(x)**, latency to contact pups **(y)**, pup investigation duration **(z)**, grooming pup duration **(aa)**, crouching over pups duration **(bb)** in animals after saline or CGRP 8-37 infusion. Numbers in parentheses or axis titles indicate the subject animal number. Circles in **g-m** and circle-line pairs in **p-u** and **w-bb** represent data of individual animals. **(c, g, i, k, l)** Mann-Whitney U test. **(d)** Cochran’s Q test followed by Benjamini Hochberg false-discovery rate correction. **(e)** Two-way ANOVA with repeated measures followed by Tukey’s multiple comparisons test. **(h, q, x)** Logrank (Mantel-Cox) test. **(j)** Unpaired t-test. **(m)** Simple linear correlation. **(p, s, w, x, z-bb)** Paired t-test. **(r, t, u, y)** Wilcoxon test. Bars and error bars represent mean ± SEM. Lines and shaded error bars represent mean ± SEM. All p < 0.05 are indicated.

Three weeks after the virus injection and before pup sensitization, a comparable number of *Calcr* shRNA (4/8) and scramble shRNA-injected males (3/9) showed spontaneous infanticide, consistent with no change in MPOA^CalcR∩VGAT^ cell properties after *Calcr* knockdown in naïve males (**Fig. 5d, Extended Data Fig. 14**). Over the course of 7-day pup sensitization, fewer *Calcr* shRNA animals showed pup retrieval compared to scrambled shRNA-injected control animals, and the retrieval reliability was consistently lower in *Calcr* shRNA animals (**Fig. 5d, e**). When comparing *Calcr* shRNA and control animals after 7 days of pup sensitization, *Calcr* shRNA animals took longer to start retrieving, retrieved fewer pups, and crouched over pups for a shorter period (**Fig. 5f-l**). The onset of pup retrieval was negatively correlated with the number of remaining CalcR-positive cells: the fewer the cells that remained, the slower the onset (**Fig. 5m**). These results suggest that CalcR in the MPOA is important for the emergence and/or expression of paternal behavior.

CalcR contributes to multiple receptors: it can, by itself, form calcitonin receptors and bind calcitonin, or form heterodimers with three receptor activity-modifying proteins (RAMPs 1-3), resulting in amylin receptors 1-3 (Amy1-3)^54^. All amylin receptors bind to amylin, a pancreatic peptide also produced in the hypothalamus of female mice^37,55,56^. Amy1 and Amy3 also bind to CGRP. In particular, Amy1 has a similarly high binding affinity to amylin and CGRP and is considered a dual receptor^54,57^. In the MPOA, previous RNAseq and in situ hybridization studies revealed expression of RAMP1^35^ and RAMP2^37^, but not RAMP3, suggesting the potential presence of Amy1 and Amy2 in the area.

To more directly test which forms of CalcR may contribute to the paternal behavior expression following pup sensitization, we implanted cannulas over the MPOA to block the receptor action locally in paternal virgin male mice (**Fig. 5n**). We found that both the general amylin receptor 1-3 blocker, AC187, and the CGRP receptor and Amy1 dual blocker, CGRP 8-37, significantly reduced the probability of pup retrieval and increased retrieval latency but did not affect other aspects of parental behaviors in pup-sensitized male mice (**Fig. 5o-bb**). No animal attacked pups after either antagonist injection. Given that Amy1 is the only receptor blocked by both AC187 and CGRP8-37, we consider Amy1 the likely receptor mediating the behavioral changes induced by the antagonists. We could not test the contribution of the calcitonin receptor due to a lack of specific antagonists^54,58,59^. Nevertheless, given that calcitonin is synthesized exclusively peripherally, the calcitonin receptor, on its own, is likely to have limited central functions^60,61^. Notably, compared with *Calcr* knockdown, which reduced receptor expression throughout the pup sensitization process, the parental behavior deficits caused by acute AC187 or CGRP 8-37 injection are relatively minor, suggesting that CalcR likely plays a more critical role in the emergence than in the acute expression of parental behaviors. Lastly, injecting AC187 or CGRP 8-37 into the MPOA of non-sensitized males did not alter pup-directed behaviors, such as infanticide probability or pup interaction duration, suggesting that MPOA CalcR signaling contributes little to modulating pup-directed behaviors in non-parental males (**Extended Data Fig. 15**).

### CalcR activation facilitates pup-relevant inputs to MPOA^CalcR∩VGAT^ cells

The functional and imaging results suggest that activation of CalcR-containing receptors, e.g., Amy1, may facilitate the MPOA^CalcR^ cell response increase to pups after repeated pup exposure, leading to increased parental behaviors. To test this hypothesis, we first aimed to identify an excitatory input to the MPOA that could carry pup cues. We previously found that posterior amygdala (PA) cells expressing estrogen receptor alpha (Esr1) are responsive to social cues and provide strong excitatory inputs to the MPOA^62^. A recent study further suggested that a subset of MPOA-projecting PA cells is uniquely activated during parental care in fathers and pharmacological activation of the cells can suppress infanticide and promote parental care in naïve males^63^.

We thus hypothesize that PA could be a relevant upstream region providing pup-related sensory information to the MPOA during pup sensitization. To this end, we used fiber photometry to record the calcium signal of PA^Esr1^ cells projecting to the MPOA (PA^Esr1◊MPOA^) by injecting a retrograde herpes simplex virus (HSV) encoding a Cre-dependent GCaMP6f (HSV-hEF1α-GCaMP6f) into the MPOA of *Esr1-2A-Cre* (Esr1^Cre^) male mice and implanted optic fibers above the PA (**Fig. 6a, b**). Ca^2+^ activity increased when animals approached pups and rose significantly during pup investigation (**Fig. 6c, e, f**). During pup attack, however, no consistent increase was observed (**Fig. 6g**). When the males investigated pups placed in a wire-mesh ball, as is the case during pup sensitization, PA^Esr1◊MPOA^ cell activity increased consistently (**Fig. 6d, h**). These results indicate that PA is well-positioned to relay pup sensory cues to the MPOA during pup sensitization. We next used PA input to investigate how pup-relevant inputs can affect the physiology of the MPOA. First, we confirmed that PA provides direct excitatory inputs to the MPOA^CalcR∩VGAT^ cells. Four weeks after virally expressing Chronos^64^ in the PA and oScarlet^43^ in MPOA^CalcR∩VGAT^ cells, we performed whole-cell voltage-clamp recordings of MPOA^CalcR∩VGAT^ cells on brain slices while optogenetically activating PA-MPOA terminals (**Fig. 6i, j**). We found that 1 ms light pulses evoked excitatory postsynaptic current (oEPSC) in 23/28 of recorded cells (**Fig. 6j-l**). The short latency of the oEPSC (2.82 ± 1.3 ms) indicated the monosynaptic nature of the connection (**Fig. 6k**).

**Fig. 6.**
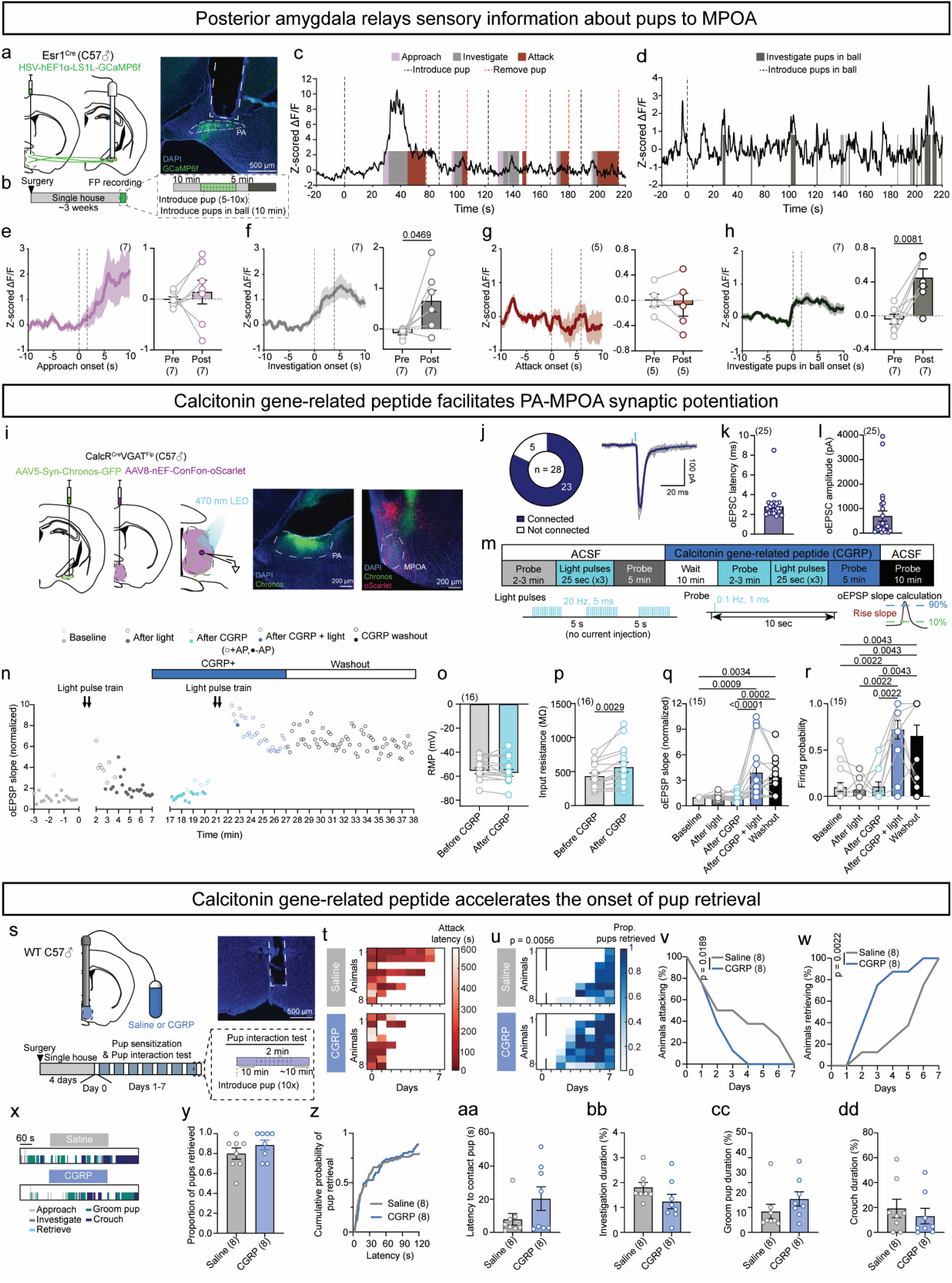
CalcR activation facilitates the potentiation of pup-relevant inputs to MPOA. **(a)** Virus injection and optical fiber implant strategy and representative histology image. Dashed white lines indicate optical fiber track and the posterior amygdala (PA) location. **(b)** Timeline to record MPOA-projecting posterior amygdala Esr1 neurons (PA^Esr1◊MPOA^) during pup interactions. **(c)** Representative Ca^2+^ trace showing Z-scored ΔF/F of PA^Esr1◊MPOA^ cells during unrestrained pup interactions. **(d)** Representative Ca^2+^ trace showing Z-scored ΔF/F of PA^Esr1◊MPOA^ cells during interactions with pups placed in a wire mesh ball. **(e-h)** Left: average PETHs of Z-scored ΔF/F of PA^Esr1◊MPOA^ cells during pup-directed approach **(e)**, unrestrained pup investigation **(f)**, pup-directed attack **(g)**, and investigation of pups in a wire mesh ball **(h)**. Dashed lines indicate the onset and average offset of behavior. Right: Average Z-scored ΔF/F of PA^Esr1◊MPOA^ cells during various pup-directed behaviors. **(i)** Virus injection and recording schematic and representative histology images from the PA and MPOA. Dashed white lines indicate the locations of PA and MPOA. **(j)** Left: Pie chart showing the number of MPOA cells connected (blue) and not connected (white) with PA. Right: Example optically-evoked excitatory post-synaptic current (oEPSC) trace recorded from an MPOA^CalcR∩VGAT^ cell, showing the oEPSC elicited by 1-ms light stimulation of PA axon terminals. **(k, l)** Latency **(k)** and amplitude of oEPSCs **(l)** of MPOA^CalcR∩VGAT^ cells following 1-ms light stimulation of PA axon terminals. **(m)** Recording and light delivery protocols and illustration of oEPSP slope calculation to probe and induce synaptic potentiation and its modulation by calcitonin gene-related peptide (CGRP). **(n)** Slopes of oEPSP of an example MPOA^CalcR∩VGAT^ cell during baseline (light gray), after light stimulation (dark gray), after CGRP perfusion (teal), after light stimulation in the presence of CGRP (blue), and during washout (black). Blue bar represents the period of CGRP perfusion. White bar with black outlines represents the washout period. Open circles represent oEPSPs with action potentials (AP), and closed circles represent those without AP. **(o, p)** Resting membrane potential (RMP) **(o)** and input resistance **(p)** before and after CGRP perfusion (n = 16 cells from 6 mice). **(q, r)** Normalized oEPSP slope **(q)** and light stimulation-evoked firing probability **(r)** at baseline, after light stimulation without CGRP, after CGRP perfusion, after light stimulation in the presence of CGRP, and during washout (n = 15 cells from 6 mice). **(s)** Cannula implant schematic, representative histology image, and timeline to test behavior across pup sensitization during continuous saline or drug infusion into the MPOA. **(t)** The latency to attack pups across the duration of the experiment (day 0: before pup sensitization, days 1-7: during pup sensitization). **(u)** The proportion of pups retrieved across the duration of the experiment (day 0: before pup sensitization, days 1-7: during pup sensitization). **(v)** Percentage of animals attacking pups in the pup interaction test across the duration of the experiment (day 0: before pup sensitization, days 1-7: during pup sensitization). **(w)** Percentage of animals retrieving pups in the pup interaction test across the duration of the experiment (day 0: before pup sensitization, days 1-7: during pup sensitization). **(x)** Representative raster plots showing pup-directed behaviors in the pup interaction test on day 7 of the experiment in a saline- and a CGRP-infused animal. Raster plots represent the 10 minutes following the introduction of the last pup. **(y-dd)** The proportion of pups retrieved **(y)**, cumulative probability of pup retrieval **(z)**, latency to contact pups **(aa)**, pup investigation duration **(bb)**, grooming pup duration **(cc)**, and crouching over pups duration **(dd)** in saline- and CGRP-injected mice during the pup interaction test on day 7 of the experiment. Numbers in parentheses or axis titles indicate the subject animal number **(e-h, t-dd)** or cell number **(k, l, o-r)**. Circles and circle-line pairs represent data of individual animals **(e-h)** or individual cells **(k, l, o-r)**. **(e, g, h, o, p)** Paired t-test. **(f)** Wilcoxon test. **(q, r)** Friedman test followed by Dunn’s multiple comparisons test. **(t, u)** Two-way ANOVA with repeated measures followed by Tukey’s multiple comparisons test. **(v, w, z)** Log rank (Mantel-Cox) test. **(y, aa-dd)** Mann-Whitney U test. Bars and error bars represent mean ± SEM. All p < 0.05 are indicated.

We then asked whether CalcR-containing receptor activation can change MPOA^CalcR∩VGAT^ cell activity and responses to the PA input. We considered Amy1 the most likely receptor of interest and bath applied 200 nM amylin or CGRP, two potential Amy1 ligands. Neither ligand changed MPOA^CalcR∩VGAT^ cell RMP, spiking activity, or PA-stimulation evoked excitatory postsynaptic potential (oEPSP) (**Fig. 6n-r, Extended Data Fig. 16a-e**). However, 200 nM CGRP, but not Amylin, consistently increased the input resistance of the recorded cells (**Fig. 6p, Extended Data Fig. 16c**). Additionally, a high concentration of amylin (800 nM) depolarized the cells moderately, although no firing change was observed, consistent with the fact that Amy1 is coupled to an excitatory GPCR, but it also suggests low receptor sensitivity to amylin (**Extended Data Fig. 16g, j**).

Given the likely role of CalcR-containing receptor in facilitating the emergence of parental behaviors and synaptic changes after pup sensitization (**Fig. 5a-m**), we further asked whether Amy1 can promote the synaptic plasticity of the PA-MPOA inputs that could carry pup-related cues. We activated PA inputs at a high frequency (5 ms, 20 Hz, 25 s each for 3 times) to mimic the increased PA input during pup sensitization and probed (1ms, 0.1 Hz) the postsynaptic responses of MPOA^CalcR∩VGAT^ cells before and after the high-frequency light stimulation in the absence or presence of amylin or CGRP (**Fig. 6m-r, Extended Data Fig. 16a-j**). We found that in the absence of any ligand, oEPSPs increased transiently following the high-frequency PA terminal stimulation but returned to baseline within a few minutes (**Fig. 6n, q, Extended Data Fig. 16a, d, f, i, k, n, p, s**). Strikingly, in the presence of CGRP, the high-frequency PA terminal stimulation tripled the oEPSP slope, and the effect persisted throughout the recording after CGRP washout (>10 min after washout onset) (**Fig. 6n, q**). The PA-evoked spiking probability also increased significantly and was maintained after the washout (**Fig. 6r**). Interestingly, neither 200 nM nor 800 nM amylin application significantly potentiated the PA-stimulation-induced synaptic plasticity of MPOA^CalcR∩VGAT^ cells, further suggesting amylin as a poor ligand for the MPOA^CalcR∩VGAT^ cells in male mice (**Extended Data Fig. 16a-j**). When CGRP was co-applied with either the amylin receptor antagonist AC187 or the dual Amy1/CGRP receptor antagonist CGRP8–37, CGRP-facilitated synaptic potentiation was completely blocked (**Extended Data Fig. 16k-t**). These results suggest that Amy1 activation, possibly by CGRP, can facilitate synaptic potentiation during repeated pup exposure to increase MPOA^CalcR∩VGAT^ cell responses to pup cues and ultimately promote parenting in male mice.

Lastly, to confirm that CGRP is the relevant ligand facilitating MPOA plasticity during pup sensitization at the functional level, we implanted cannulas into the MPOA of wild-type male mice. The cannulae were connected to osmotic pumps to continuously infuse either saline or CGRP as the males underwent continuous pup sensitization (**Fig. 6s**). To avoid a floor effect, we selected animals that showed spontaneous infanticidal behavior for this experiment. Compared to controls, CGRP-treated animals stopped attacking pups more quickly (days to last infanticidal event, CGRP: mean ± SEM: 1.5 ± 0.9 days, saline: mean ± SEM: 2.8 ± 2.6 days) (**Fig. 6t, v**) and took fewer days to start retrieving pups (days to first retrieval, CGRP: mean ± SEM: 3.1 ± 1.4 days, saline: mean ± SEM: 5.3 ± 1.8 days) (**Fig. 6u, w**). However, once all males became parental after 7 days of pup sensitization, there was no significant difference in pup-directed behaviors between CGRP- and saline-infused groups (**Fig. 6x-dd**). In a separate cohort of naïve males, we injected saline or CGRP acutely into the MPOA and observed no increase in parental behaviors: 3/6 saline-injected males attacked and 3/6 ignored pups, 4/6 CGRP-injected males attacked and 2/6 ignored pups (**Extended Data Fig. 17**). Together, these results support that CGRP facilitates the emergence of parental behaviors during pup sensitization, although its acute role in promoting parental behaviors is limited.

## Discussion

In this study, we demonstrated that male mice can become parental via repeated pup interaction. This process depends on MPOA^CalcR^ cells; inhibition of this population impairs both the execution and acquisition of parental behaviors in male mice. In contrast, optogenetic activation of GABAergic but not glutamatergic MPOA^CalcR^ cells rapidly induced pup retrieval in virgin male mice. As the virgin males became parental, MPOA^CalcR^ cells responded more robustly and selectively to pups, whereas responses to adult social cues remained unchanged or decreased. Importantly, CalcR-containing receptors are themselves functionally important for the emergence of parental behaviors over pup sensitization, likely by facilitating the potentiation of synapses carrying pup cues.

### The function of MPOA^CalcR^ cells

MPOA^CalcR^ cells are critical for both the expression and emergence of parental behaviors in virgin males, similar to the critical role of these cells in the expression of maternal behaviors in females^37^. Acute chemogenetic inhibition of MPOA^CalcR^ cells reduced all aspects of parental behaviors. Tonic inhibition of the cells blocked the emergence of parental behaviors in all virgin males over 7 days of pup sensitization. After lifting the inhibition post-pup-sensitization, parental behaviors remained unexpressed, suggesting that MPOA^CalcR^ cell activation during pup exposure is required for the plasticity to occur, which leads to behavioral changes.

Strikingly, when MPOA^CalcR∩VGAT^ cells were optogenetically activated, naïve male mice began caring for the pups within seconds of stimulation. They quickly approached the pups, brought them back to the nest, and repeatedly groomed them. These results suggest that the circuit downstream of MPOA^CalcR∩VGAT^ cells is ready to drive the behavior in naïve males, highlighting the importance of MPOA^CalcR∩VGAT^ plasticity in the emergence of parental behavior. Notably, activating MPOA^CalcR^ cells only promotes pup approach but not specific caring behaviors, suggesting that non-GABAergic MPOA^CalcR^ cells likely counteract the output of MPOA^CalcR∩VGAT^ cells. Indeed, GABAergic and glutamatergic MPOA cells have been reported to mediate distinct behaviors and induce opposite valences^42^. In particular, while GABAergic cells promote parental behaviors, glutamatergic cells induce stress and suppress parental behaviors^42^. Our results also align with a previous study showing that MPOA^Esr1∩VGAT^ is more effective in driving male sexual behaviors than MPOA^Esr1^ cells^52^. Thus, the GABAergic MPOA population appears to be the social behavior-relevant subset in general^65^.

Our imaging results suggest that MPOA^CalcR^ cells are heterogeneous: while the majority of cells respond to at least one parental behavior, over one-third respond preferentially to adult males or females. Nevertheless, at the functional level, activation of MPOA^CalcR∩VGAT^ cells or inactivation of MPOA^CalcR^ cells does not alter male sexual or aggressive behaviors, suggesting that these cells are more important for driving parental care than for adult-directed behaviors.

### Responses of MPOA^CalcR^ cells

MPOA^CalcR^ cell responses can be separated into action-related and sensory cue-driven. For action-related responses, following pup sensitization, approximately two-thirds of MPOA^CalcR^ cells (clusters 1-3) are activated during at least one aspect of pup-directed behaviors. Among these, ∼15% (cluster 1) are most strongly activated during pup retrieval, whereas ∼10% (cluster 2) are strongly and selectively activated during pup grooming and crouching. The existence of distinct pup-retrieval and grooming MPOA^CalcR^ subpopulations suggests that these two pup-directed behaviors may be independently modulated. Consistent with this possibility, previous studies found that optogenetic activation of MPOA Gal-expressing cells induces pup grooming but not retrieval^38^, whereas activating MPOA^Esr1^ cells induces pup retrieval but not grooming^30^. Notably, MPOA^CalcR^ cells partially overlap with both Gal- and Esr1-expressing cells^35^ and can bi-directionally modulate both pup retrieval and grooming. Thus, MPOA^CalcR^ cells that co-express Gal or Esr1 may preferentially drive pup grooming or pup retrieval, respectively.

MPOA^CalcR^ cells also increase activity during pup and adult investigation, likely reflecting sensory-driven responses. Several insights emerge from examining the relationship between MPOA^CalcR^ responses during parental actions and to social sensory cues. We consider responses during parental actions to reflect the functional role of these neurons in parental behavior, whereas responses to social cues indicate the sensory inputs they receive.

First, MPOA parental behavior neurons do not exclusively receive inputs conveying pup-related information. In fact, cluster 1 and cluster 2 neurons, which show the strongest activation during parental behaviors, respond more strongly to adult cues than pup cues prior to sensitization. Thus, inputs carrying adult and pup cues both reach parental MPOA neurons in naïve animals. However, with repeated pup interactions, pup cues evoke stronger responses in parental neurons, while responses to adult cues decrease. This shift likely reflects synaptic potentiation of pup-cue–carrying inputs and depression of adult-cue–carrying inputs. Importantly, neurons not associated with parental behavior (cluster 4) showed minimal responses during pup investigation both before and after sensitization, suggesting that pup-cue-carrying inputs largely avoid these cells. Together, these observations suggest the following model: in naïve animals, pup-related inputs weakly target MPOA neurons that drive parental behavior while largely sparing neurons unrelated to parental actions, whereas adult-related inputs reach both populations. With repeated pup interactions, pup-related inputs strengthen their connections with parental neurons—likely facilitated by activation of CalcR-containing receptors—while adult-related inputs weaken their connections through currently unknown mechanisms. As a result, pup cues become more effective at driving parental neurons, thereby enhancing discrimination between pup and adult cues.

Second, after pup sensitization, we observed an expansion of pup-responsive population and increased discriminability between pup and adult stimuli. Interestingly, in non-sensitized males, although the number of pup-responsive neurons did not increase, we still observed a modest increase in social stimulus discriminability during the second recording session. This response divergence may arise from social interaction experience accumulated during the first recording session (∼1 hour total). Indeed, previous Ca^2+^ imaging in the male ventrolateral part of the ventromedial hypothalamus (VMHvl), a medial hypothalamic region interconnected with the MPOA, showed that 30 minutes of free interaction with females can induce divergence of VMHvl cell responses to males and females 24 hours later^66^. These findings suggest that social experience in general may enhance MPOA cell response selectivity, whereas pup sensitization is specifically required to expand MPOA^CalcR^ responses to pups.

Third, at the population level, we found that response patterns to adult male and adult female cues are highly correlated, both before and after pup sensitization. We speculate that these correlated responses reflect synaptic inputs commonly recruited by adult conspecifics of both sexes. Adult males and females resemble each other in visual appearance and tactile features, and although their chemosensory signals are distinguishable, they share many volatile components^67^. Consistent with this interpretation, our previous single-unit recordings in female mice also revealed highly correlated MPOA neuronal responses to adult male and female cues^30^. In contrast, MPOA^CalcR^ cell responses to adult and pup cues are largely uncorrelated in naïve animals, suggesting that adult- and pup-related inputs target MPOA neurons through largely independent pathways. After pup sensitization, as parental neurons show increased responses to pups and decreased responses to adults, the responses evoked by pup and adult cues become anti-correlated.

Fourth, MPOA^CalcR^ responses exhibit a spatially biased organization that is independent of social experience. Specifically, pup-responsive neurons are largely absent from the most anterior MPOA, which is dominated by adult-responsive neurons. In contrast, the posterior MPOA contains both adult- and pup-responsive neurons, but with a higher proportion responding to pups. These spatial patterns suggest that pup-related inputs preferentially target medial-to-posterior MPOA regions, whereas adult-related inputs likely project broadly across the MPOA. This organization may explain why pup cues fail to activate non-parental neurons concentrated in the anterior MPOA, whereas parental neurons receive both adult- and pup-related sensory inputs.

### CalcR role in the emergence of parental behaviors

*CalcR* knockdown in the MPOA delayed the emergence of all aspects of parental behavior, and the extent of this delay positively correlated with the degree of knockdown. Pharmacological antagonism of CalcR-containing receptors also impaired pup retrieval in sensitized males, although the effect was weaker than that observed with *CalcR* knockdown. Neither manipulation affected pup-directed behaviors in non-sensitized males. Together, these results indicate that MPOA CalcR-containing receptors are primarily required for the emergence of parental behavior and, to a lesser extent, for the expression of specific behavioral components, while playing little to no role in non-parental males.

How do CalcR-containing receptors facilitate pup-sensitization–induced parental behavior? Addressing this question requires identifying which CalcR-containing receptor is involved. CalcR is complicated in that it participates in the formation of four receptors: the calcitonin receptor and the amylin receptors Amy1–Amy3, which are heterodimers composed of CalcR and RAMP1–3^54^. Several lines of evidence suggest that Amy1 is the most relevant receptor in the MPOA. First, single-nucleus and bulk RNA sequencing show that Ramp1 and Ramp2 are expressed in the MPOA^35^. Second, both AC187 and CGRP8–37 suppress parental behavior when injected into the MPOA, and among CalcR-containing receptors, Amy1 is the only receptor effectively antagonized by both drugs^58,68^. Third, calcitonin is not present in the central nervous system, making the calcitonin receptor an unlikely physiological target^60,61^.

If Amy1 is the relevant receptor, what is its endogenous ligand? Pharmacological studies in cultured cells suggest that Amy1 responds similarly to amylin and CGRP^54,58,68^. However, our recordings revealed that male MPOA^CalcR^ neurons respond quite differently to these two peptides. CGRP consistently increased input resistance without altering the resting membrane potential, whereas amylin had little effect on input resistance and only depolarized the cell at an unnaturally high concentration. It is worth noting that amylin expression is sexually dimorphic. Amylin level is much lower in males than in females throughout the hypothalamus^69^. In fact, our immunostaining failed to detect amylin in the male MPOA, while the signal is readily observed in females (Data not shown). Thus, although amylin was previously suggested to be the ligand of Calcr-containing receptors in the female MPOA^37^, we think this is unlikely the case in males. Even in females, knockout amylin induced no deficits in maternal behavior, suggesting that other ligands likely exist to activate CalcR-containing receptors^37^. Among the known Amy1 ligands^54,57^, CGRP appears to be the most likely one. Several MPOA-projecting brain regions contain CGRP-expressing cells, including locus coeruleus, posterior thalamus, and parabrachial nucleus^70–72^. Supporting CGRP as the relevant ligand, continuous CGRP infusion into the MPOA during pup sensitization accelerated the emergence of parental care.

Notably, CGRP is also a ligand for the CGRP receptor, a heterodimer composed of calcitonin receptor-like receptor (CALCRL) and Ramp1^54^. However, CGRP is unlikely to act on CalcR-expressing cells through CALCRL. Bulk RNA sequencing revealed negligible *Calcrl* expression in the MPOA, and single-nucleus RNA sequencing indicates that MPOA^CalcR^ cells do not express *Calcrl*^35^. Furthermore, the effect of CGRP on MPOA^Calcr^ cells can be blocked by AC 187, which does not antagonize the CGRP receptor^68^.

How does CGRP facilitate parental behavior through CalcR-containing receptors? Our data suggest that CGRP does not directly excite CalcR-expressing neurons but instead profoundly influences synaptic plasticity. When PA terminal activation occurs in the presence of CGRP, the PA→MPOA synapse undergoes strong potentiation, and this enhancement persists even after CGRP washout. PA neurons are naturally activated when naïve males interact with pups during sensitization. We therefore speculate that CGRP is released during pup sensitization and facilitates the potentiation of modestly activated PA→MPOA synapses carrying pup-related signals.

Importantly, although we focused on PA inputs, any inputs to MPOA^CalcR∩Vgat^ neurons that are activated during pup exposure could potentially be potentiated through a similar mechanism. Indeed, during pup sensitization, animals experience a barrage of pup-related olfactory and auditory cues, both of which have well-described roles in parental behaviors^73^. PA likely conveys primarily pup-related olfactory information, given its strong inputs from major olfactory relays such as the medial amygdala and posterolateral cortical amygdala^62^. Beyond PA, olfactory cues may also reach MPOA via the ventral premammillary nucleus, medial amygdala, and bed nucleus of the stria terminalis, while auditory cues, such as pup vocalizations, may reach the MPOA through the posterior intralaminar thalamic complex and the paraventricular nucleus of the hypothalamus^74–78^. Beyond CalcR, perturbation of Prlr^34,36^ and Esr1^79^ in the MPOA also impaired parental behaviors in both fathers and mothers. Notably, CalcR-positive neurons express Prlr, progesterone receptor (PR), and Esr1, suggesting that multiple hormones and neuropeptides may modulate parental behavior through the same neuronal population. Recently, Ammari et al. showed that during pregnancy, progesterone and estrogen induce synaptic plasticity and increased excitability of MPOA^Gal^ cells^53^. Although not tested here, MPOA^CalcR^ cells likely undergo similar hormone-dependent modulation given their strong PR and Esr1 expression. Thus, parental circuits may be strengthened through multiple signaling pathways, some hormone-dependent and others independent. Such diverse modulatory mechanisms may enable parental neurons to be activated “automatically” during parenthood while also allowing parental behavior to emerge gradually through repeated interactions with offspring in virgin animals. This experience-dependent plasticity may be particularly advantageous for species living in social groups^16^.

MPOA^CalcR^ cells are likely not the only MPOA population involved in parental behaviors. For example, the oxytocin system is an evolutionarily conserved regulator of parental care^2,3,80,81^. Disruption of OXTR signaling in the MPOA impairs parental behaviors, particularly under stressful conditions^82,83^. Interestingly, OXTR-expressing MPOA neurons show little overlap with MPOA^CalcR^ neurons^35^. How these two populations interact to coordinate parental behavior remains an important question for future studies.

It has been known since 1967 that pup sensitization can induce maternal behavior in virgin females independently of sex hormones^5^. However, the neural mechanisms underlying this phenomenon have remained unclear. Our study shows that virgin males, like females, can develop parental behavior after repeated pup exposure in the absence of sex hormones, and provides new insight into the neural mechanisms supporting sensitization-induced parental care. More broadly, the MPOA has increasingly been implicated in adult prosocial behaviors^84–88^. In this sense, the mechanisms described here may reflect a general principle by which familiarity breeds fondness.

## Methods

### Mice

All animal procedures were approved by the Institutional Animal Care and Use Committee (IACUC) of NYU Langone Health. Adult mice aged 8-36 weeks were used for all studies. Mice were housed at 20-22°C with 30–70% humidity under a 12 hr light-dark cycle (10 a.m. to 10 p.m. dark) with water and food available ad libitum. All animals were group-housed until adulthood and single-housed after surgery. All experiments were performed during the dark phase of the daily light/dark cycle. Calcr-Cre mice were originally provided by Dr. Martin Myers (now Jackson stock no. 0307028) ^40^. Esr1-2A-Cre mice were originally provided by Dr. David Anderson (now Jackson stock no. 017911). Vgat-Flp and Vglut2-Flp mice were purchased from Jackson Laboratory (stock no. 029591 and 030212). Calcr-Cre × Vgat-Flp and Calcr-Cre × Vglut2-Flp were acquired by crossing Calcr-Cre mice with Vgat-Flp or Vglut2-Flp mice, respectively. Wild-type C57BL/6 mice were purchased from Charles River or bred in-house and were used for behavior, shRNA, and pharmacology experiments. P1-P6 C57BL/6 pups were used for behavioral experiments and were bred in-house. Castrated wild-type C57BL/6 male mice (8-10 weeks) were purchased from Charles River. Adult (>8 weeks), wild-type Balb/c male and C57BL/6J female mice used as stimulus animals were bred in-house.

### Viruses and stereotaxic surgery

AAV1-hSyn-Flex-GCaMP6s (1.6 × 10^13^ vg per ml), AAV8-hSyn-DIO-hM4Di-mCherry (1.8 × 10^13^ vg per ml), AAV2-hSyn-DIO-mCherry (1.9 × 10^13^ vg per ml), AAV8-nEF-ConFon-ChRmine-oScarlet (1.9 × 10^13^ vg per ml), AAV8-EF1α-ConFon-oScarlet (2.2 × 10^13^ vg per ml), and AAV5-Syn-Chronos-GFP (5.3 × 10^12^ vg per ml) were purchased from Addgene. AAV2-EF1a-DIO-hChR2-EYFP (4.0 × 10^12^ vg per ml), AAV2-EF1a-DIO-hChR2-mCherry (3.1 × 10^12^ vg per ml), and AAV5-hSyn-ConFon-eYFP (4.8 × 10^12^ vg per ml) were purchased from the University of North Carolina vector core. HSV-hEF1α-LS1L-GCaMP6f (1.0 × 10^9^ vg per ml) was purchased from the Massachusetts General Hospital Neuroscience vector core. AAV8-hSyn-CalcR-shRNA-Clover3 (3.11 × 10^14^ vg per ml) and AAV8-hSyn-CalcR-scrambledmRNA-Clover3 (1.02 × 10^15^ vg per ml) were generated by Drs. Qian Wang and Xiaoke Chen. Viruses containing *Calcr* shRNA and scrambled shRNA were diluted 1:10 in 0.9% saline prior to virus injection.

Mice (8-24 weeks old) were anesthetized with isoflurane (3.5% induction and 1-2% maintenance vaporized in oxygen delivered at a rate of 2 L/min) and positioned on a stereotaxic apparatus (Kopf Instruments Model 1900). Anesthesia was monitored by checking breathing rate and foot-pinch reflex throughout the surgery. Viruses were injected into the brain using a glass capillary attached to a nanoinjector (World Precision Instruments, Nanoliter 2000). In all cases, viruses were injected at a rate of 30 nL per minute, and capillaries were left in place for a minimum of 5 minutes after virus injection before being slowly retracted from the brain.

For two-photon imaging of MPOA^CalcR^ cells, 200 nL AAV1-hSyn-DIO-GCaMP6s was unilaterally injected into the MPOA (AP: 0.0 mm, ML: -0.31 mm, DV: -4.9 mm) of heterozygous Calcr-Cre male mice. After virus injection, the tissue above the MPOA was aspirated using a blunt-end 23-gauge needle. A 600-μm GRIN lens (Inscopix, 1050-004597) attached to a 4mm cover glass (Warner Instruments, CS-4R) using optic UV glue (Norland Products, NOA87) was implanted 250 μm above the virus injection site and secured to the skull using dental cement (C&B Metabond, S380). After GRIN lens implantation, a custom 3D-printed head plate was secured onto the skull using dental cement (C&B Metabond, S380). We began imaging experiments 4-7 weeks after surgery when GCaMP-expressing cells were clearly visible.

We attempted to perform two-photon imaging of MPOA^CalcR∩VGAT^ cells by implanting a 600-μm GRIN lens into the MPOA of Calcr-Cre × Vgat-Flp × Ai195 male mice. However, only 1 of 7 implanted mice exhibited stable GCaMP signals during recording. Testing multiple AAVs expressing ConFon-GCaMP likewise failed to yield sufficient, stable expression for two-photon imaging.

For chemogenetic inhibition of MPOA^CalcR^ cells, 100 nL of AAV8-EF1α-DIO-hM4Di-mCherry or AAV8-hSyn-DIO-mCherry was bilaterally injected into the MPOA (AP: 0.0 mm, ML: ±0.31 mm, DV: -4.9 mm) of heterozygous Calcr-Cre male mice. Animals were allowed to recover for at least 3 weeks prior to testing.

For optogenetic activation of MPOA^CalcR^ subpopulations, 90 nL of AAV8-nEF-ConFon-ChRmine-oScarlet was bilaterally injected into the MPOA of Calcr-Cre × Vgat-Flp or Calcr-Cre × Vglut2-Flp male mice. For optogenetic activation of MPOA^CalcR^ cells, AAV2-EF1a-DIO-hChR2-EYFP/mCherry was bilaterally injected into the MPOA of Calcr-Cre male mice. For all experiments, two 200-μm optical fiber assemblies (RWD Life Science, R-FOC-L200C-0.39NA, L: 5mm) were implanted at 10° angles 200 μm above the injection sites. Optical fibers were secured on the skull using dental cement (C&B Metabond, S380), and a 3D-printed head fixation ring ^89^ was cemented to the skull. Animals were allowed to recover for 3-5 weeks prior to testing.

To knock down *Calcr* expression, 90 nL of AAV8-hSyn-Calcr-shRNA-Clover3 or AAV8-hSyn-Calcr-scrambledshRNA-Clover3 was bilaterally injected into the MPOA of wild-type C7BL/6 male mice. The target sequences for knockdown experiments were as follows: *Calcr* shRNA: 5’-GATGAGGTTCCTTCTCGTGAA-3’; scrambled shRNA: 5’-GGTTTCCATTGCGCTTAGAGA-3’. Animals were allowed to recover for 3-5 weeks after surgery.

To block CalcR signaling in MPOA, we implanted bilateral cannulae (Plastics One, center-to-center distance: 0.7 mm) 400 μm above the MPOA of wild-type C57BL/6 male mice. The cannulae were secured to the skull using dental cement (C&B Metabond, S380), and a 3D-printed head fixation ring^89^ was cemented to the skull. Animals were allowed to recover for 1-2 weeks after surgery.

To record Ca^2+^ activity from PA^Esr1^ cells projecting to the MPOA, we injected 100 nL of HSV-hEF1α-LS1L-GCaMP6f into one side of the MPOA. A 400-μm fiber optic assembly (RWD Life Science, R-FOC-L400C-0.5NA, L: 5.5 mm) was implanted 100 μm above the ipsilateral PA (AP: −2.3 mm; ML: -2.4 mm; DV: −4.7 mm). The optical fiber was secured using dental cement (C&B Metabond, S380), and a 3D-printed head fixation ring was secured to the skull. All experiments began 2-3 weeks after surgery.

For probing PA inputs to MPOA in acute brain slices, 100 nL of AAV5-Syn-Chronos was bilaterally injected into the PA, and 200 nL of AAV8-nEF-ConFon-oScarlet was bilaterally injected into the MPOA of Calcr-Cre × Vgat-Flp male mice.

To record electrophysiological properties of MPOA^CalcR∩VGAT^ cells, 200 nL of AAV8-nEF-ConFon-oScarlet was bilaterally injected into the MPOA of Calcr-Cre × Vgat-Flp mice.

To record electrophysiological properties of MPOA^CalcR∩VGAT^ after *Calcr* knockdown, 200 nL of AAV8-nEF-ConFon-oScarlet mixed 1:1 with AAV8-Syn-Calcr-shRNA-Clover3 was injected unilaterally. On the other side of the MPOA, 200 nl of AAV8-nEF-ConFon-oScarlet mixed 1:1 with AAV8-Syn-Calcr-scrambledshRNA-Clover3 was injected. The brains were used for recording 3-5 weeks later.

To continuously infuse CGRP into the MPOA, we implanted a unilateral cannula connected to a 100 μL osmotic pump (Azlet, 1002) pre-filled with 0.9% saline or 200 μM CGRP (Tocris, 1161). A small incision was made in the back to implant the pump subcutaneously. The cannula and catheter tubing were secured using dental cement (C&B Metabond, S380). Experiments began 4 days after surgery.

To trace the projections of MPOA^CalcR∩VGAT^ cells, we injected 80 nL of AAV8-nEF-ConFon-oScarlet or AAV5-hSyn-ConFon-EYFP into the MPOA. Animals were sacrificed 4 weeks later for histological analysis.

### Behavioral assays and analysis

All behaviors in all experiments were recorded from the side and top of the cage using two synchronized cameras (Basler, acA640-120gc) using commercial video acquisition software (StreamPix 8, Norpix, RRID: SCR_015774) in a semi-dark room with infrared illumination at a frame rate of 25 frames per second. Behaviors were manually annotated on a frame-by-frame basis using custom MATLAB software (https://github.com/pdollar/toolbox) ^90^.

During pup interaction tests, we manually annotated the following behaviors: approach, attack, investigation, retrieve, groom pup, crouching, and nest building. Approach is defined as the first step toward the pup from the farthest point away from the pup. Investigation is defined as nose contact with the pup’s body. Retrieval starts from the moment the male picks up the pup from the floor of the cage and ends when it drops the pup in or around the nest. Grooming the pup is defined as direct mouth-to-body contact with the pup, associated with rhythmic up and down head movements by the male and displacement of the pup. Crouching is characterized by the male hovering above the pups without obvious movements. Nest building is defined as the male collecting and repositioning the nesting material with his mouth to form a mass. The expression of infanticide would override the classification of a mouse as ‘parental.’ We found that the expression of pup retrieval was almost mutually exclusive with the expression of infanticide (attack). In one case (mouse 3, day 6 in **Fig. 1c, d**), a mouse showed infanticide and retrieval on the same day. In this case, this mouse was classified as infanticidal. No other mice in the course of our study showed both infanticide and pup retrieval on the same day.

Pup interaction tests were performed by introducing pups one by one into the males’ home cages. First, males were habituated to the removal of the cage top for at least 20 minutes. Then, the first pup was introduced into its cage. As some animals attacked pups with a relatively long latency (>5 minutes), we allowed animals to interact with the pup for 10 minutes on this first trial. If an animal attacked the pup, we ended the experiment and quantified the attack latency. For each subsequent trial, we introduced each pup one at a time at 2-minute intervals and continued to record behavioral videos for 10-20 minutes after introducing the last pup. Up to 6 pups were left in the cage after being retrieved. The proportion of retrieval trials was calculated as the number of trials during which the introduced pup was retrieved to the nest within 2 minutes among all trials. For quantification of other behaviors, we scored the 10 minutes after the last pup introduction.

This large number of retrieval trials permitted a scalable metric for pup retrieval, wherein we could resolve quantitative differences in parenting performance across the sensitization process and under different experimental manipulations. However, this repeated-trial design constrained the ability to quantify other parental behaviors during the retrieval phase, as the cage was repeatedly disturbed to introduce new pups. To address this, we recorded each animal undisturbed for an additional 10 minutes at the end of the session to quantify parental behaviors other than pup retrieval.

Pup sensitization was done by placing 2-3 pups (P1-6) in a wire mesh ball (2.1-inch tea strainers, Amazon) and co-housing them with males in their home cages. Pups were removed from the wire mesh ball, returned to their mothers to nurse, and immediately replaced with new pups every 6-8 hours. Control animals were sensitized to pup-sized objects (4-40 stainless steel cap screws, 3/8”, Thorlabs SH4S0384-40), housed in the same wire mesh balls.

For tests with adult male intruders, one group-housed BALB/c adult (>8 weeks) male was introduced into the animals’ home cages. Investigation is defined as nose-to-nose, nose-to-trunk, or nose-to-anogenital contact. Attack is classified by lunging and biting movements.

For tests with adult female intruders, one group-housed C57BL/6 adult (>8 weeks) female was introduced into the animals’ home cages. Investigation is defined as nose-to-nose, nose-to-trunk, or nose-to-anogenital contact. Mounting is defined as the male extending his forelimbs to grasp the back of the female to position himself on top of her.

### Immunohistochemistry

Mice were anesthetized and transcardially perfused with 1 × PBS, followed by 20 ml of 4% paraformaldehyde (PFA). Brains were fixed in 4% PFA for 24 hours at 4°C and dehydrated in 30% sucrose overnight. Brains were then embedded in O.C.T. compound (Sakura, 4583), flash frozen, and cut into 50-μm sections using a cryostat (Leica, CM1950). Every third section was used for immunohistochemistry. Free-floating brain slices were rinsed three times with 1 x PBS (10 min per wash) and once with PBST (0.1% Trition X-100 in PBS, 30 minutes) at room temperature. Slices were then incubated in a blocking solution (10% normal donkey serum (Jackson ImmunoResearch, 017-000-121) in PBS) at room temperature for 1 hour. Primary antibodies (guinea pig anti-Fos, 1:1000 dilution, Synaptic Systems, 226005; rabbit anti-CalcR, 1:1000 dilution, Abcam ab11042) were diluted in PBST with 3% normal donkey serum and incubated overnight at 4°C. We then washed the brain slices with PBST (10 minutes) three times and incubated them with secondary antibodies (Cy3-goat anti-guinea pig, 1:500 dilution, Jackson ImmunoResearch, 706-165-148 or 488-goat anti-guinea pig, 1:500 dilution, Jackson ImmunoResearch, 706-545-148; Cy5-donkey anti-rabbit, 1:500 dilution, Jackson ImmunoResearch, 7110175-152) for 2 hours at room temperature. Slices were then washed three times with PBST (10 minutes) and once with 1 × PBS (10 min) and mounted onto slides (Thermo Fisher Scientific, 12-550-15). Slides were coverslipped with 50% glycerol-containing DAPI (Invitrogen, 00-4959-52). We imaged slides using a slide scanner microscope (Olympus, VS120).

### Histological analysis

For c-Fos and CalcR quantification (**Fig. 1p, q, and Fig. 5c**), we identified the MPOA based on the mouse brain atlas^91^ and counted cells manually using ImageJ. We summed the number of counted cells across three 50 µm sections of the MPOA (Bregma levels 0.14mm, -0.10 mm, and -0.22 mm).

To quantify the projection density from MPOA^CalcR∩VGAT^ cells, QuPath was used^92^. We selected a boxed area (200 × 200 μm^2^) in each region containing fibers and calculated the average pixel intensity as *F_raw_.* On the same image, a boxed area of the same size but in a region containing no fiber terminals was selected for calculating the background intensity (*F_background_*). We calculated *F_signal_* as *F_raw_-F_background_*. *F_signal_* was normalized by the maximum *F_signal_* across all brain regions within each animal. The normalized *F_signal_* is reported in **Extended Data Fig. 6**.

### Immediate early gene mapping

All animals were wild-type C57BL/6 males, assigned randomly to experimental or control conditions. Experimental animals were pup-sensitized for 7 days and tested for robust paternal behavior (retrieval of at least 7 of 10 pups) the day before c-Fos induction. Control animals were single-housed and tested for behavior toward pups the day before c-Fos induction. We introduced 3 pups in wire mesh balls into the males’ home cages for 20 minutes and sacrificed animals 60 minutes after removal of the pups. Histology for pup-sensitized and single-housed animals was performed side-by-side in 3 cohorts.

### Two-photon imaging

To record Ca^2+^ signals from individual MPOA^CalcR^ neurons in freely-moving mice, we used a miniature head-mounted two-photon microscope, as previously described^49,93^ (Transcend Vivoscope, FHIRM-TPM). GCaMP was imaged with an excitation wavelength of 920 nm from a Mai Tai laser (Spectra-Physics, Mai Tai HP 1040s). The laser power was optimized for each animal and ranged from ∼30-50 mW, similar to previous studies^94^.

Starting 4 weeks after surgery, mice were checked for GCaMP expression by being head-fixed and placed on a running wheel under a widefield microscope (Thorlabs, SFMGFP2). Once sufficient expression was reached (4-7 weeks after surgery), we attached the two-photon miniscope to a baseplate using M1 screws to identify the optimal imaging field. We then secured the baseplate to the mouse’s headplate using dental cement (C&B Metabond, S380). We let the dental cement dry completely and then removed the two-photon microscope from the implanted baseplate. We attached a screw to the baseplate (4-40 stainless steel cap screws, 3/8”, Thorlabs Thorlabs SH4S0384-40) to habituate the mouse to the weight of the microscope and returned the mouse to its home cage.

On the days of recording, we attached the two-photon miniscope to the baseplate and acquired images at 4.8 frames per second. Behavior videos were simultaneously recorded from the top and side at 25 frames per second. Animals were habituated to the miniscope in their home cage for at least 15 minutes before image acquisition. On each recording day, we recorded 5-10 minutes of baseline activity when the animal was alone in its home cage. We then introduced pups one by one into the mouse’s cage and continued to record neural activity and behavior for approximately 20 minutes after introducing the last pup. For most sessions, we also introduced adult male and female intruders after the pup presentation. When adult intruders were included in the experiment, we included an inter-trial interval of at least 5 minutes between intruder types. The stimulus presentation order was kept consistent across days.

TIF stacks from two-photon imaging experiments were pre-processed using Suite2p (https://github.com/MouseLand/suite2p). We performed motion correction separately for each day of imaging. We defined regions of interest corresponding to cell bodies semi-automatically using an activity-based algorithm, along with manual curation, and extracted fluorescence traces for each region of interest.

We tracked cells across days by registering average projection images for each session and computing the spatial overlap of cell masks using the MATLAB program ROIMatchPub (https://github.com/ransona/ROIMatchPub). We semi-automatically classified neurons as matches if 50% of the pixels in the cell masks overlapped, with additional manual curation to inspect the cell shape and size and the local environment (e.g., neighboring cells and neurites) surrounding the cells.

### Two-photon imaging analysis

We performed all further analysis on the extracted fluorescence traces using custom-written MATLAB scripts. The MATLAB function ‘msbackadj’ with a moving window of 1/8 of the total recording duration was first applied to the fluorescence trace to obtain the moment-to-moment baseline signal (F_baseline_). The instantaneous ΔF/F was calculated as (F_raw_ − F_baseline_) / F_baseline_. The z-score of the ΔF/F signal was obtained using the MATLAB function ‘zscore’ over the entire duration of the recording. The average response during each behavior was calculated as the average z-scored ΔF/F of all frames when the behavior occurred.

We performed principal components analysis on the matrix of averaged peri-event time histograms (from -10 to +10 seconds surrounding behavior onset) in all cells using the MATLAB function ‘pca’ (**Fig. 3g, h, Extended Data Figure 8**). We performed k-means clustering on the first PCs that could explain over 90% of the variability in the data. We determined the number of clusters using the point at which the within-cluster sum of squares slope changes sharply (elbow method).

To determine significantly responsive cells during parental behaviors (**Fig. 3e**), we bootstrapped the average behavior-evoked z-scored ΔF/F due to a low number of trials of some behaviors (fewer than 6 events). We resampled the distributions 1,000 times and performed a permutation test against the baseline (no interaction epochs). P-values were adjusted using the false discovery rate (FDR) using the Benjamini & Hochberg method.

Pup, male, and female investigatory events all occurred more than 6 trials per session. Thus, we performed a one-sample t-test on the averaged behavior-evoked responses to identify cells that were significantly responsive during these behaviors (**Fig. 4d, e, Extended Data Fig. 12b, c, Extended Data Fig. 13a-g**). P-values were adjusted for multiple comparisons using the false discovery rate (FDR) with the Benjamini–Hochberg procedure. For all behaviors, cells with corrected p-values (q-values) less than 0.01 and whose average z-scored ΔF/F response exceeded 0.5 were classified as significantly responsive.

The preferred tuning of each cell was determined solely based on significant responses (**Fig. 4i, Extended Data Fig. 12g**). For cells that were significantly responsive to only one stimulus, that stimulus is the preferred stimulus. For cells that were significantly responsive to multiple stimuli, we considered the social stimulus that elicited the largest response as the preferred stimulus.

To calculate the discriminability of MPOA^CalcR^ cells to social stimuli, we constructed histograms of z-scored ΔF/F distributions for each cell’s responses to each social stimulus and then generated an ROC curve based on the histograms for each pair of stimuli (**Extended Data Fig. 11**). We subtracted 0.5 from the area under the ROC curve (auROC) and took the absolute value as the discriminability index (**Extended Data Fig. 11c, f**). A discriminability index close to 0 indicates chance levels, wherein a cell cannot discriminate between two stimuli. The selectivity index was calculated as the auROC, reflecting a cell’s preference for one stimulus over another. A selectivity index of 0 or 1 indicates sole preference for one stimulus, while a selectivity index of 0.5 indicates no selectivity (**Extended Data Fig. 13h-j**).

To determine if MPOA^CalcR^ neuron activity could decode individual parental behaviors (**Fig. 3o**), we trained a separate linear support vector machine (SVM) decoder for each behavior in each animal to predict whether the behavior was occurring or not, using similar methods to those published previously^86^. Manual annotations of each behavior bout were used as training labels for each behavior. We took the mean z-scored ΔF/F from all neurons during each bout, yielding one population activity vector per bout. We did not subtract any baseline period from the z-scored ΔF/F. A random sample of bouts in which the animal was not interacting with pups was labeled as negative training data. This negative training data was constrained to times when pups were in the cage. We randomly downsampled the negative data to ensure equal numbers of positive and negative data in our training and test datasets. A linear SVM was trained and evaluated using leave-one-out cross-validation (LOO-CV), in which each bout was held out for testing, and the remaining bouts were used for training. Across each of these validation folds, a prediction score was generated. We compared all prediction scores to the ground truth labels using an ROC curve, and the area under the ROC curve was taken as the metric of decoder performance. Decoding was repeated 20 times for each behavior, and decoder performance was reported as the average auROC. Shuffling was performed by randomly permuting the behavior and ‘no interaction’ labels. Shuffling was repeated 1000 times. Shuffle iterations were averaged for each animal to calculate a shuffled decoder accuracy.

To classify different pup-directed behaviors from MPOA^CalcR^ neuron activity (**Fig. 3p**), a multi-class SVM (MATLAB’s fitcecoc function) was used. Similar to the binary decoder, we took the mean z-scored ΔF/F from all neurons during each behavior bout to produce a population activity vector per bout. Using LOO-CV, a single bout was held out as a test sample, and the classifier was trained on the remaining bouts. To account for class imbalance across behaviors, we used a class weighting approach. Specifically, misclassification costs were weighted inversely to class frequency in the training data. Class weights were proportional to the square root of the inverse class frequency, normalized across classes, and capped at a maximum of 2x to prevent extreme weighting. Weights were incorporated into a cost matrix that penalized misclassification of underrepresented behaviors during training. We calculated the per-class F1 score as 2 (precision × recall)/(precision + recall), where precision is the fraction of predicted positives that are true positives ((true positives)/(true positives + false positives)) and recall is the fraction of true positives that are correctly predicted ((true positives)/(true positives + false negatives)) for each behavior. Decoding was repeated 20 times, and the F1 scores were averaged across all iterations. A confusion matrix was constructed and normalized by the number of true samples per class. Shuffling was performed by randomly permuting the behavior labels. Shuffling was repeated 100 times, and shuffle iterations were averaged for each animal and each behavior to calculate a shuffled decoder accuracy.

To test if MPOA^CalcR^ neuron activity could distinguish pairs of pup-directed behaviors (**Fig. 3q**), we trained separate linear SVMs for each behavior pair in each animal. The mean z-scored ΔF/F from all neurons during each behavior bout was used to produce a population activity vector per bout. We again used LOO-CV to train and assess decoder performance. We used the same class weighting approach as the multi-class decoder to balance across behaviors. For each validation fold, we generated a prediction score and compared all prediction scores to the ground truth labels using an ROC curve. The area under the ROC curve was taken as the metric of decoder performance. Decoding was repeated 20 times for each behavior and decoder performance was reported as the average auROC.

### Chemogenetic-mediated silencing

For chemogenetic inhibition experiments, naïve male mice were used. After recovery from surgery, animals were tested for spontaneous paternal behavior. If animals showed paternal behavior (retrieval of 1 or more pups), they were excluded from the experiment.

For acute chemogenetic experiments, animals were sensitized to pups for a minimum of 5 days to robust paternal behavior (retrieval of at least 7 of 10 pups). Mice were injected with saline or clozapine-n-oxide (CNO; Enzo Life Sciences BML-NS105-0025), 1mg/kg on sequential days. Thirty minutes after the injection, behavior experiments began. We ran the pup interaction test on each day. To probe the animals’ behavior toward adults, one adult male and one adult female were introduced to the animals’ cage for 10 minutes, with 10 minutes in between each intruder type. Females were not screened for estrous stage or sexual receptivity.

For chronic chemogenetic experiments, animals were first tested in the pup interaction test (day 0). Immediately following this test, we replaced the standard drinking water with water containing CNO (Enzo Life Sciences BML-NS105-0025) at a concentration of 25 mg/L, as described previously^2,95^. We added sucrose to the water to an effective concentration of 1%. On average, we observed that the mice drank approximately 6 mL of water per day, leading to an effective CNO dose of 5 mg/kg/day. Approximately 6-8 hours after water replacement, we began pup sensitization. We continued pup sensitization and performed the next pup interaction test approximately 24 hours following the start of sensitization. We repeated the sensitization and pup interaction test daily for one week (days 1-7). We refreshed the CNO water daily. After the pup interaction test on day 7, we replaced the CNO water with standard drinking water and ceased pup sensitization. For 3 subsequent days (days 8-10), we performed the pup interaction test daily, with approximately 24 hours between tests.

### Optogenetic stimulation

For optogenetic stimulation experiments, naïve male mice were used. After recovery from surgery, animals were tested for spontaneous paternal behavior. If animals showed paternal behavior (retrieval of 1 or more pups), they were excluded from the experiment.

To stimulate ChRmine-expressing cells, we used a 589 nm laser (Shanghai Dream Laser) or 545 nm LED (Prizmatix). For ChR2-expressing cell activation, we used a 473 nm laser (Shanghai Dream Laser). In both cases, light pulses were delivered by TTL signals generated using an RP2 or RZ5 processor (Tucker Davis Technologies).

On days of testing, mice were head-fixed, and optical fibers were attached to patch cords using a zirconia split sleeve (Thorlabs, ADAL1-5). To test the animals’ behavior towards pups, we introduced one pup (P1-6) at a time into the animals’ home cage. To account for the potential gradual or long-lasting change in behavior caused by stimulation, as we previously observed after activating MPOA^Esr1^ cells in females ^30^, we ran all sham trials prior to any light trials. We delivered 5-10 sham stimulation trials (0 mW, 20 s) and then 7-13 light stimulation trials (1-3 mW, 20 ms, 20 Hz, 20 s). In all cases, the sham and light onset were triggered when the animal was on the opposite side of the cage from the pup. The inter-trial interval was approximately 20 s, although it could be longer depending on the animals’ behavior and location in the cage. To test behavior toward adult males and females, one adult BALB/c male or one adult C57BL/6 female was introduced. Sham and light stimulation were interleaved, with an inter-trial interval of approximately 20 seconds. Stimulation began when the animal first contacted the intruder.

For the real-time place preference test, animals were habituated to a custom two-chamber arena for 10 minutes and could explore each chamber freely. They then received sham stimulation when they entered the predesignated stimulation chamber for a 10-minute test. In the subsequent 10 minutes, animals received light stimulation (2 mW, 20 ms, 20 Hz) when they entered the stimulation chamber; stimulation ceased when animals moved out of the stimulation chamber. Sham and stimulation chambers were counterbalanced across mice. Video was recorded using a top-view camera, and the location of the animal was tracked and analyzed using custom-written MATLAB code (https://github.com/pdollar/toolbox)^90^.

### shRNA-mediated gene knockdown

For gene knockdown experiments, naïve male mice were used. Three to five weeks after surgery, animals were tested for spontaneous paternal behavior. If animals showed paternal behavior (retrieval of 1 or more pups), they were excluded from the experiment. Pup sensitization and pup interaction tests were performed as previously described. After testing, animals were sacrificed, and immunohistochemistry was performed to confirm *Calcr* knockdown.

### Pharmacology

For acute pharmacology experiments, all animals were allowed to recover for one to two weeks following surgery. In experiments in naïve animals, we injected 300 nL per side into the MPOA with 600 μM AC187 (Tocris, 3419), 600 μM CGRP 8-37 (Tocris, 1169), 200 μM CGRP (Tocris, 1161), or 0.9% sterile saline using a syringe (Hamilton, 65457-2) when animals were head-fixed on a running wheel. For studies of CalcR antagonism in sensitized males, we sensitized animals for at least 5 days until they showed robust paternal behavior (retrieval of at least 7 of 10 pups). To antagonize CalcR-associated receptors, we injected 300 nL per side into the MPOA with 600 μM AC187 (Tocris, 3419) or 600 μM CGRP 8-37 (Tocris, 1169) or 0.9% sterile saline using a syringe (Hamilton, 65457-2) when animals were head-fixed on a running wheel. After drug infusion, animals were returned to their home cage for 20 minutes prior to starting the pup interaction test. The order of saline and antagonist injections was counterbalanced across animals.

For continuous infusion of CGRP, naïve male mice were used. Animals were pre-screened for infanticidal behavior. Experiments began 4 days after surgical implantation of the osmotic pump (Azlet, 1002). Pup sensitization and pup interaction tests were performed as previously described.

### Fiber photometry

Fiber photometry was performed as described previously^30,46,62^. Briefly, 390 Hz sinusoidal blue LED light (30 μW; LED light: M470F1; LED driver: LEDD1B; Thorlabs) was band-pass filtered (passing band: 472 ± 15 nm, FF02-472/30-25, Semrock) and delivered to the brain to excite GCaMP6f. The emission light traveled back through the same optic fiber and was band-pass filtered (passing bands: 535 ± 25 nm, FF01-535/505, Semrock), passed through an adjustable zooming lens (Thorlabs, SM1NR01 and Edmund optics, 62-561), detected by a Femtowatt Silicon Photoreceiver (Newport, 2151), and recorded using a real-time processor (RP5, Tucker Davis Technologies). The envelope of the 390 Hz signals reflected the intensity of the GCaMP and was extracted in real-time using a custom TDT program. The signal was low-pass filtered with a cut-off frequency of 10 Hz. The blue LED was adjusted so that the light intensity at the tip of the patch cord was 30 μW.

For PA^Esr1◊^ ^MPOA^ cell fiber photometry, mice were not pre-screened for infanticidal or paternal behavior. Before recording, mice were head-fixed, and optic fibers were attached to a patch cord using a zirconia split sleeve (Thorlabs, ADAL1-5). Mice were recorded for 10 minutes alone in their home cage prior to pup introduction. Pups (P1-P6) were introduced one by one, each for approximately 2 minutes, for up to 10 trials. If the animal attacked a pup, we removed the pup and reintroduced a pup to start a new trial. After a 5-minute inter-trial interval, we introduced 3 pups housed in a wire mesh ball for 10 minutes. None of the mice showed paternal behavior during testing. Animals that showed correct fiber placement and GCaMP expression were used for final analysis.

The MATLAB function ‘msbackadj’ with a moving window of 1/8 of the total recording duration was first applied to the fluorescence trace to obtain the moment-to-moment baseline signal (F_baseline_). The instantaneous ΔF/F was calculated as (F_raw_ − F_baseline_) / F_baseline_. The z-score of the ΔF/F signal was obtained using the MATLAB function ‘zscore’ over the entire duration of the recording. PETHs were constructed by aligning the ΔF/F signal for each trial to the onset of each behavior and z-scored. Behavior-evoked responses were calculated by subtracting the average pre-behavior activity (z-scored ΔF/F values -5 to -3 seconds before behavior onset) from the average post-behavior activity (z-scored ΔF/F values 0 to 2 seconds after behavior onset) of each animal.

### Brain slice electrophysiology

To prepare brain slices for whole-cell recording, mice were anesthetized with isoflurane, and brains were quickly removed and immersed in an ice-cold cutting solution for 1-2 min. The cutting solution composition consisted of the following in mM: 110 choline chloride, 25 NaHCO_3_, 2.5 KCl, 7 MgCl_2_, 0.5 CaCl_2_, 1.25 NaH_2_PO_4_, 25 glucose, 11.6 ascorbic acid, and 3.1 pyruvic acid. Coronal sections of the MPOA region with a thickness of 275 μm were prepared using a Leica VT1200s vibratome and then placed in oxygenated (95% O_2_ and 5% CO_2_) and pre-heated (32-34 °C) ACSF solution containing 125 NaCl, 2.5 KCl, 1.25 NaH_2_PO_4_, 25 NaHCO_3_, 1 MgCl_2_, 2 CaCl_2_, and 11 glucose for a 30-minute incubation period. Subsequently, the sections were transitioned to room temperature and continuously oxygenated until further use. The intracellular solution for the voltage clamp recording contained (in mM): 135 CsMeSO3, 10 HEPES, 1 EGTA, 3.3 QX-314 (chloride salt), 4 Mg-ATP, 0.3 Na-GTP, and 8 sodium phosphocreatine (pH 7.3 adjusted with CsOH). The intracellular solution for the current clamp recordings was composed of the following in mM: 145 potassium gluconate, 2 MgCl_2_, 2 Na_2_ATP, 10 HEPES, 0.2 EGTA (286 mOsm, pH 7.2). Signals were recorded using a MultiClamp 700B amplifier (Molecular Devices) and Clampex 11.0 software (Axon Instruments) and digitized at 20 kHz using Digidata 1550B (Axon Instruments). Following data acquisition, analyses were performed using Clampfit software (Molecular Devices).

Four weeks after surgery, brain slices were obtained for whole-cell recordings from MPOA^CalcR∩VGAT^ cells, identified by oScarlet expression. For experiments probing the synaptic connection from PA to MPOA, Chronos-GFP expression in the PA was assessed using an Olympus ×40 water-immersion objective with a GFP filter. For experiments characterizing the effects of *Calcr* shRNA (or scrambled shRNA) on the physiological properties of MPOA^CalcR∩VGAT^ cells, we recorded from cells co-expressing oScarlet and GFP.

We performed whole-cell current-clamp recordings to determine the intrinsic excitability of MPOA^CalcR∩VGAT^ cells. We recorded the voltage changes of the cells in response to a series of 500ms current steps from 0 to 250 pA in 10 pA increments. In addition, the intrinsic properties of each cell, such as rheobase, resting membrane potential, and input resistance, were characterized. The rheobase was defined as the minimum current required to induce the first action potential. The resting membrane potential was measured directly after rupturing the cell membrane with 0 pA holding current. The input resistance was read from the membrane test window of the software after breaking into the membrane (Clampex 11.0 software). The maximum action potential number was the highest number of action potentials fired in response to any current injection.

To record the synaptic properties of MPOA^CalcR∩VGAT^ cells, we performed whole-cell voltage-clamp recordings. The cell membrane potential was held at -70mV to record mEPSCs and 0mV to record mIPSCs. For mEPSC and mIPSC recordings, we bath-applied 1 μM tetrodotoxin citrate (TTX, Tocris).

To investigate the synaptic connectivity between PA and MPOA^CalcR∩VGAT^ cells, we performed whole-cell voltage-clamp recordings of MPOA^CalcR∩VGAT^ cells (oScarlet positive) and then delivered 1-ms pulses of 470-nm full-field illumination (pE-300white; CoolLED) at 30 s intervals to activate Chronos-expressing PA terminals. oEPSCs were recorded while holding the membrane potential of recorded neurons at -70 mV.

To measure excitatory synaptic responses of recorded MPOA^CalcR∩VGAT^ cells, we performed whole-cell current-clamp recordings. We injected a small positive or negative current to keep the cell membrane potential around −70 mV and delivered a 1 ms 470-nm full-field light pulse every 10 s (0.1 Hz) (pE-300white; CoolLED). After 2-3 min of initial probing, we then delivered 3 trains of 20 Hz, 5 ms light pulses with 25 s per train and 5 s in between. During light pulse train delivery, the cells were not injected with any positive or negative current. After light-train delivery, we again injected negative or positive current to maintain the membrane potential around −70 mV, probed light-evoked EPSPs for 5 min, and then bath-perfused CGRP (200 nM) or amylin (200 or 800 nM). After 10 min of CGRP or amylin perfusion, we probed optically-evoked EPSPs (oEPSPs) for 2-3 min. Then, we delivered a second set of light pulse trains and probed the oEPSPs for 5 min, and then CGRP or amylin was washed out with ACSF while the oEPSPs were continuously probed at 0.1 Hz for 10 min. In an alternative protocol, oEPSPs were probed before and after the co-perfusion of 200 nM CGRP and 600 nM antagonist (AC 187 or CGRP 8-37) for 10 minutes. Following the second stimulation with light, the light-evoked EPSPs were probed for 5 minutes, and then CGRP and the antagonist were washed out with ACSF, while continuous probing of oEPSPs was carried out for an additional 10 minutes.

### Statistics

We did not use statistical methods to predetermine sample sizes but used similar sample sizes to those reported in the literature^30,46,62^. Each experiment was conducted using 2-5 cohorts of animals. We did not observe differences across cohorts, and all animals were combined for final analysis. All statistical analyses were performed using MATLAB (versions 2019b and 2023a) or GraphPad Prism (version 10) software. Statistical analyses were two-tailed. Parametric tests, including paired and unpaired t-tests and one- and two-way ANOVAs, were used if all samples passed the Kolmogorov-Smirnov test for normality. One-way and two-way ANOVA comparisons were corrected for multiple comparisons using Tukey’s or Šidák’s multiple comparisons tests, as indicated in figure legends. For comparison of two non-normally distributed distributions, Wilcoxon matched-pairs signed-rank tests, permutation tests, or Mann-Whitney tests were used. When one-sample t-tests were performed to determine mean deviations from a specific value, p-values were adjusted using the false discovery rate method of the Benjamini-Hochberg process. For comparisons of discrete variables, we used chi-square (when more than two outcomes are possible), Fisher’s exact test (when only two outcomes are compared), or Cochran’s Q-test (for paired comparisons), followed by the false-discovery rate method of the Benjamini-Hochberg process. All p-values < 0.05 were indicated on the figure when space was allowed.

---

**Extended Data Fig. 1.**
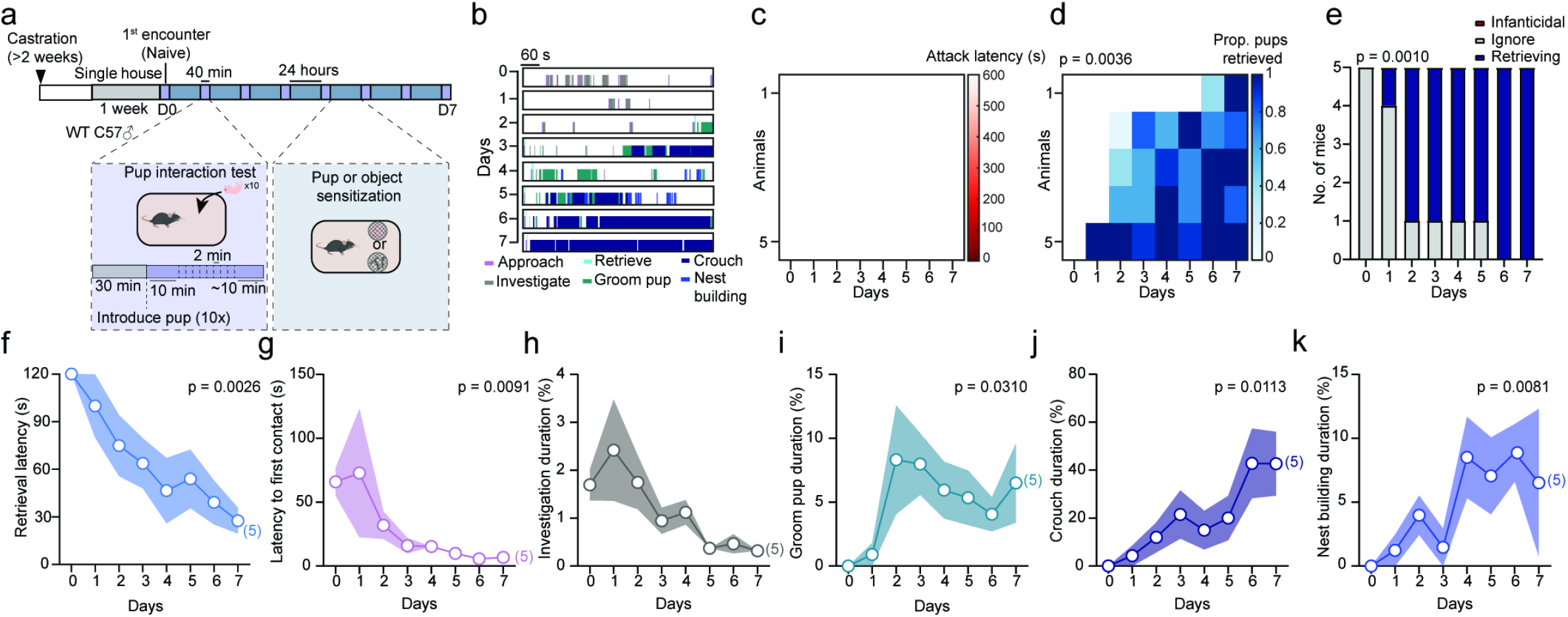
Pup sensitization promotes paternal behavior independent of sex hormones. **(a)** Pup sensitization paradigm. **(b)** Representative raster plots showing pup-directed behaviors in the pup interaction test. Raster plots represent the 10 minutes after introducing the last pup during the pup interaction test. **(c, d)** The latency to attack **(c)** and proportion of pups retrieved **(d)** during the pup interaction test across 7 days of pup sensitization. Day 0: before pup sensitization, days 1-7: during pup sensitization. If no attack occurred, the latency was scored as 600 s. **(e)** The number of mice that were infanticidal, non-infanticidal and non-parental (ignore), or retrieving during the pup interaction test across 7 days of pup sensitization. **(f-k)** The latency to retrieve pups **(f)**, latency to contact pups **(g)**, pup investigation duration **(h)**, grooming pup duration **(i)**, crouching over pups duration **(j)**, and nest building duration **(k)** during the pup interaction test across 7 days of pup sensitization. Day 0: before pup sensitization, days 1-7: during pup sensitization. If no retrieval occurred, the latency was scored as 120 s. Mouse cartoons in this and all other figures are adopted from scidraw.io. Lines and shaded error bars represent mean ± SEM. **(c, d, f, g, i-k)** Friedman test followed by Dunn’s multiple comparisons test (post-hoc test results not shown). **(e)** Cochran’s Q test followed by Benjamini Hochberg false-discovery rate correction. **(h)** One-way ANOVA with repeated measures followed by Tukey’s multiple comparisons test. Lines and shaded error bars represent mean ± SEM. All p < 0.05 are indicated.

**Extended Data Fig. 2.**
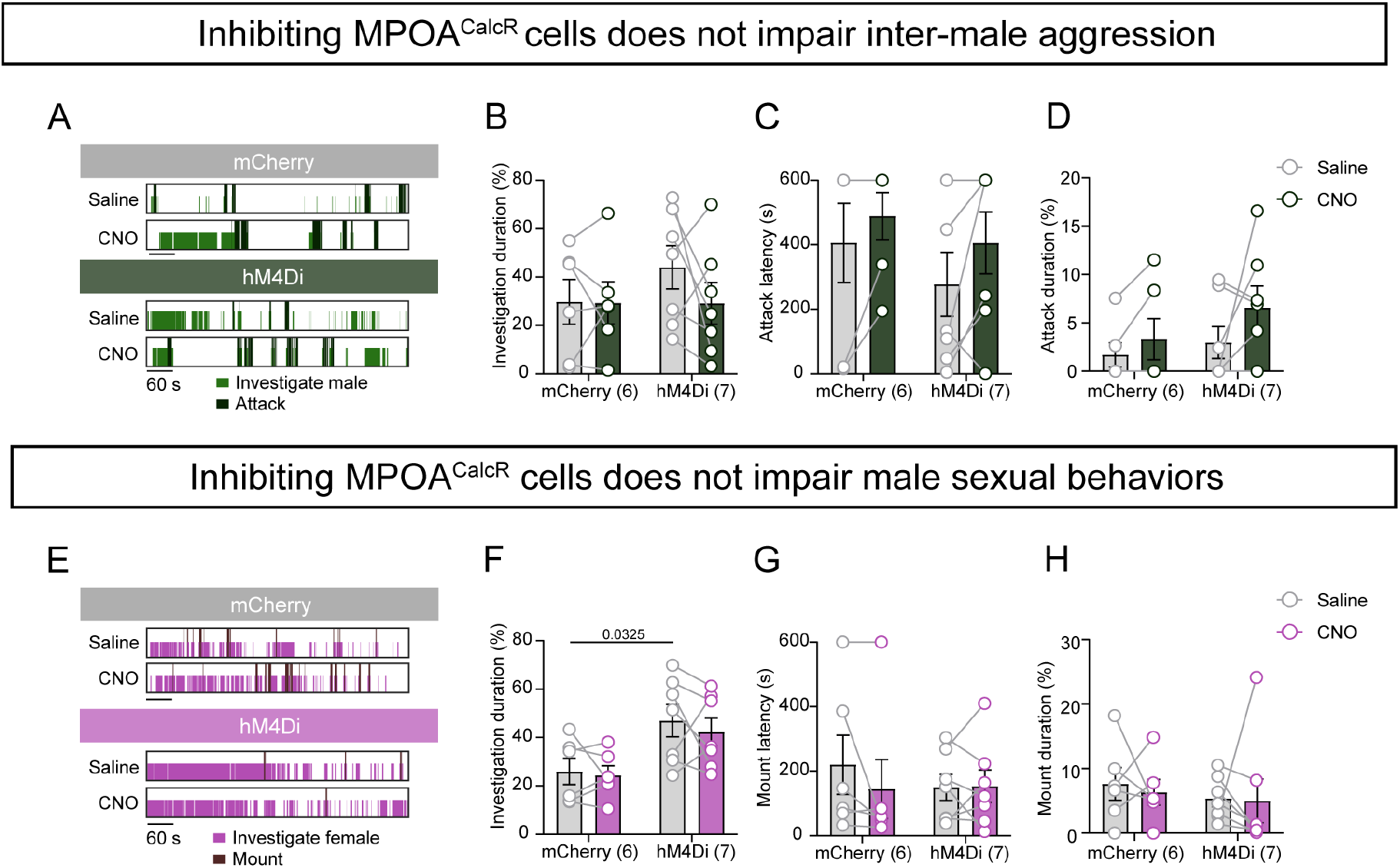
Chemogenetic inhibition of MPOA^CalcR^ cells does not affect behaviors toward adult males or females. **(a)** Representative raster plots showing male-directed behaviors following saline or CNO injection in an example mCherry and hM4Di animal. **(b-d)** Male investigation duration **(b)**, latency to attack male **(c)**, and attack male duration **(d)** in mCherry and hM4Di mice after saline or CNO injection. **(e)** Representative raster plots showing female-directed behaviors following saline or CNO injection in an example mCherry and hM4Di animal. **(f-h)** Female investigation duration **(f)**, latency to mount female **(g)**, and mount female duration **(h)** in mCherry and hM4Di mice after saline or CNO injection. Numbers in parentheses indicate the subject animal number. Circle-line pairs represent data of individual animals. **(b-d, f-h)** Two-way ANOVA with repeated measures followed by Šidák’s multiple comparisons test. Bars and error bars represent mean ± SEM. All p < 0.05 are indicated.

**Extended Data Fig. 3.**
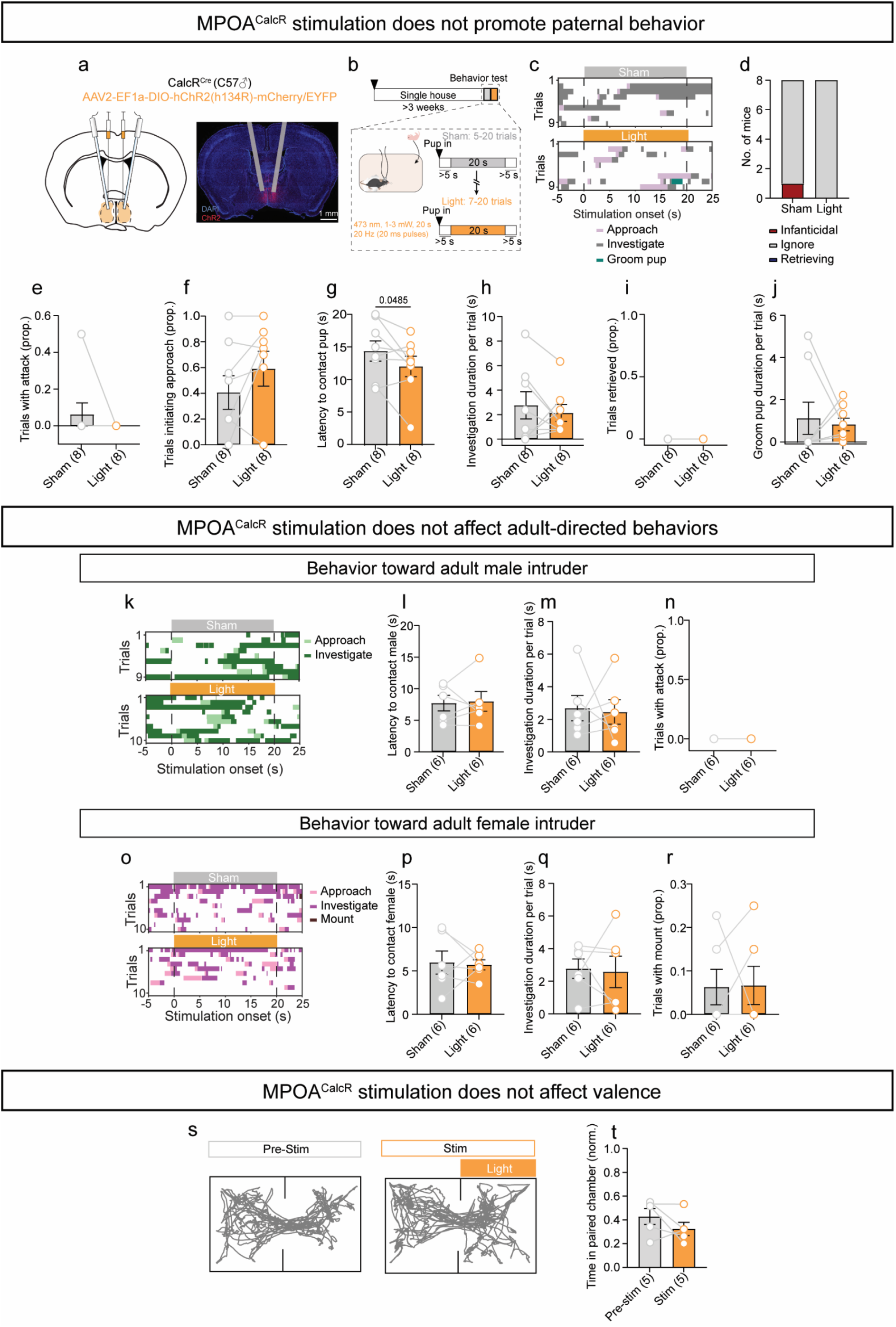
Behaviors associated with optogenetic stimulation of MPOA^CalcR^ cells. **(a)** Virus injection and optical fiber implantation schematic and representative histology image. **(b)** Timeline to optogenetically activate MPOA^CalcR^ cells. **(c)** Representative raster plots showing pup-directed behaviors aligned to the onset of sham or light stimulation in a ChR2 animal. **(d)** The number of mice that were infanticidal, non-infanticidal and non-retrieving (ignore), and retrieving following sham or light stimulation. **(e)** Proportion of sham or light trials with pup-directed attack. **(f)** Proportion of sham or light trials with pup approach. **(g)** The latency to contact pups following light or sham onset. **(h)** The time spent investigating pups during sham or light stimulation. **(i)** Proportion of sham or light trials with pup retrieval. **(j)** The time spent grooming pups during sham or light stimulation. **(k)** Representative raster plots showing male-directed behaviors aligned to the onset of sham or light stimulation in a ChR2-injected animal. **(l)** The latency to contact the male intruder following light or sham onset. **(m)** The time spent investigating the male during sham or light stimulation. **(n)** Proportion of sham or light trials with attack toward the male intruder. **(o)** Representative raster plots showing female-directed behaviors aligned to the onset of sham or light stimulation in a ChR2 animal. **(p)** The latency to contact the female intruder following light or sham onset. **(q)** The time spent investigating the female during sham or light stimulation. **(r)** Proportion of sham or light trials with mounting toward the female intruder **(s)** Representative tracking results from the real-time place preference assay (10 minutes pre-stimulation and 10 minutes during stimulation). **(t)** Proportion of time spent in the light-paired chamber before and during the light-pairing period (2mW, 20 ms, 20Hz). The brain illustration was produced based on a reference atlas from https://atlas.brain-map.org/. Numbers in parentheses indicate the subject animal numbers. Circle-line pairs represent data of individual animals. **(d)** McNemar’s test. **(e, j, n, q, r)** Wilcoxon test. **(f-h, l, m, p, t)** Paired t-test. Bars and error bars represent mean ± SEM. All p < 0.05 are indicated.

**Extended Data Fig. 4.**
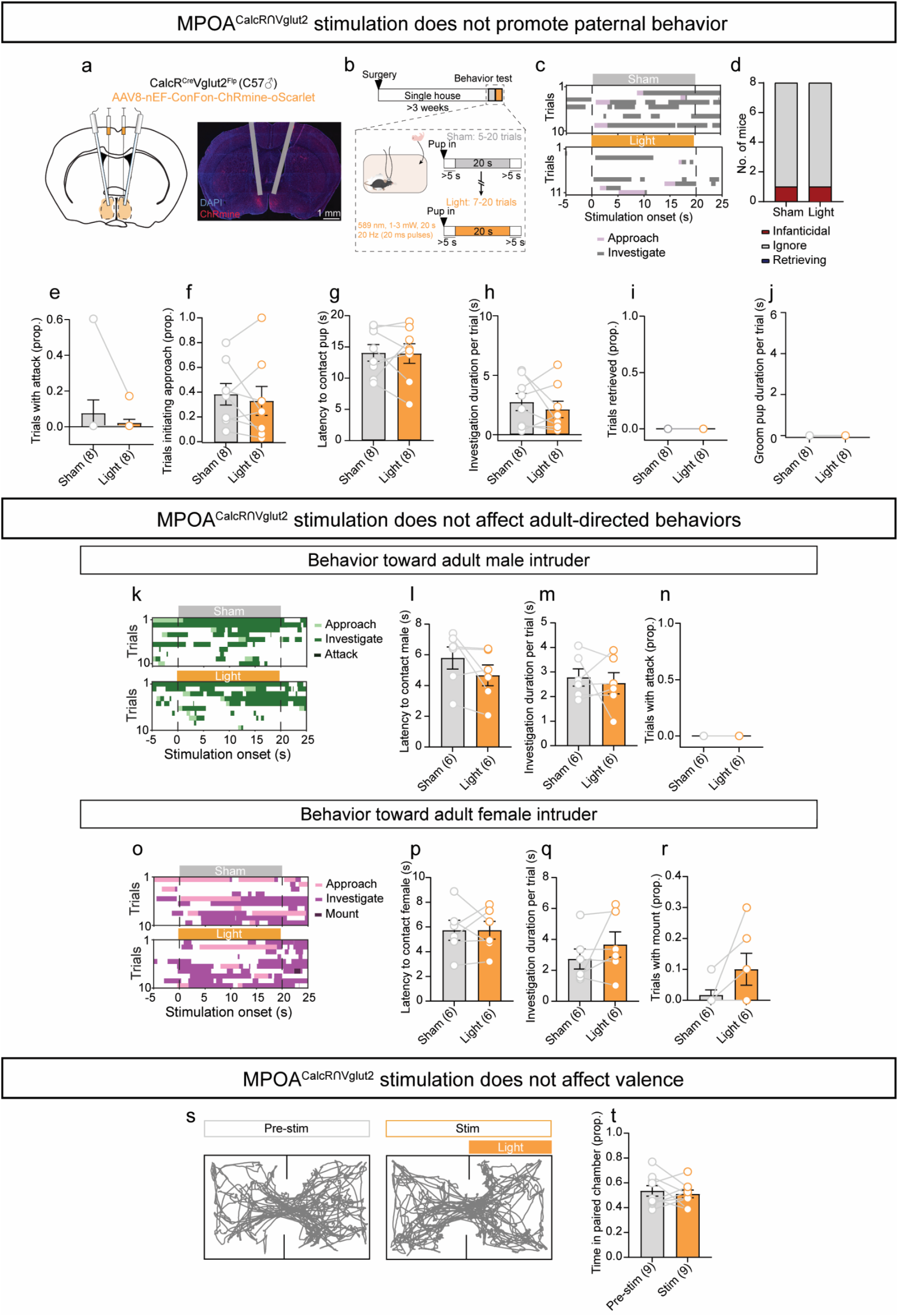
Behaviors associated with optogenetic stimulation of MPOA^CalcR∩Vglut2^ cells. **(a)** Virus injection and optical fiber implantation schematic and representative histology image. **(b)** Timeline to optogenetically activate MPOA^CalcR∩Vglut2^ cells. Mouse cartoons in this and all other figures are adopted from scidraw.io. **(c)** Representative raster plots showing pup-directed behaviors aligned to the onset of sham or light stimulation in a ChRmine-injected animal. **(d)** The number of mice that were infanticidal, non-infanticidal and non-retrieving (ignore), and retrieving following sham or light stimulation. **(e)** Proportion of sham or light trials with pup-directed attack **(f)** Proportion of sham or light trials with with pup approach. **(g)** The latency to contact pups following light or sham onset. **(h)** The time spent investigating pups during sham or light stimulation. **(i)** Proportion of sham or light trials with pup retrieval. **(j)** The time spent grooming pups during sham or light stimulation. **(k)** Representative raster plots showing male-directed behaviors aligned to the onset of sham or light stimulation in a ChRmine-injected animal. **(l)** The latency to contact the male intruder following light or sham onset. **(m)** The time spent investigating the male during sham or light stimulation. **(n)** Proportion of sham or light trials with attack toward the male intruder. **(o)** Representative raster plots showing female-directed behaviors aligned to the onset of sham or light stimulation in a ChRmine-injected animal. **(p)** The latency to contact the female intruder following light or sham onset. **(q)** The time spent investigating the female during sham or light stimulation. **(r)** Proportion of sham or light trials with mounting toward the female intruder **(s)** Representative tracking results from the real-time place preference assay (10 minutes pre-stimulation and 10 minutes during stimulation). **(t)** Proportion of time spent in the light-paired chamber before and during the light-paring period (2mW, 20 ms, 20Hz). Numbers in parentheses indicate the subject animal number. Circle-line pairs represent data of individual animals. **(d)** McNemar’s test. **(e, r)** Wilcoxon test. **(f-h, l, m, p, q, t)** Paired t-test. Bars and error bars represent mean ± SEM. All p < 0.05 are indicated.

**Extended Data Fig. 5.**
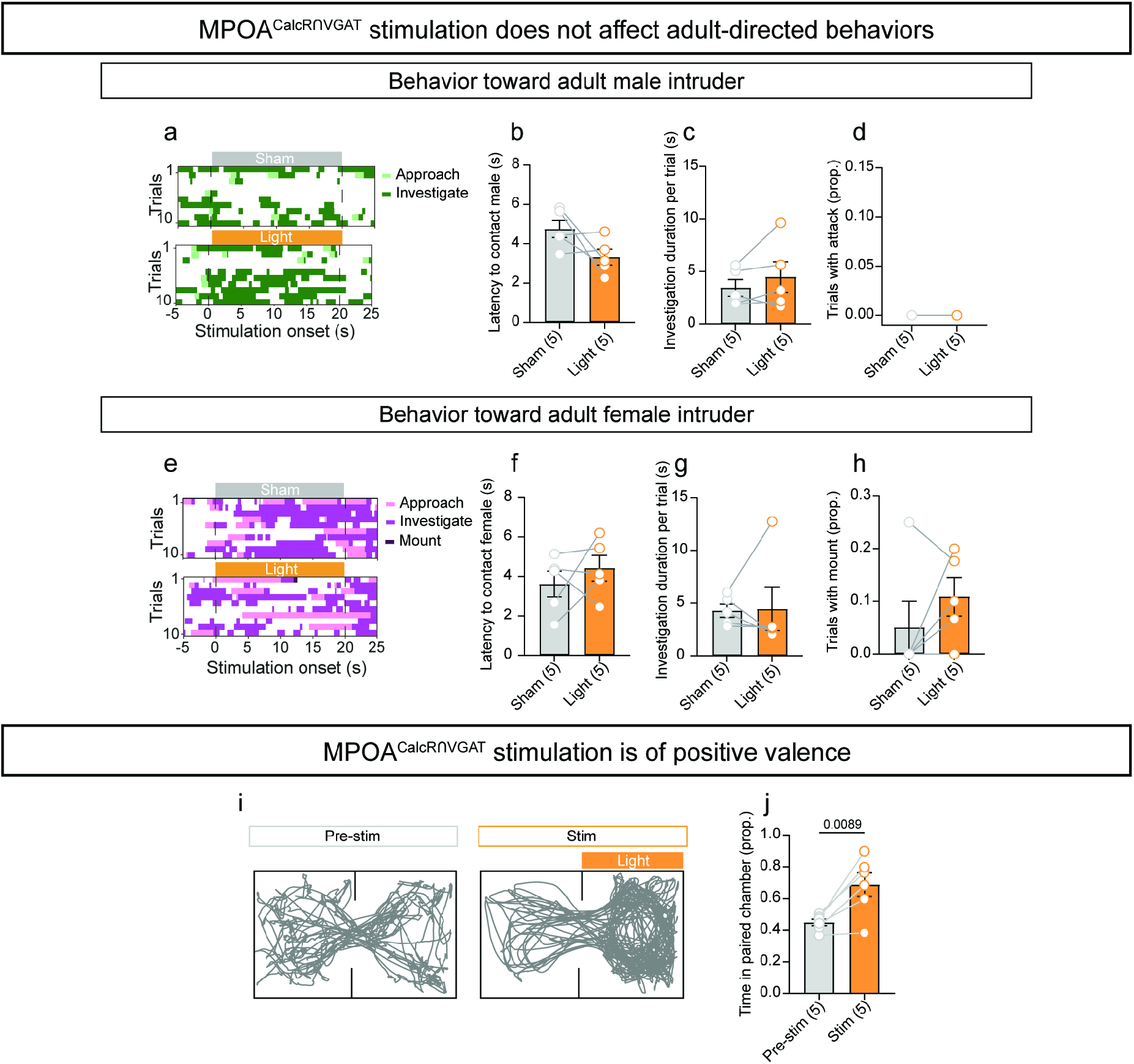
Additional behaviors associated with optogenetic stimulation of MPOA^CalcR∩VGAT^ cells. **(a)** Representative raster plots showing male-directed behaviors aligned to the onset of sham or light stimulation in a ChRmine-injected animal. **(b)** The latency to contact the male intruder following light or sham onset. **(c)** The time spent investigating the male during sham or light stimulation. **(d)** Proportion of sham or light trials with attack toward the male intruder. **(e)** Representative raster plots showing female-directed behaviors aligned to sham or light stimulation onset in a ChRmine-injected animal. **(f)** The latency to contact the female intruder following light or sham onset. **(g)** The time spent investigating the female during sham or light stimulation. **(h)** Proportion of sham or light trials with mounting toward the female intruder. **(i)** Representative tracking results from the real-time place preference assay (10 minutes pre-stimulation and 10 minutes during stimulation). **(j)** Proportion of time spent in the light-paired chamber before and during the light-paring period (2mW, 20 ms, 20Hz). Numbers in parentheses indicate the subject animal number. Circle-line pairs represent data of individual animals. **(b, c, f-h, j)** Paired t-test. **(d)** Wilcoxon test. Bars and error bars represent mean ± SEM. All p < 0.05 are indicated.

**Extended Data Fig. 6.**
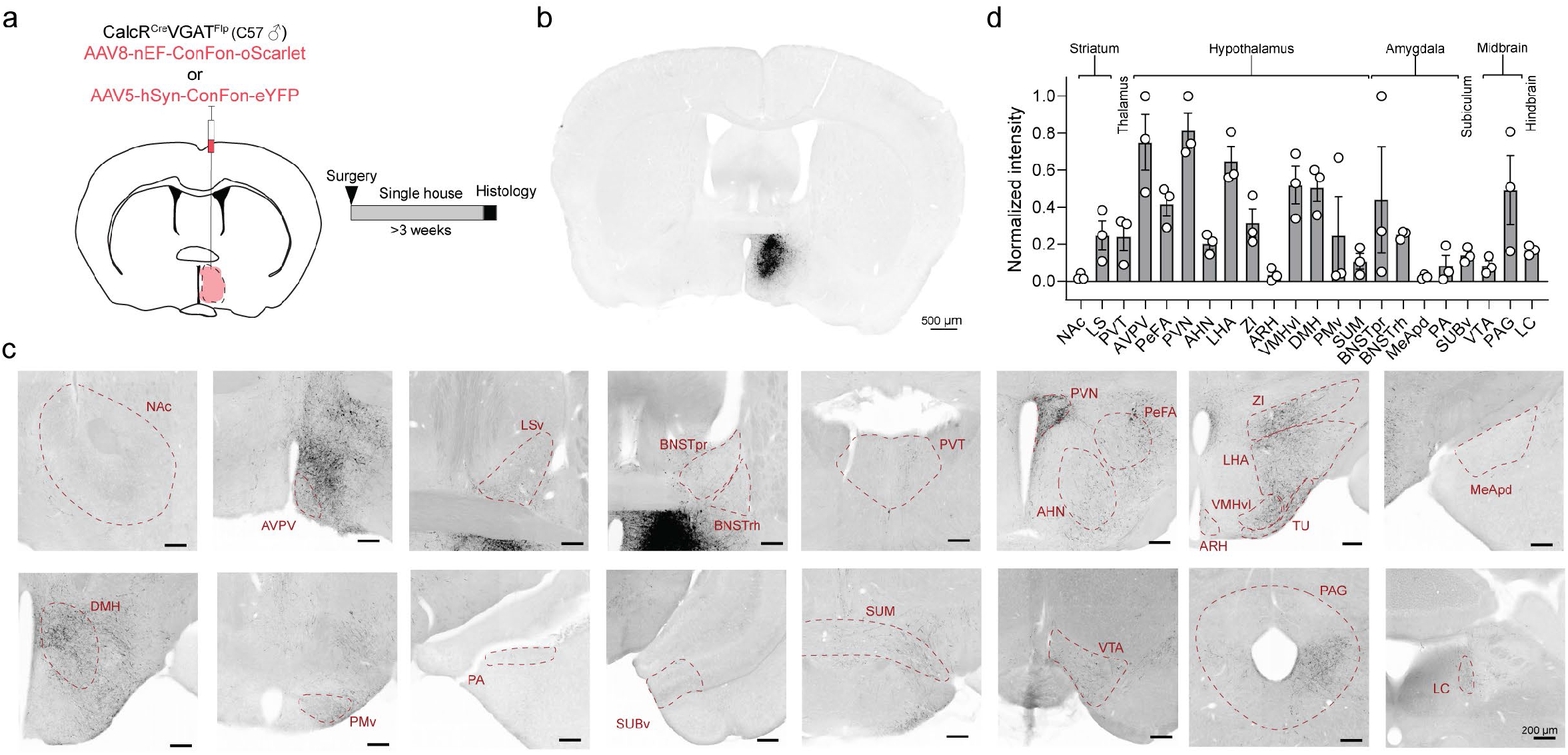
Anterograde tracing of MPOA^CalcR∩VGAT^ cells. **(a)** Virus injection schematic and timeline to trace the outputs of MPOA^CalcR∩VGAT^ cells. **(b)** Representative histology image showing the infected cells in a CalcR^Cre^VGAT^Flp^ mouse. Black: EYFP expression. **(c)** Representative histology images showing MPOA^CalcR∩VGAT^ projections in various downstream regions. **(d)** The normalized intensity of MPOA^CalcR∩VGAT^ projections in downstream regions. Bars and error bars represent mean ± SEM. Abbreviations: NAc: nucleus accumbens; AVPV: anteroventral periventricular nucleus; LSv: ventral lateral septum; BNSTpr: principal nucleus of the bed nucleus of the stria terminalis; BNSTrh: rhomboid nucleus of the bed nucleus of the stria terminalis; PVT: paraventricular thalamus; PVN: paraventricular nucleus of the hypothalamus; PeFA: perifornical area; AHN: anterior hypothalamic nucleus; ZI: zona incerta; LHA: lateral hypothalamus; VMHvl: ventrolateral part of the ventromedial hypothalamus; TU: tuberal nucleus; ARH: arcuate nucleus of the hypothalamus; MeApd: posterodorsal part of the medial amygdala; DMH: dorsomedial hypothalamus; PMv: ventral premammillary nucleus; PA: posterior amygdala; SUBv: ventral subiculum; SUM: supramammillary nucleus; VTA: ventral tegmental area; PAG: periaqueductal gray; LC: locus coeruleus.

**Extended Data Fig. 7.**
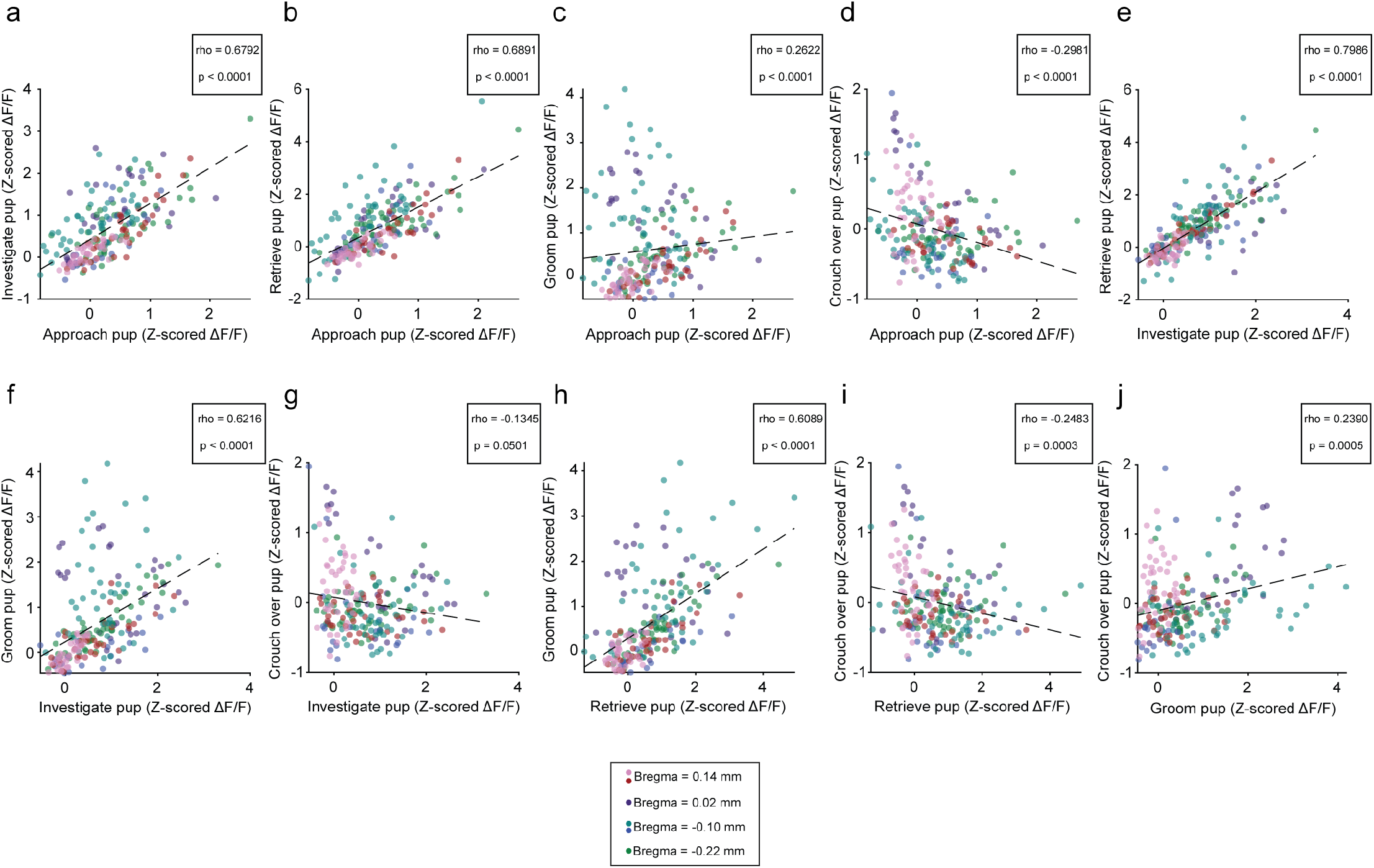
Correlations of MPOA^CalcR^ activity between pairs of parental behaviors. Scatter plots showing the the average Z-scored ΔF/F during **(a)** approach pup vs. investigate pup, **(b)** approach pup vs. retrieve pup, **(c)** approach pup vs. groom pup, **(d)** approach pup vs. crouch over pup, **(e)** investigate pup vs. retrieve pup, **(f)** investigate pup vs. groom pup, **(g)** investigate pup vs. crouch over pup, **(h)** retrieve pup vs. groom pup, **(i)** retrieve pup vs. crouch over pup, and **(j)** groom pup vs. crouch over pup. Circles represent individual cells. Circle colors correspond to individual animals. Dashed line indicates the regression line. Spearman correlation rho and p-values indicated on the figure.

**Extended Data Fig. 8.**
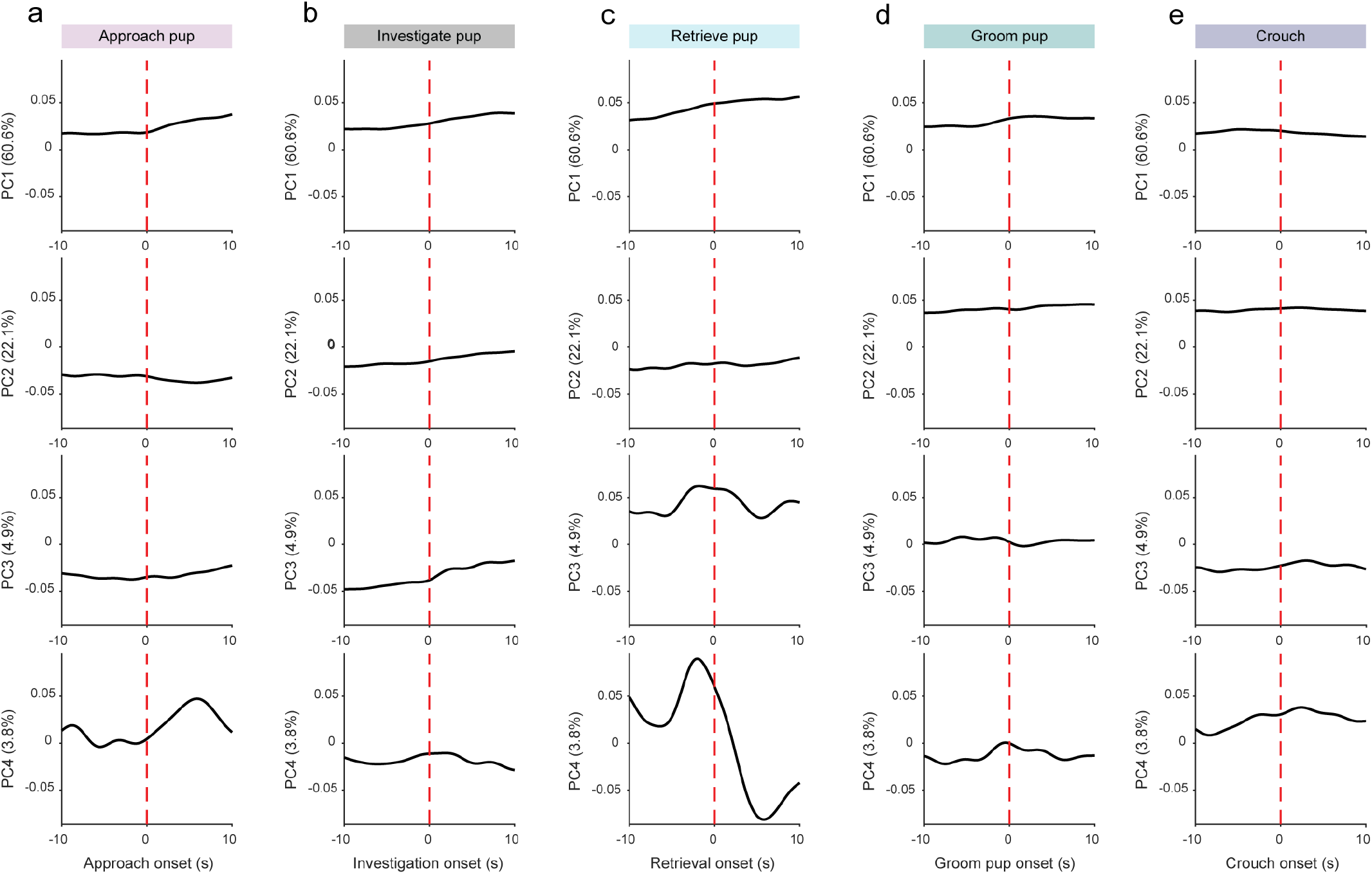
Loading vectors of principal components from dimensionality reduction of 2-photon imaging of MPOA^CalcR^ cells during various parental behaviors. **(a-e)** Loading scores of principal components (PCs) aligned to behavior onset during pup-directed approach **(a)**, investigation **(b)**, retrieval **(c)**, pup grooming **(d)**, and crouching **(e)**. Top: PC1, Second row: PC2, Third row: PC3, Bottom: PC4. Percentages in parentheses denote the percentage of variance explained by each principal component. Dashed lines indicate behavior onset.

**Extended Data Fig. 9.**
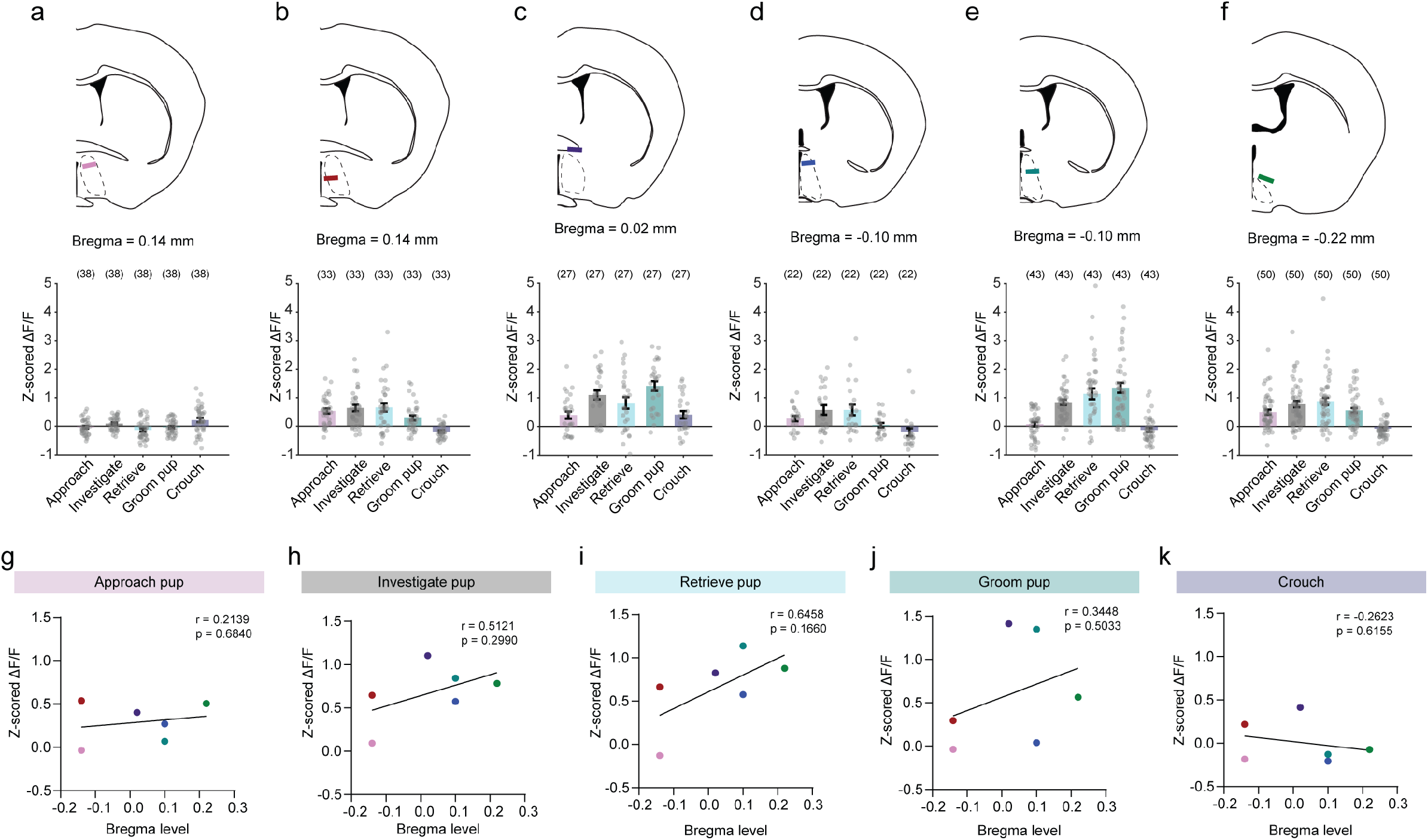
GRIN lens locations and parental behavior-evoked activity of MPOA^CalcR^ cells. **(a-f)** Top: locations of GRIN lenses (colored lines) for each mouse showing robust parental behaviors, ordered by anterior-posterior lens position. **a** and **b**: Bregma level 0.14 mm; **c**: Bregma level 0.02 mm; **d** and **e**: Bregma level - 0.10 mm; **f**: Bregma level -0.22 mm. Bottom: the average Z-scored ΔF/F of MPOA^CalcR^ cells during various parental behaviors of each animal. **(g-k)** The correlation between MPOA^CalcR^ activity and GRIN lens location during pup-directed approach **(f)**, investigation **(h)**, retrieval **(i)**, pup grooming **(j)**, and crouching **(k)**. Each circle represents the average Z-scored ΔF/F from all recorded cells in one mouse. Colors matched GRIN lens line color in **a-f**. Numbers in parentheses indicate the cell number. Circles represent data of individual cells **(a-f)** or individual animals **(g-k)**. **(g-k)** Simple linear correlation. Bars and error bars represent mean ± SEM. All p < 0.05 are indicated.

**Extended Data Fig. 10.**
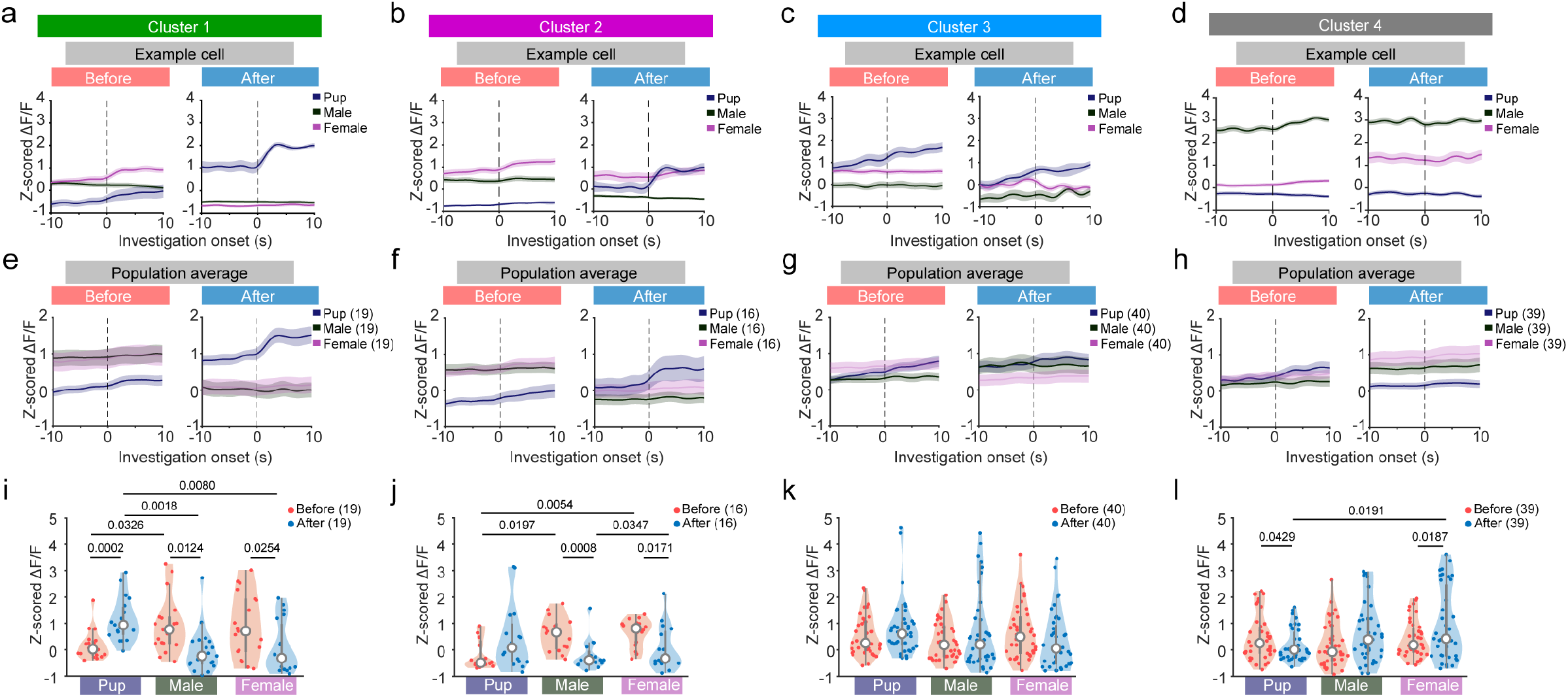
MPOA^CalcR^ cell responses to social stimuli before and after pup sensitization. **(a-d)** PETHs of Z-scored ΔF/F aligned to pup investigation, male investigation, and female investigation from example l ngitudinally tracked MPOA^CalcR^ cells in 1^st^ **(a)**, 2^nd^ **(b)**, 3^rd^ **(c)** and 4^th^ **(d)** clusters. Cluster membership is based on Fig. 3e, g, h. The cells were tracked from before (left) to after (right) pup sensitization. **(e-h)** Averaged PETHs of Z-scored ΔF/F of all cells in each cluster before and after pup sensitization. **(e)** 1^st^ Cluster: n = 19 cells. **(f)** 2^nd^ Cluster: n = 16 cells. **(g)** 3^rd^ Cluster: n = 40 cells. **(h)** 4^th^ Cluster: n = 39 cells. **(i-l)** Average Z-scored ΔF/F during pup, male, and female investigation before and after pup sensitization of cells in 1^st^ **(i)**, 2^nd^ **(j)**, 3^rd^ **(k)** and 4^th^ **(l)** clusters. Red: before pup sensitization; Blue: after pup sensitization. Numbers in parentheses or axis titles indicate the cell number. Circles represent data of individual cells. **(i-l)** Two-way repeated measures ANOVA followed by Tukey’s multiple comparisons test. Lines and shaded error bars represent mean ± SEM. Violin plots: white circles represent the median and interquartile and 1.5x interquartile ranges are represented by thick and thin gray bars, respectively; shaded areas represent the range of all values. All p < 0.05 are indicated.

**Extended Data Fig. 11.**
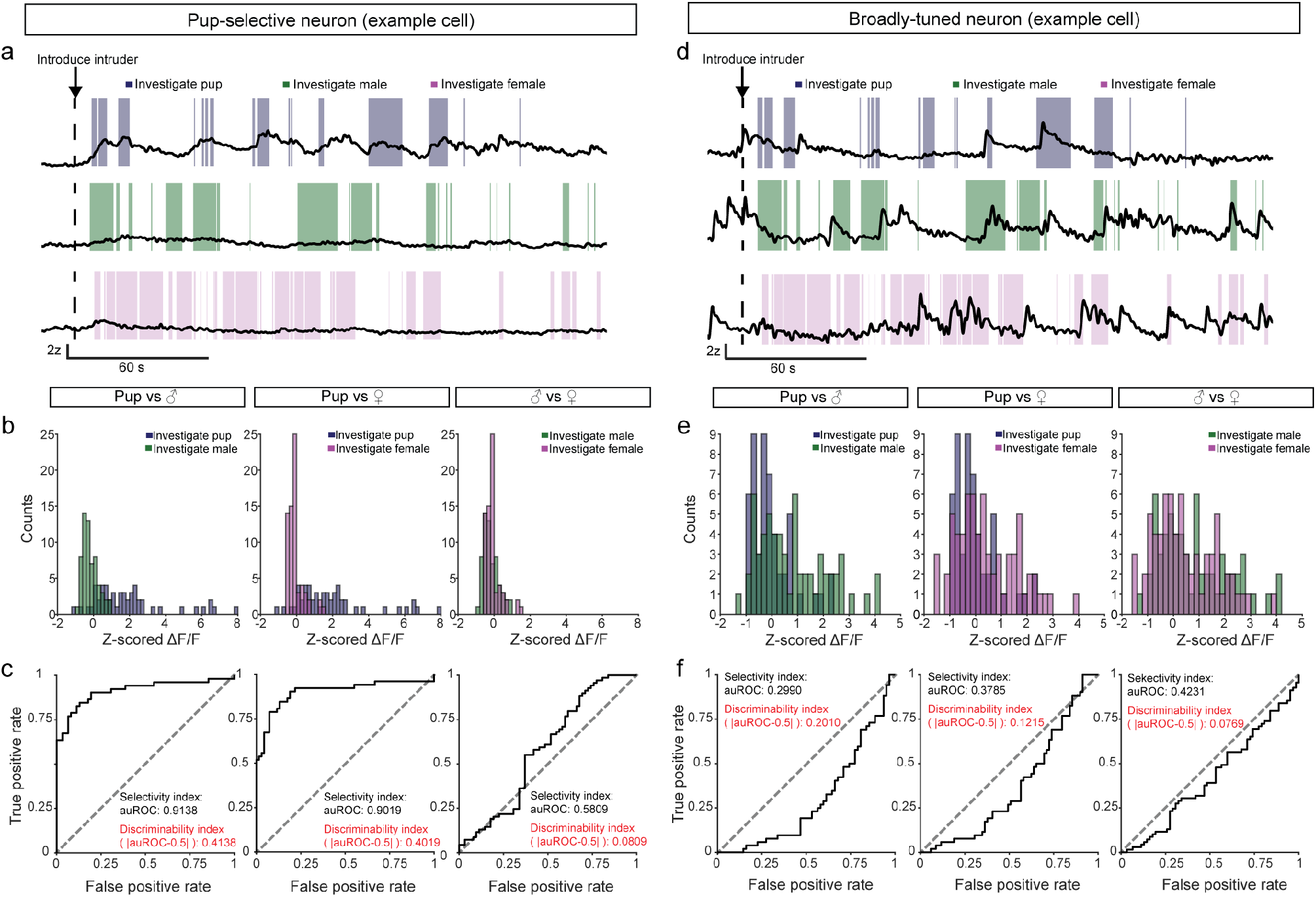
Calculation of discriminability index of MPOA^CalcR^ responses. **(a, d)** Traces of one representative pup-selective neuron **(a)** and one broadly-tuned neuron **(d)** during pup (top), male (middle), and female (bottom) interactions. **(b, e)** The distributions of z-scored ΔF/F activity during various behavior bouts of the pup-selective neuron represented in **a (b)** and broadly-tuned neuron represented in **d (e)**. Left: Distributions of pup and male investigation-evoked activity. Middle: Distributions of pup and female investigation-evoked activity. Right: Distributions of male and female investigation-evoked activity. **(c, f)** The receiving operating characteristic (ROC) curve, area under the ROC curve (auROC), and discriminability index between pairs of stimuli for the pup-selective cell represented in **a (c)** and broadly-tuned neuron represented in **d (e)**.

**Extended Data Fig. 12.**
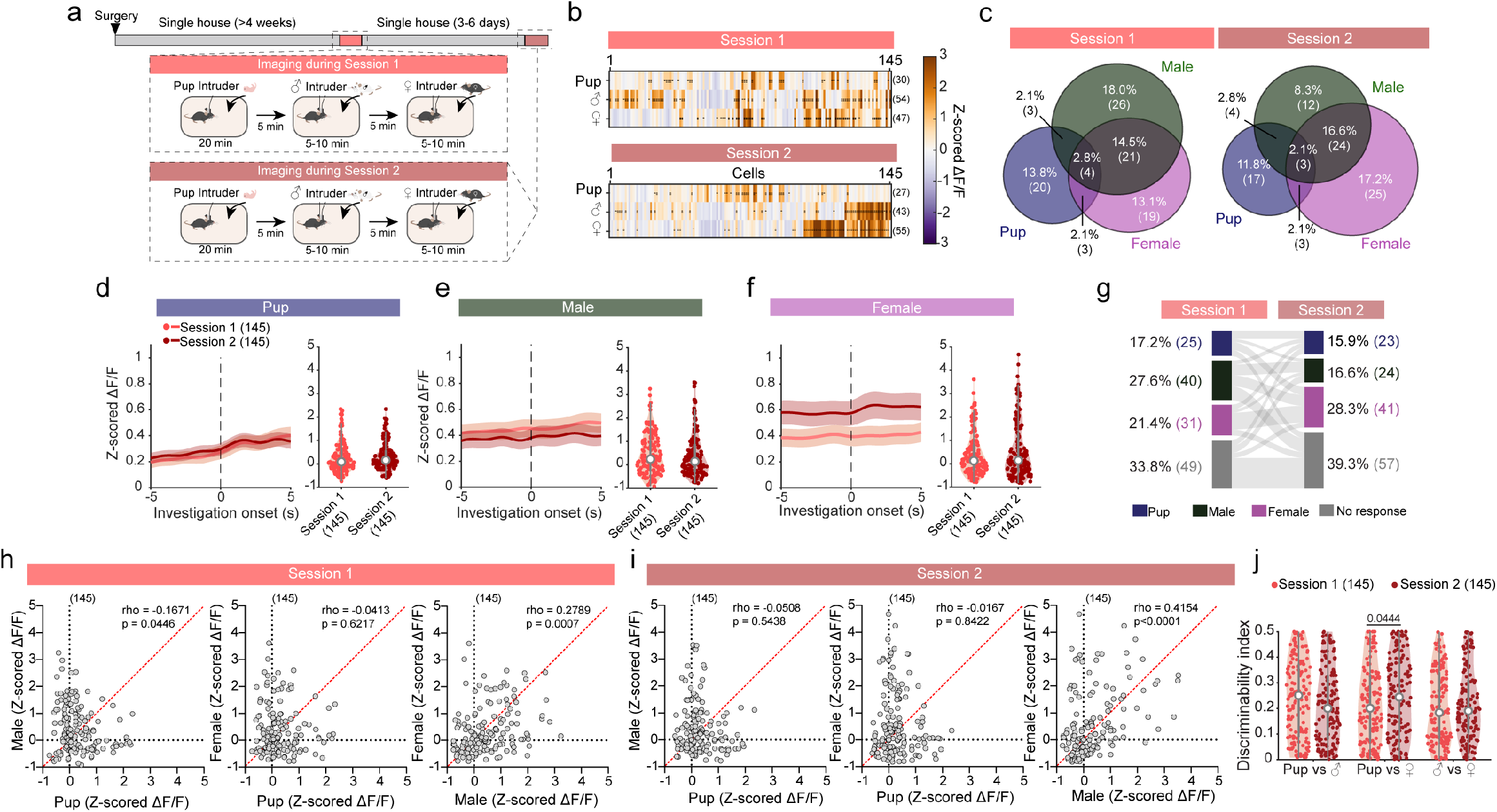
Comparing responses of MPOA^CalcR^ cells to social stimuli across days without pup sensitization. **(a)** Timeline to image the activity of the same set of MPOA^CalcR^ cells during pup, male, and female interactions across two sessions without pup sensitization. **(b)** Heatmaps showing averaged Z-scored ΔF/F of MPOA^CalcR^ cells during pup, male, and female investigation. Cells are arranged in the same order before (top) and after pup sensitization (bottom) based on their membership in k-means clustering analyses (n = 145 longitudinally-imaged cells from 7 mice). Numbers on the right indicate the number of significantly responsive cells. Black circles indicate q values. ●q<0.01; ●●q<0.001. **(c)** Venn diagrams showing cells significantly responsive to pup, male, and female investigation during the first and second session. **(d-f)** Left: population-averaged PETHs during pup **(d)**, male **(e)**, and female **(f)** investigation. Dashed lines indicate the onset of behavior. Light red: first session, Dark red: second session. Right: Average Z-scored ΔF/F of each cell during pup **(d)**, male **(e)**, and female **(f)** investigation. Light red: first session, Dark red: second session. **(g)** Alluvial diagram showing pup-, male-, and female-preferred and non-responsive cells. (left: first session, right: second session; n = 145 cells from 7 mice). **(h, i)** Correlations of cell responses (Z-scored ΔF/F) during pup and male investigation (left), pup and female investigation (middle), and male and female investigation (right) during the first **(h)** and second **(i)** session. **(j)** Discriminability index of cell responses (Z-scored ΔF/F) across stimulus pairs. Numbers in parentheses or axis titles indicate the cell number. **(b)** One-sample t-test followed by Benjamini Hochberg false-discovery rate correction. **(c, g)** Chi-square test. **(d-f)** Permutation test. **(h, i)** Spearman correlation. **(g)** Two-way ANOVA with repeated measures followed by Tukey’s multiple comparisons test. Lines and shaded error bars represent mean ± SEM. Violin plots: each circle (light red and dark red) represents one cell; white circles represent the median and interquartile, and 1.5x interquartile ranges are represented by thick and thin gray bars, respectively; shaded areas represent the range of all values. All p < 0.05 are indicated.

**Extended Data Fig. 13.**
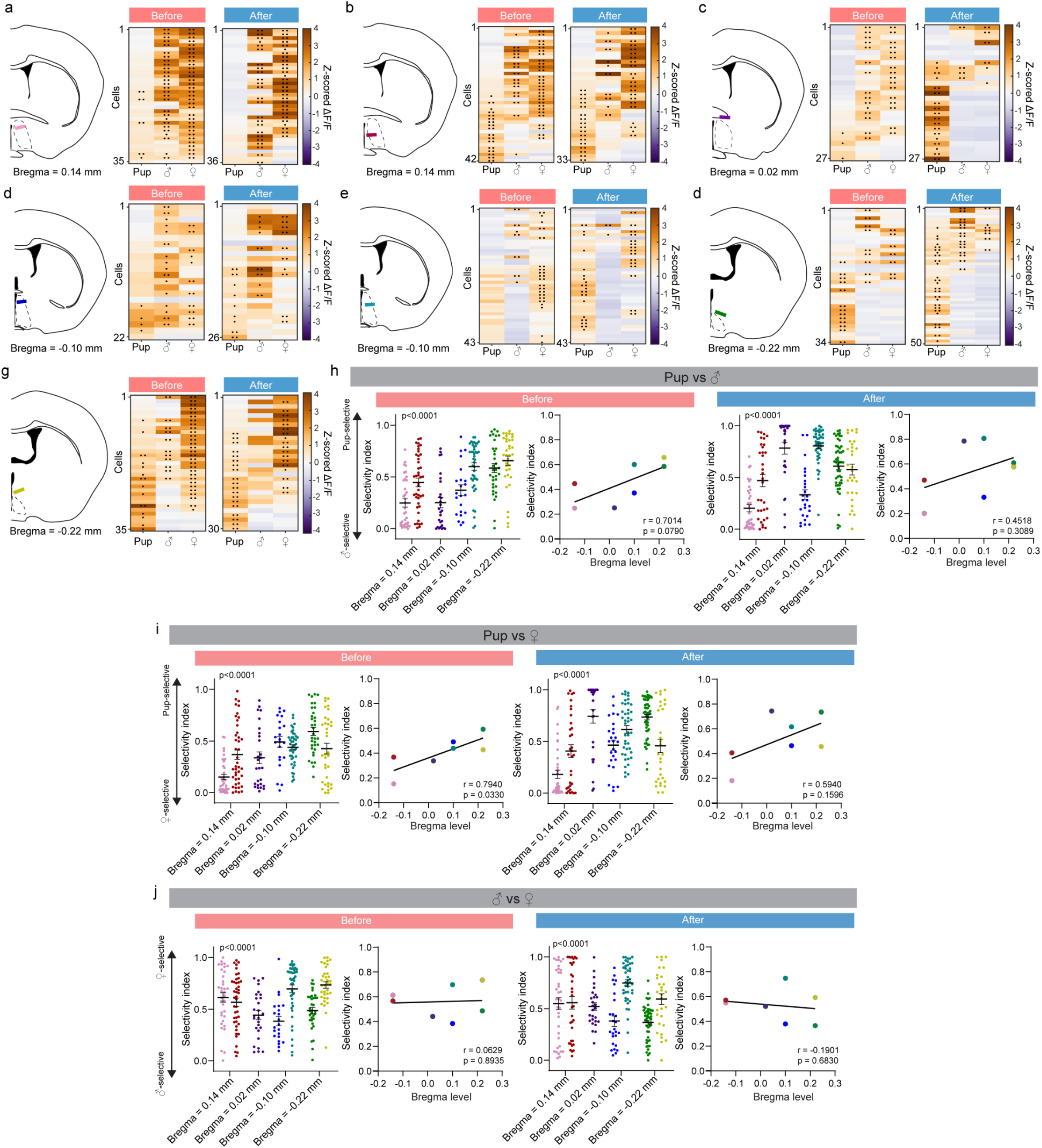
GRIN lens locations and relationship to social stimulus tuning. **(a-g)** Left: Locations of GRIN lenses (colored lines) for each mouse, ordered by Bregma level. Middle: MPOA^CalcR^ cell averaged Z-scored ΔF/F before pup sensitization during pup, male, and female investigation (n = 35 cells in **a**, n = 42 cells in **b**, n = 27 cells in **c**, n = 22 cells in **d**, n = 43 cells in **e**, n = 34 cells in **f**, n = 35 cells in **g**). Right: MPOA^CalcR^ cell averaged Z-scored ΔF/F after pup sensitization during pup, male, and female investigation (n = 36 cells in **a**, n = 33 cells in **b**, n = 27 cells in **c**, n = 26 cells in **d**, n = 43 cells in **e**, n = 50 cells in **f**, n = 30 cells in **g**). Black circles indicate q values. ●q<0.01; ●●q<0.001. The brain illustrations were produced based on a reference atlas from https://atlas.brain-map.org/ **(h)** Left: Selectivity index between pup and male for each cell in every mouse before (left) and after (right) pup sensitization, ordered by Bregma level, and correlation between Bregma level and selectivity index. Selectivity index is calculated as: auROC between the distributions of male- and pup-evoked z-scored ΔF/F. Before (left) and after (left): each circle corresponds to one cell; each color corresponds to one mouse (colors matched GRIN lens line color in **a-g**). Before (right) and after (right): Correlation showing the relationship between Bregma level and pup selectivity. Each circle corresponds to the average selectivity from all cells in one mouse. **(i)** Selectivity index between pup and female for all cells in every mouse (before, left and after, left) and the average selectivity index of all cells in each mouse (before, right and after, right) before and after pup sensitization. Follows the color convention in **h**. **(j)** Selectivity index between male and female for all cells in every mouse (before, left and after, left) and the average selectivity index of all cells in each mouse (before, right and after, right) before and pup sensitization. Follows the color convention in **h**. **(a-g)** One-sample t-test followed by Benjamini Hochberg false-discovery rate correction. **(h-j before and after, left)** Kruskal-Wallis test followed by Dunn’s multiple comparisons test (post-hoc test results not shown). **(h-j before and after, right)** Simple linear correlation. Lines and error bars represent mean ± SEM.

**Extended Data Fig. 14.**
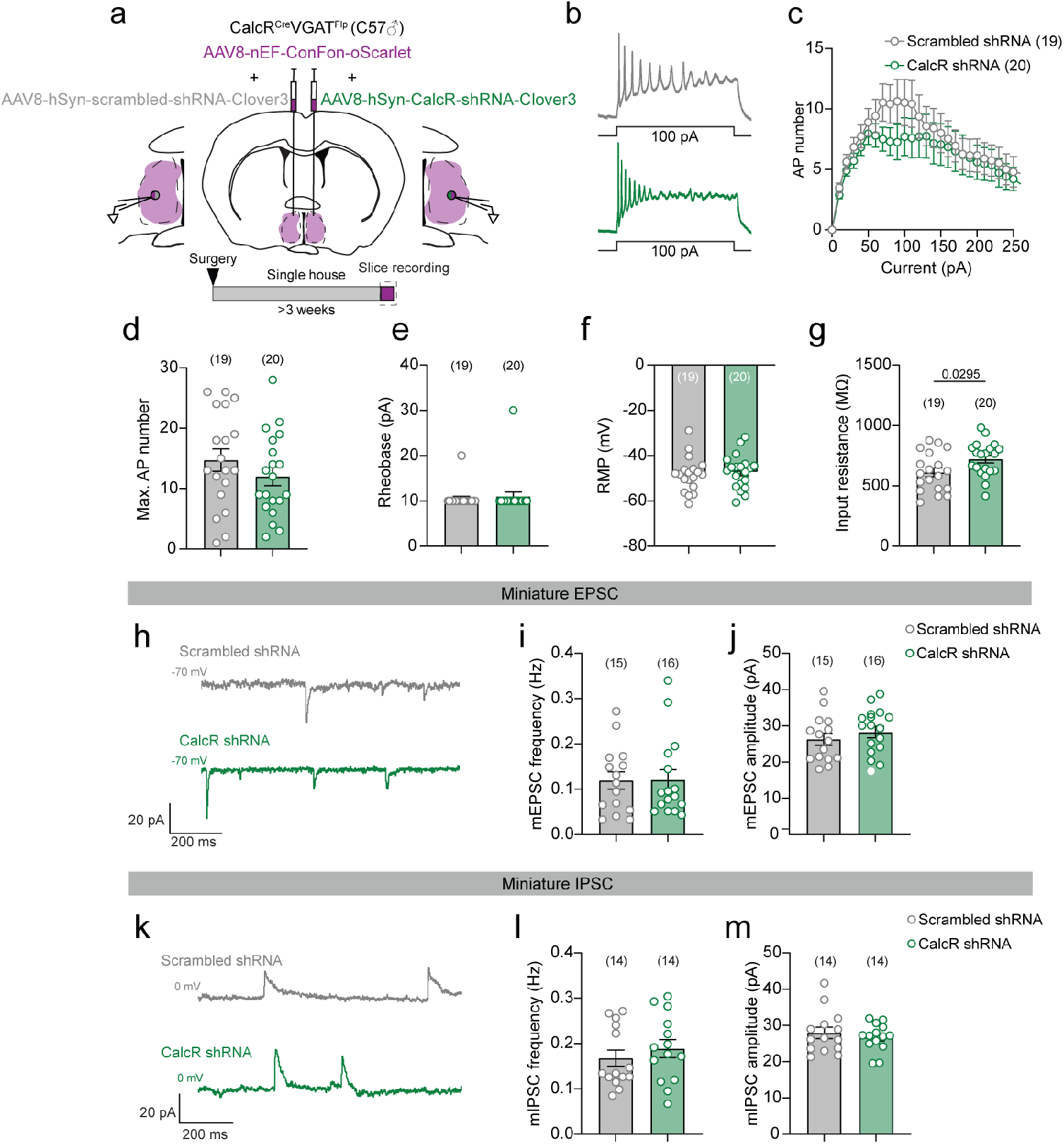
Physiological characterization of MPOA^CalcR∩VGAT^ cells after *Calcr* knockdown. **(a)** Virus injection, recording schematic, and experimental timeline for in vitro whole-cell recordings of _MPOA_^CalcR∩VGAT^ _cells._ **(b)** Left: Representative current-clamp recording traces of MPOA^CalcR∩VGAT^ cells after 100pA current injection. Right: F-I curve of MPOA^CalcR∩VGAT^ cells expressing scrambled shRNA or CalcR shRNA. **(d-e)** Maximum action potential number across current steps **(d)**, rheobase **(e)**, resting membrane potential **(f)**, and input resistance **(g)** of MPOA^CalcR∩VGAT^ cells expressing scrambled shRNA or CalcR shRNA (scrambled shRNA: n = 19 cells from 4 mice; CalcR shRNA: n = 20 cells from 4 mice). **(h, k)** Representative voltage-clamp recording traces of MPOA^CalcR∩VGAT^ cells in 1 μM TTX. Numbers indicate the holding potential. **(i, j, l, m)** The frequency **(i, l)** and amplitude **(j, m)** of mEPSCs **(i, j)** and mIPSCs **(l, m)** of MPOA^CalcR∩VGAT^ cells expressing scrambled shRNA or CalcR shRNA (scrambled shRNA: n = 15, 14 cells from 4 mice; CalcR shRNA: n = 16, 14 cells from 4 mice). Numbers in parentheses or axis titles indicate cell number. Circles represent data of individual cells. **(c)** Two-way ANOVA with repeated measures followed by Tukey’s multiple comparisons test. **(d, f, g, j, m)** Unpaired t-test. **(e, i, l)** Mann-Whitney test. Bars and error bars represent mean ± SEM. All p < 0.05 are indicated.

**Extended Data Fig. 15.**
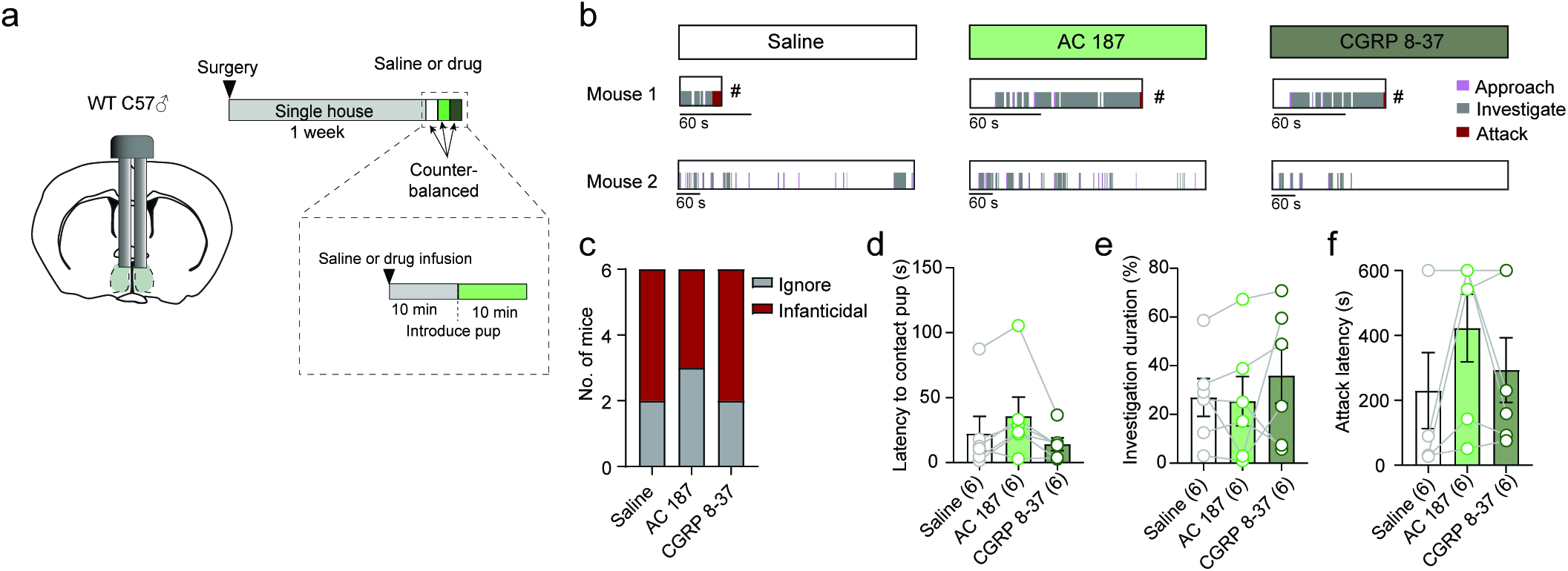
MPOA CalcR antagonism does not affect infanticide. **(a)** Cannula implant schematic and timeline to test behavior in non-pup sensitized mice following saline or drug infusion into the MPOA. **(b)** Representative raster plots showing pup-directed behaviors following saline, AC187, or CGRP 8-37 infusion. **(c)** The number of mice that were infanticidal or non-infanticidal and non-parental (ignore) from animals injected with saline, AC187, or CGRP 8-37. **(d-f)** The latency to contact pups **(d)**, percentage of time spent on investigating **(e)**, and latency to attack pups **(f)** following saline, AC187, or CGRP 8-37 infusion. Numbers in parentheses indicate the subject animal number. **(c)** Fisher’s exact test. **(d, f)** Friedman test followed by Dunn’s multiple comparisons test. **(e)** One-way ANOVA with repeated measures followed by Tukey’s multiple comparisons test. Lines and error bars represent mean ± SEM. All p < 0.05 are indicated.

**Extended Data Fig. 16.**
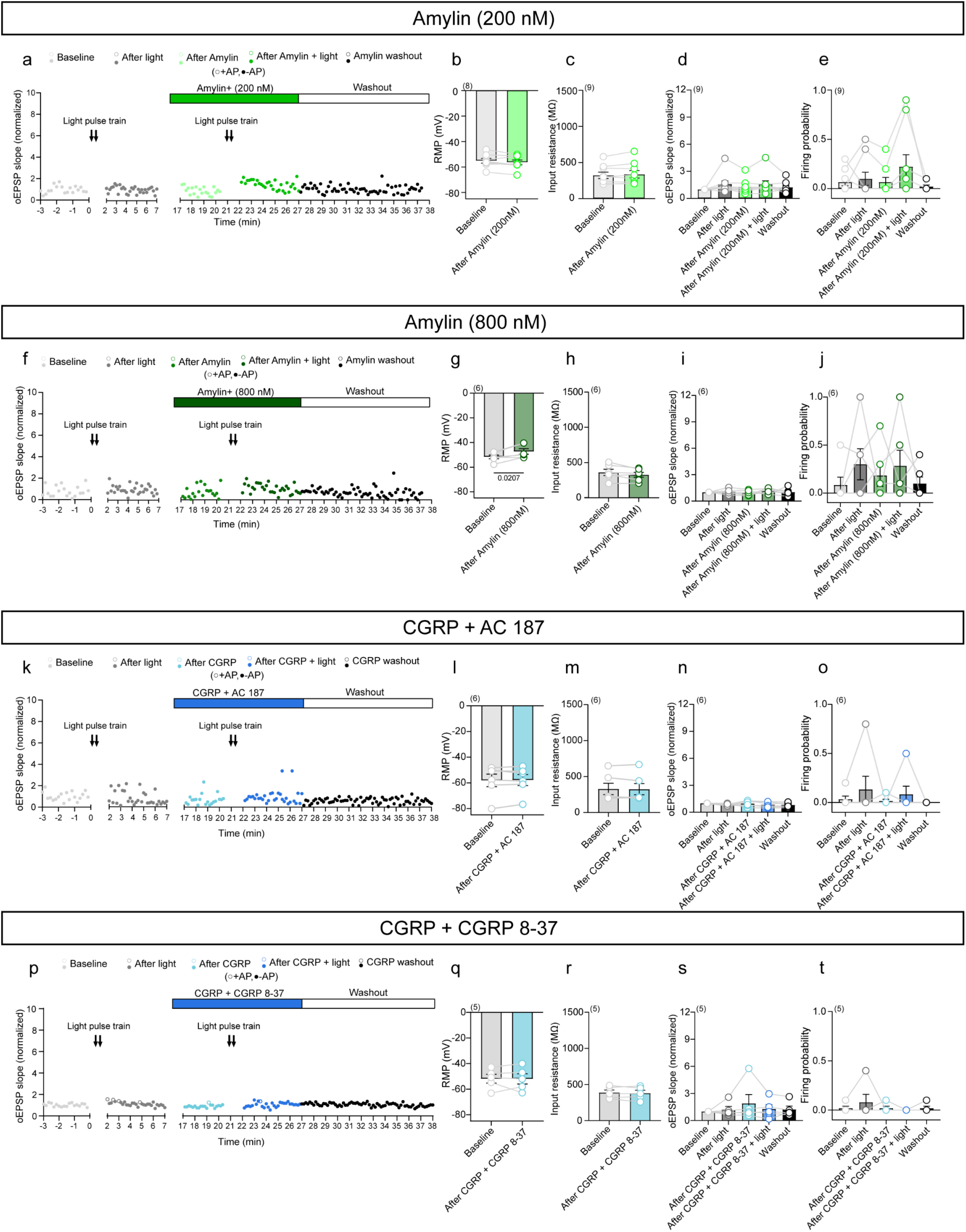
CGRP facilitates PA-MPOA^CalcR∩VGAT^ pathway potentiation through Amylin receptor. **(a)** Slopes of oEPSPs of an example MPOA^CalcR∩VGAT^ cell during baseline (light gray), after light stimulation (dark gray), after 200 nM amylin perfusion (light green), after light stimulation in the presence of 200 nM amylin (green), and during washout (black). Green bar represents the period of 200 nM amylin perfusion. White bar with black outlines represents the washout period. Open circles represent oEPSPs with action potentials (AP) and closed circles represent those without AP. **(b, c)** Resting membrane potential **(b)** and input resistance **(c)** before and after 200 nM amylin perfusion (n = 8, 9 cells from 4 mice). **(d, e)** Normalized oEPSP slope **(d)** and light stimulation-evoked firing probability **(e)** at baseline, after light stimulation without 200 nM amylin, after 200 nM amylin perfusion, after light stimulation in the presence of 200 nM amylin, and during washout (n = 9 cells from 4 mice). **(f-j)** The PA terminal activation-evoked responses of MPOA^CalcR∩VGAT^ cells after 800 nM Amylin and PA terminal stimulation. Plots follow the convention of **a-e**. (n = 6 cells from 3 mice). **(k-o)** The PA terminal activation-evoked responses of MPOA^CalcR∩VGAT^ cells after 200 nM CGRP and 600 nM AC187, an amylin receptor antagonist. Plots follow the convention of **a-e**. (n = 6 cells from 3 mice). **(p-t)** The PA terminal activation-evoked responses of MPOA^CalcR∩VGAT^ cells after 200 nM CGRP and 600 nM CGRP 8-37, an Amy1 and CGRP receptor dual antagonist. Plots follow the convention of **a-e**. (n = 5 cells from 3 mice). Numbers in parentheses indicate the number of cells. Circle-line pairs represent data of individual cells. **(b, c, g, h, l, q, r)** Paired t-test. **(m)** Wilcoxon test. **(d, e, i, j, n, o, s)** Friedman test followed by Dunn’s multiple comparisons test. Bars and error bars represent mean ± SEM. All p < 0.05 are indicated.

**Extended Data Fig. 17.**
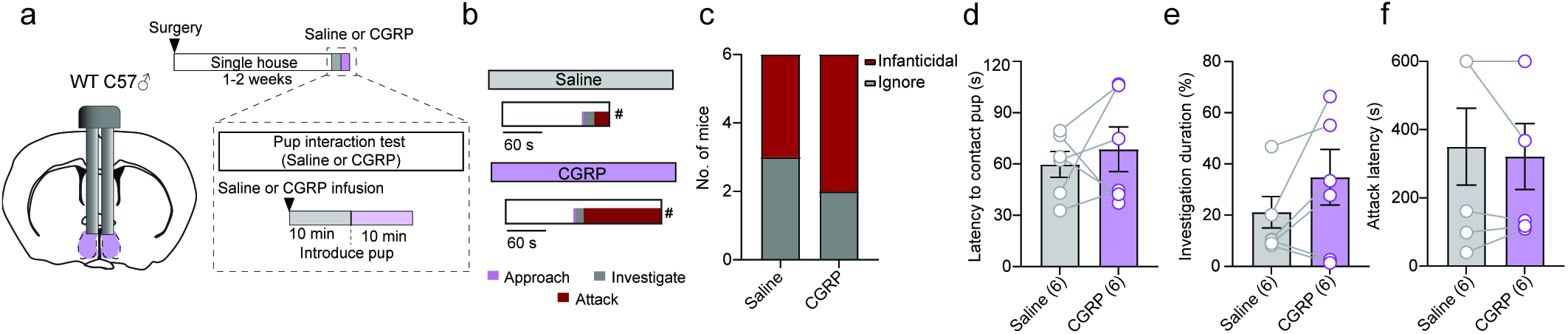
Acute CGRP infusion into the MPOA does not alter pup-directed behaviors in naïve males. **(a)** Cannula implant schematic and timeline to test behavior in non-pup sensitized animals after saline or CGRP infusion. **(b)** Representative raster plots showing pup-directed behaviors following saline or CGRP infusion into the MPOA. **(c)** The number of mice that were infanticidal or non-infanticidal and non-parental (ignore) during the pup interaction test. **(d-f)** The latency to contact pups **(d)**, investigation duration **(e)**, and latency to attack pups **(f)** following saline or CGRP infusion. If no attack occurred, the latency was scored as 600 s. Numbers in parentheses indicate the subject animal number. Circle-line pairs represent data of individual animals. **(c)** Fisher’s exact test. **(d, f)** Paired t-test. **(e)** Wilcoxon test. Bars and error bars represent mean ± SEM. All p < 0.05 are indicated.

## Notes

### Competing Interest Statement

The authors have declared no competing interest.

### Funding

- Eunice Kennedy Shriver National Institute of Child Health and Human Development: R01HD116127, R01HD092596, R01HD088411
- National Institute of Neurological Disorders and Stroke: U19NS107616
- U.S. National Science Foundation: Graduate Research Fellowship
- Brain Research Foundation
- National Institute of Mental Health: T32MH096331
- Brain & Behavior Research Foundation: Young Investigator Grant

## References

1 Gandelman, R. The ontogeny of maternal responsiveness in female Rockland-Swiss albino mice. Horm Behav 4, 257–268 (1973). 10.1016/0018-506x(73)90010-x

2 Carcea, I. et al. Oxytocin neurons enable social transmission of maternal behaviour. Nature 596, 553–557 (2021). 10.1038/s41586-021-03814-7

3 Marlin, B. J., Mitre, M., D’Amour J, A., Chao, M. V. & Froemke, R. C. Oxytocin enables maternal behaviour by balancing cortical inhibition. Nature 520, 499–504 (2015). 10.1038/nature14402

4 Fleming, A. S. & Rosenblatt, J. S. Maternal behavior in the virgin and lactating rat. J Comp Physiol Psychol 86, 957–972 (1974). 10.1037/h0036414

5 Rosenblatt, J. S. Nonhormonal basis of maternal behavior in the rat. Science 156, 1512– 1514 (1967). 10.1126/science.156.3781.1512

6 Stolzenberg, D. S. & Mayer, H. S. Experience-dependent mechanisms in the regulation of parental care. Front Neuroendocrinol 54, 100745 (2019). 10.1016/j.yfrne.2019.04.002

7 Stolzenberg, D. S. & Rissman, E. F. Oestrogen-independent, experience-induced maternal behaviour in female mice. J Neuroendocrinol 23, 345–354 (2011). 10.1111/j.1365-2826.2011.02112.x

8 vom Saal, F. S. & Howard, L. S. The regulation of infanticide and parental behavior: implications for reproductive success in male mice. Science 215, 1270–1272 (1982). 10.1126/science.7058349

9 Kennedy, H. F. & Elwood, R. W. Strain differences in the inhibition of infanticide in male mice (Mus musculus). Behav Neural Biol 50, 349–353 (1988). 10.1016/s0163-1047(88)91066-7

10 Tachikawa, K. S., Yoshihara, Y. & Kuroda, K. O. Behavioral transition from attack to parenting in male mice: a crucial role of the vomeronasal system. J Neurosci 33, 5120– 5126 (2013). 10.1523/JNEUROSCI.2364-12.2013

11 Elwood, R. W. Inhibition of infanticide and onset of paternal care in male mice (*Mus musculus*). J Comp Psych 99, 457 (1985).

12 Okabe, S., Kitano, K., Nagasawa, M., Mogi, K. & Kikusui, T. Testosterone inhibits facilitating effects of parenting experience on parental behavior and the oxytocin neural system in mice. Physiol Behav 118, 159–164 (2013). 10.1016/j.physbeh.2013.05.017

13 McCarthy, M. M. Short-term early exposure to pups alters infanticide in adulthood in male but not in female wild house mice (Mus domesticus). J Comp Psychol 104, 195– 197 (1990). 10.1037/0735-7036.104.2.195

14 Mitre, M. et al. Sex-Specific Differences in Oxytocin Receptor Expression and Function for Parental Behavior. Gend Genome 1, 142–166 (2017). 10.1089/gg.2017.0017

15 M, A.-L., B, L. & De, O. Repeated non-contact exposure to pups inhibits infanticidal and facilitates paternal behavior in virgin adult male mice (C57BL6). Physiology & Behavior 295, 114885 (2025). 10.1016/j.physbeh.2025.114885

16 Kenkel, W. M., Perkeybile, A. M. & Carter, C. S. The neurobiological causes and effects of alloparenting. Dev Neurobiol 77, 214–232 (2017). 10.1002/dneu.22465

17 Feldman, R. The Neurobiology of Human Attachments. Trends Cogn Sci 21, 80–99 (2017). 10.1016/j.tics.2016.11.007

18 Numan, M. Medial preoptic area and maternal behavior in the female rat. J Comp Physiol Psychol 87, 746–759 (1974). 10.1037/h0036974

19 Dulac, C., O’Connell, L. A. & Wu, Z. Neural control of maternal and paternal behaviors. Science 345, 765–770 (2014). 10.1126/science.1253291

20 O’Neill, P. T. & Lin, D. Neural plasticity supporting parental behaviors. Current Opinion in Neurobiology 95, 103111 (2025). 10.1016/j.conb.2025.103111

21 Numan, M. Neural basis of maternal behavior in the rat. Psychoneuroendocrinology 13, 47–62 (1988). 10.1016/0306-4530(88)90006-6

22 Numan, M. & Callahan, E. C. The connections of the medial preoptic region and maternal behavior in the rat. Physiol Behav 25, 653–665 (1980). 10.1016/0031-9384(80)90367-4

23 Numan, M., Corodimas, K. P., Numan, M. J., Factor, E. M. & Piers, W. D. Axon-sparing lesions of the preoptic region and substantia innominata disrupt maternal behavior in rats. Behav Neurosci 102, 381–396 (1988). 10.1037//0735-7044.102.3.381

24 Terkel, J., Bridges, R. S. & Sawyer, C. H. Effects of transecting lateral neural connections of the medial preoptic area on maternal behavior in the rat: nest building, pup retrieval and prolactin secretion. Brain Res 169, 369–380 (1979). 10.1016/0006-8993(79)91037-0

25 Tsuneoka, Y. et al. Functional, anatomical, and neurochemical differentiation of medial preoptic area subregions in relation to maternal behavior in the mouse. J Comp Neurol 521, 1633–1663 (2013). 10.1002/cne.23251

26 Lee, A., Clancy, S. & Fleming, A. S. Mother rats bar-press for pups: effects of lesions of the mpoa and limbic sites on maternal behavior and operant responding for pup-reinforcement. Behav Brain Res 100, 15–31 (1999). 10.1016/s0166-4328(98)00109-0

27 Jacobson, C. D., Terkel, J., Gorski, R. A. & Sawyer, C. H. Effects of small medial preoptic area lesions on maternal behavior: retrieving and nest building in the rat. Brain Res 194, 471–478 (1980). 10.1016/0006-8993(80)91226-3

28 Sturgis, J. D. & Bridges, R. S. N-methyl-DL-aspartic acid lesions of the medial preoptic area disrupt ongoing parental behavior in male rats. Physiol Behav 62, 305–310 (1997). 10.1016/s0031-9384(97)88985-8

29 Tsuneoka, Y. et al. Distinct preoptic-BST nuclei dissociate paternal and infanticidal behavior in mice. EMBO J 34, 2652–2670 (2015). 10.15252/embj.201591942

30 Fang, Y. Y., Yamaguchi, T., Song, S. C., Tritsch, N. X. & Lin, D. A Hypothalamic Midbrain Pathway Essential for Driving Maternal Behaviors. Neuron 98, 192–207 e110 (2018). 10.1016/j.neuron.2018.02.019

31 Kohl, J. et al. Functional circuit architecture underlying parental behaviour. Nature 556, 326–331 (2018). 10.1038/s41586-018-0027-0

32 Wei, Y. C. et al. Medial preoptic area in mice is capable of mediating sexually dimorphic behaviors regardless of gender. Nat Commun 9, 279 (2018). 10.1038/s41467-017-02648-0

33 Smiley, K. O., Brown, R. S. E. & Grattan, D. R. Prolactin Action Is Necessary for Parental Behavior in Male Mice. J Neurosci 42, 8308–8327 (2022). 10.1523/jneurosci.0558-22.2022

34 Brown, R. S. E. et al. Prolactin action in the medial preoptic area is necessary for postpartum maternal nursing behavior. Proceedings of the National Academy of Sciences of the United States of America 114, 10779–10784 (2017). 10.1073/pnas.1708025114

35 Moffitt, J. R. et al. Molecular, spatial, and functional single-cell profiling of the hypothalamic preoptic region. Science 362 (2018). 10.1126/science.aau5324

36 Stagkourakis, S. et al. A Neuro-hormonal Circuit for Paternal Behavior Controlled by a Hypothalamic Network Oscillation. Cell 182, 960–975 e915 (2020). 10.1016/j.cell.2020.07.007

37 Yoshihara, C. et al. Calcitonin receptor signaling in the medial preoptic area enables risk-taking maternal care. Cell Rep 35, 109204 (2021). 10.1016/j.celrep.2021.109204

38 Wu, Z., Autry, A. E., Bergan, J. F., Watabe-Uchida, M. & Dulac, C. G. Galanin neurons in the medial preoptic area govern parental behaviour. Nature 509, 325–330 (2014). 10.1038/nature13307

39 Perrigo, G., Bryant, W. C. & vom Saal, F. S. Fetal, hormonal and experiential factors influencing the mating-induced regulation of infanticide in male house mice. Physiol Behav 46, 121–128 (1989). 10.1016/0031-9384(89)90244-8

40 Pan, W. et al. Essential Role for Hypothalamic Calcitonin Receptor‒Expressing Neurons in the Control of Food Intake by Leptin. Endocrinology 159, 1860–1872 (2018). 10.1210/en.2017-03259

41 Knoedler, J. R. et al. A functional cellular framework for sex and estrous cycle-dependent gene expression and behavior. Cell 185, 654–671.e622 (2022). 10.1016/j.cell.2021.12.031

42 Zhang, G. W. et al. Medial preoptic area antagonistically mediates stress-induced anxiety and parental behavior. Nat Neurosci 24, 516–528 (2021). 10.1038/s41593-020-00784-3

43 Fenno, L. E. et al. Comprehensive Dual- and Triple-Feature Intersectional Single-Vector Delivery of Diverse Functional Payloads to Cells of Behaving Mammals. Neuron 107, 836–853 e811 (2020). 10.1016/j.neuron.2020.06.003

44 Daigle, T. L. et al. A Suite of Transgenic Driver and Reporter Mouse Lines with Enhanced Brain-Cell-Type Targeting and Functionality. Cell 174, 465–480.e422 (2018). 10.1016/j.cell.2018.06.035

45 Scott, N., Prigge, M., Yizhar, O. & Kimchi, T. A sexually dimorphic hypothalamic circuit controls maternal care and oxytocin secretion. Nature 525, 519–522 (2015). 10.1038/nature15378

46 Mei, L., Yan, R., Yin, L., Sullivan, R. M. & Lin, D. Antagonistic circuits mediating infanticide and maternal care in female mice. Nature 618, 1006–1016 (2023). 10.1038/s41586-023-06147-9

47 Autry, A. E. et al. Urocortin-3 neurons in the mouse perifornical area promote infant-directed neglect and aggression. Elife 10 (2021). 10.7554/eLife.64680

48 Chen, T. W. et al. Ultrasensitive fluorescent proteins for imaging neuronal activity. Nature 499, 295–300 (2013). 10.1038/nature12354

49 Zong, W. et al. Miniature two-photon microscopy for enlarged field-of-view, multi-plane and long-term brain imaging. Nat Methods 18, 46–49 (2021). 10.1038/s41592-020-01024-z

50 Falkner, A. L., Grosenick, L., Davidson, T. J., Deisseroth, K. & Lin, D. Hypothalamic control of male aggression-seeking behavior. Nat Neurosci 19, 596–604 (2016). 10.1038/nn.4264

51 Fukumitsu, K., Yoshihara, C., Huang, A. J., McHugh, T. J. & Kuroda, K. O. In vivo recording from calcitonin receptor-expressing neurons in the medial preoptic area during affiliative social behaviors. Neurosci Res 216, 104878 (2025). 10.1016/j.neures.2025.01.007

52 Karigo, T. et al. Distinct hypothalamic control of same- and opposite-sex mounting behaviour in mice. Nature 589, 258–263 (2021). 10.1038/s41586-020-2995-0

53 Ammari, R. et al. Hormone-mediated neural remodeling orchestrates parenting onset during pregnancy. Science 382, 76–81 (2023). 10.1126/science.adi0576

54 Hay, D. L., Garelja, M. L., Poyner, D. R. & Walker, C. S. Update on the pharmacology of calcitonin/CGRP family of peptides: IUPHAR Review 25. Br J Pharmacol 175, 3–17 (2018). 10.1111/bph.14075

55 Dobolyi, A. Central amylin expression and its induction in rat dams. J Neurochem 111, 1490–1500 (2009). 10.1111/j.1471-4159.2009.06422.x

56 Lutz, T. A. Creating the amylin story. Appetite 172, 105965 (2022). 10.1016/j.appet.2022.105965

57 Garelja, M. L., Walker, C. S. & Hay, D. L. CGRP receptor antagonists for migraine. Are they also AMY(1) receptor antagonists? Br J Pharmacol 179, 454–459 (2022). 10.1111/bph.15585

58 Bailey, R. J. et al. Pharmacological characterization of rat amylin receptors: implications for the identification of amylin receptor subtypes. Br J Pharmacol 166, 151–167 (2012). 10.1111/j.1476-5381.2011.01717.x

59 Hay, D. L., Christopoulos, G., Christopoulos, A., Poyner, D. R. & Sexton, P. M. Pharmacological discrimination of calcitonin receptor: receptor activity-modifying protein complexes. Mol Pharmacol 67, 1655–1665 (2005). 10.1124/mol.104.008615

60 Morimoto, S. et al. Calcitonin in plasma and cerebrospinal fluid from normal subjects and patients with medullary thyroid carcinoma: possible restriction of calcitonin by the blood-brain barrier. J Clin Endocrinol Metab 55, 594–596 (1982). 10.1210/jcem-55-3-594

61 Wimalawansa, S. J. Amylin, calcitonin gene-related peptide, calcitonin, and adrenomedullin: a peptide superfamily. Crit Rev Neurobiol 11, 167–239 (1997). 10.1615/critrevneurobiol.v11.i2-3.40

62 Yamaguchi, T. et al. Posterior amygdala regulates sexual and aggressive behaviors in male mice. Nat Neurosci 23, 1111–1124 (2020). 10.1038/s41593-020-0675-x

63 Sato, K. et al. Distinct neuronal populations mediate parenting and infanticide in the amygdalohippocampal area. Research Square (2022). 10.21203/rs.3.rs-2261172/v1

64 Klapoetke, N. C. et al. Independent optical excitation of distinct neural populations. Nat Methods 11, 338–346 (2014). 10.1038/nmeth.2836

65 Mei, L., Osakada, T. & Lin, D. Hypothalamic control of innate social behaviors. Science 382, 399–404 (2023). 10.1126/science.adh8489

66 Remedios, R. et al. Social behaviour shapes hypothalamic neural ensemble representations of conspecific sex. Nature 550, 388–392 (2017). 10.1038/nature23885

67 Zhang, J. X., Rao, X. P., Sun, L., Zhao, C. H. & Qin, X. W. Putative chemical signals about sex, individuality, and genetic background in the preputial gland and urine of the house mouse (Mus musculus). Chem Senses 32, 293–303 (2007). 10.1093/chemse/bjl058

68 Walker, C. S. et al. A second trigeminal CGRP receptor: function and expression of the AMY1 receptor. Ann Clin Transl Neurol 2, 595–608 (2015). 10.1002/acn3.197

69 Li, Z., Kelly, L., Heiman, M., Greengard, P. & Friedman, J. M. Hypothalamic Amylin Acts in Concert with Leptin to Regulate Food Intake. Cell Metab 22, 1059–1067 (2015). 10.1016/j.cmet.2015.10.012

70 Dobolyi, A., Irwin, S., Makara, G., Usdin, T. B. & Palkovits, M. Calcitonin gene-related peptide-containing pathways in the rat forebrain. J Comp Neurol 489, 92–119 (2005). 10.1002/cne.20618

71 Lechner, J., Leah, J. D. & Zimmermann, M. Brainstem peptidergic neurons projecting to the medial and lateral thalamus and zona incerta in the rat. Brain Res 603, 47–56 (1993). 10.1016/0006-8993(93)91298-7

72 Lein, E. S. et al. Genome-wide atlas of gene expression in the adult mouse brain. Nature 445, 168–176 (2007). 10.1038/nature05453

73 Okabe, S. et al. Pup odor and ultrasonic vocalizations synergistically stimulate maternal attention in mice. Behav Neurosci 127, 432–438 (2013). 10.1037/a0032395

74 Valtcheva, S. et al. Neural circuitry for maternal oxytocin release induced by infant cries. Nature 621, 788–795 (2023). 10.1038/s41586-023-06540-4

75 Yang, M., Keller, D., Dobolyi, A. & Valtcheva, S. The lateral thalamus: a bridge between multisensory processing and naturalistic behaviors. Trends Neurosci 48, 33–46 (2025). 10.1016/j.tins.2024.11.005

76 Bayless, D. W. et al. Limbic Neurons Shape Sex Recognition and Social Behavior in Sexually Naive Males. Cell 176, 1190–1205 e1120 (2019). 10.1016/j.cell.2018.12.041

77 Chen, A. X. et al. Specific Hypothalamic Neurons Required for Sensing Conspecific Male Cues Relevant to Inter-male Aggression. Neuron 108, 763–774 e766 (2020). 10.1016/j.neuron.2020.08.025

78 Li, Y. et al. Neuronal Representation of Social Information in the Medial Amygdala of Awake Behaving Mice. Cell 171, 1176–1190 e1117 (2017). 10.1016/j.cell.2017.10.015

79 Ribeiro, A. C. et al. siRNA silencing of estrogen receptor-alpha expression specifically in medial preoptic area neurons abolishes maternal care in female mice. Proceedings of the National Academy of Sciences of the United States of America 109, 16324–16329 (2012). 10.1073/pnas.1214094109

80 Froemke, R. C. & Young, L. J. Oxytocin, Neural Plasticity, and Social Behavior. Annu Rev Neurosci 44, 359–381 (2021). 10.1146/annurev-neuro-102320-102847

81 Inada, K. et al. Plasticity of neural connections underlying oxytocin-mediated parental behaviors of male mice. Neuron 110, 2009–2023 e2005 (2022). 10.1016/j.neuron.2022.03.033

82 Pedersen, C. A., Caldwell, J. D., Walker, C., Ayers, G. & Mason, G. A. Oxytocin activates the postpartum onset of rat maternal behavior in the ventral tegmental and medial preoptic areas. Behav Neurosci 108, 1163–1171 (1994). 10.1037//0735-7044.108.6.1163

83 Inada, K. et al. Vasopressin-to-oxytocin receptor crosstalk in the preoptic area underlying parental behaviors in male mice. Nat Commun 16, 10844 (2025). 10.1038/s41467-025-66908-0

84 Fukumitsu, K. et al. Amylin-Calcitonin receptor signaling in the medial preoptic area mediates affiliative social behaviors in female mice. Nat Commun 13, 709 (2022). 10.1038/s41467-022-28131-z

85 Sun, F. et al. Shared neural substrates of prosocial and parenting behaviours. Nature (2026). 10.1038/s41586-026-10327-8

86 Wu, Y. E. et al. Neural control of affiliative touch in prosocial interaction. Nature 599, 262–267 (2021). 10.1038/s41586-021-03962-w

87 Hu, R. K. et al. An amygdala-to-hypothalamus circuit for social reward. Nat Neurosci 24, 831–842 (2021). 10.1038/s41593-021-00828-2

88 Liu, D. et al. A hypothalamic circuit underlying the dynamic control of social homeostasis. Nature 640, 1000–1010 (2025). 10.1038/s41586-025-08617-8

89 Osborne, J. E. & Dudman, J. T. RIVETS: a mechanical system for in vivo and in vitro electrophysiology and imaging. PLoS One 9, e89007 (2014). 10.1371/journal.pone.0089007

90 Lin, D. et al. Functional identification of an aggression locus in the mouse hypothalamus. Nature 470, 221–226 (2011). 10.1038/nature09736

91 Franklin, K. B. J., Paxinos, G. The mouse brain in stereotaxic coordinates. Third Edition edn, (Elsevier, 2007).

92 Bankhead, P. et al. QuPath: Open source software for digital pathology image analysis. Sci Rep 7, 16878 (2017). 10.1038/s41598-017-17204-5

93 Zong, W. et al. Fast high-resolution miniature two-photon microscopy for brain imaging in freely behaving mice. Nat Methods 14, 713–719 (2017). 10.1038/nmeth.4305

94 McHenry, J. A. et al. Corrigendum: Hormonal gain control of a medial preoptic area social reward circuit. Nat Neurosci 20, 1427–1430 (2017). 10.1038/nn1017-1427c

95 Zhan, J., Komal, R., Keenan, W. T., Hattar, S. & Fernandez, D. C. Non-invasive Strategies for Chronic Manipulation of DREADD-controlled Neuronal Activity. J Vis Exp (2019). 10.3791/59439

